# Feed-forward regulation adaptively evolves via dynamics rather than topology when there is intrinsic noise

**DOI:** 10.1101/393884

**Authors:** Kun Xiong, Alex K. Lancaster, Mark L. Siegal, Joanna Masel

## Abstract

We develop a null model of the evolution of transcriptional regulatory networks, and use it to support an adaptive origin for a canonical “motif”, a 3-node feed-forward loop (FFL) hypothesized to filter out short spurious signals by integrating information from a fast and a slow pathway. Our mutational model captures the intrinsically high prevalence of weak affinity transcription factor binding sites. We also capture stochasticity and delays in gene expression that distort external signals and intrinsically generate noise. Functional FFLs evolve readily under selection for the hypothesized function, but not in negative controls. Interestingly, a 4-node “diamond” motif also emerged as a short spurious signal filter. The diamond uses expression dynamics rather than path length to provide fast and slow pathways. When there is no external spurious signal to filter out, but only internally generated noise, only the diamond and not the FFL evolves.

## Introduction

Transcriptional regulatory networks (TRNs) are integral to development and physiology, and underlie all complex traits. An intriguing finding about TRNs is that certain topological “motifs” of interconnected transcription factors (TFs) are over-represented relative to random re-wirings that preserve the frequency distribution of connections. The significance of this finding remains open to debate.

The canonical example is the feed-forward loop (FFL), in which TF A regulates a target C both directly, and indirectly via TF B, and no regulatory connections exist in the opposite direction^1-3^. Each of the three regulatory interactions in a FFL can be either activating or repressing, so there are eight distinct kinds of FFLs (**Fig. S1**)^4^. Given the eight frequencies expected from the ratio of activators to repressors, two of these kinds of FFLs are significantly over-represented^4^. In this paper, we focus on one of these two over-represented types, namely the type 1 coherent FFL (C1-FFL), in which all three links are activating rather than repressing (**Fig. S1**, top left). C1-FFL motifs are an active part of systems biology research today, e.g. they are used to infer the function of specific regulatory pathways^5, 6^.

The over-representation of FFLs in observed TRNs is normally explained in terms of selection favoring a function of FFLs. Specifically, the most common adaptive hypothesis is that cells often benefit from ignoring short-lived signals and responding only to durable signals^3, 4, 7^. Evidence that C1-FFLs can perform this function comes from the behavior both of theoretical models^4^ and of *in vivo* gene circuits^7^. A C1-FFL can achieve this function when its regulatory logic is that of an “AND” gate, i.e. both the direct path from A to C and the indirect path from A to B to C must be activated before the response is triggered. In this case, the response will only be triggered if, by the time the signal trickles through the longer path, it is still active on the shorter path as well. This yields a response to long-lived signals but not short-lived signals.

However, just because a behavior is observed, we cannot conclude that the behavior is a historical consequence of past selection favoring that behavior^8, 9^. The explanatory power of this adaptive hypothesis of filtering out short-lived and spurious signals needs to be compared to that of alternative, non-adaptive hypotheses^10^. The over-representation of C1-FFLs might be a byproduct of some other behavior that was the true target of selection^11^. Alternatively, it might be an intrinsic property of TRNs generated by mutational processes – gene duplication patterns have been found to enrich for FFLs in general^12^, although not yet C1-FFLs in particular. Adaptationist claims about TRN organization have been accused of being just-so stories, with adaptive hypotheses still in need of testing against an appropriate null model of network evolution^13-23^.

Here we develop such a computational null model of TRN evolution, and apply it to the case of C1-FFL over-representation. We include sufficient realism in our model of cis-regulatory evolution to capture the non-adaptive effects of mutation in shaping TRNs. In particular, we consider “weak” TF binding sites (TFBSs) that can easily appear *de novo* by chance alone, and from there be selected to bind a TF more strongly, as well as simulating mutations that duplicate and delete genes.

We also capture the stochasticity of gene expression, which causes the number of mRNAs and hence proteins to fluctuate^24, 25^. This is important, because demand for spurious signal filtering and hence C1-FFL function may arise not just from external signals, but also from internal fluctuations. Stochasticity in gene expression also shapes how external spurious signals are propagated. Stochasticity is a constraint on what TRNs can achieve, but it can also be adaptively co-opted in evolution^26^; either way, it might underlie the evolution of certain motifs. Most other computational models of TRN evolution that consider gene expression as the major phenotype do not simulate stochasticity in gene expression (but see three notable exceptions^27-29^).

Here we ask whether AND-gated C1-FFLs evolve as a response to selection for filtering out short and spurious external signals. Our new model allows us to compare the frequencies of network motifs arising in the presence of this hypothesized evolutionary cause to motif frequencies arising under non-adaptive control simulations, i.e. evolution under conditions that lack short spurious external signals while controlling both for mutational biases and for less specific forms of selection. We also ask whether other network motifs evolve to filter out short spurious signals, and if so, whether different conditions favor the appearance of different motifs during evolution.

## Model overview

We simulate the dynamics of TRNs as the TFs activate and repress one another’s transcription over developmental time, to generate gene expression phenotypes on which selection then acts over longer evolutionary timescales. For each moment in developmental time, we simulate the numbers of nuclear and cytoplasmic mRNAs in a cell, the protein concentrations, and the chromatin state of each gene in a haploid genome. Transitions between three possible chromatin states -- Repressed, Intermediate, and Active -- are a stochastic function of TF binding, and transcription initiation from the Active state is also stochastic. An overview of the model is shown in **Fig. 1**. The pattern of TF binding affects chromatin, which affects transcription rates, eventually feeding back to affect the concentration of TFs and hence their binding. The genotype is specified by a set of cis-regulatory sequences that contain TFBSs to which TFs may bind, by which consensus sequence each TF recognizes and with what affinity, and by 5 gene-specific parameters that control gene expression as a function of TF binding: mean duration of transcriptional bursts, mRNA degradation, protein production, and protein degradation rates, and gene length (which affects delays in transcription and translation). An external signal (**Fig. 1A red**) is treated like another TF, and the concentration of an effector gene (**Fig. 1A blue**) in response is a primary determinant of fitness, combined with a cost associated with gene expression (**Fig. 1B**). Mutants replace resident genotypes as a function of the difference in estimated fitness (**Fig. 1C**). Parameter values, taken as far as possible from *Saccharomyces cerevisiae*, are summarized in **Table S1**. Source code in C is available at https://github.com/MaselLab/network-evolution-simulator.

**Figure 1.**
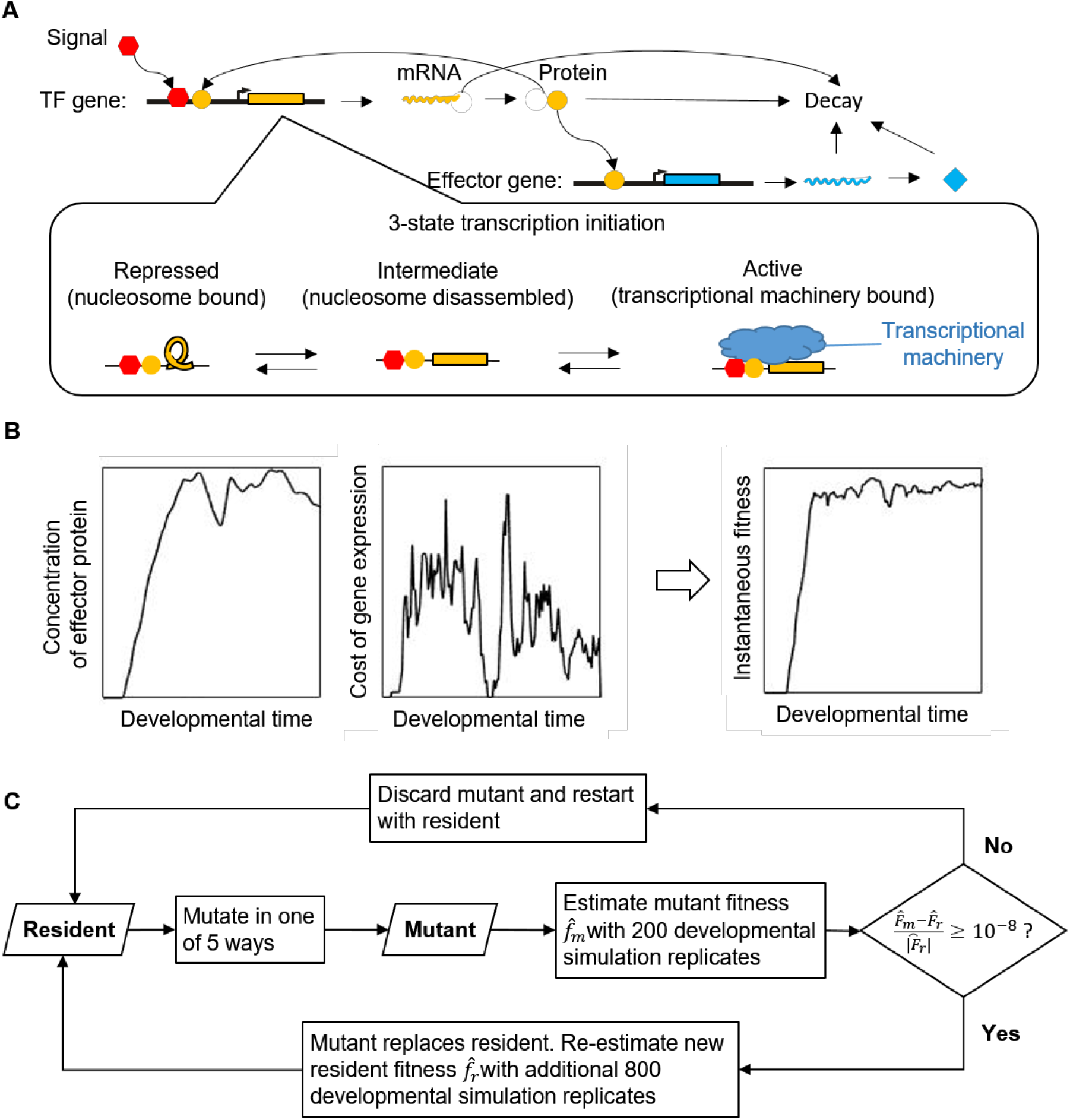
Overview of the model. **(A) Simulation of gene expression phenotypes.** We show a simple TRN with one TF (yellow) and one effector gene (blue), with arrows for major biological processes simulated in the model. **(B) Phenotype-fitness relationship.** Fitness is primarily determined by the concentration of an effector protein (here shown as beneficial as in Eq. 1, but potentially deleterious in a different environment as in Eq. 2), with a secondary component coming from the cost of gene expression (proportional to the rate of protein production), combined to give an instantaneous fitness at each moment in developmental time. **(C) Evolutionary simulation.** A single resident genotype is replaced when a mutant’s estimated fitness is high enough. Stochastic gene expression adds uncertainty to the estimated fitness, allowing less fit mutants to occasionally replace the resident, capturing the flavor of genetic drift.

### Transcription factor binding

Transcription of each gene is controlled by TFBSs present within a 150-bp cis-regulatory region. When bound, a TF occupies a stretch of DNA 14 bp long. In the center of this stretch, each TF recognizes an 8-bp consensus sequence, and binds to it with a TF-specific (and mutable) dissociation constant *K*_*d*_(0). TFs also bind somewhat specifically when there are one or two mismatches, with *K*_*d*_(1) and *K*_*d*_(2) values calculated from *K*_*d*_(0) according to a model of approximately additive binding energy per base pair. With three mismatches, binding occurs at the same background affinity as to any 14 bp stretch of DNA. We model competition between a smaller number of specific higher-affinity binding sites and the much larger number of non-specific binding sites, the latter corresponding to the total amount of nucleosome-free sequence in *S. cerevisiae*. Competition with non-specific binding can be approximated by using an effective dissociation constant 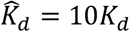. See Supplementary Text Section 1 for justification and details of these model choices.

Each TF is either an activator or a repressor. The algorithm for obtaining the probability distribution for *A* activators and *R* repressors being bound to a given cis-regulatory region at a given moment in developmental time is described in Supplementary Text Section 2.

### Transcriptional regulation

Activation of the effector gene requires at least two TFBSs to be occupied by activators – not necessarily different activators. The requirement for two activators makes the effector gene capable of evolving an AND-gate via a configuration of TFBSs in which the only way to have two TFs bound is for them to be different TFs (**Fig. 2**). All other genes are AND-gate-incapable, meaning that their activation requires only one TFBS to be occupied by an activator. *P*_*A*_ denotes the probability of having at least one activator bound for an AND-gate-incapable gene, or two for an AND-gate-capable gene. *P*_*R*_ denotes the probability of having at least one repressor bound.

**Figure 2.**
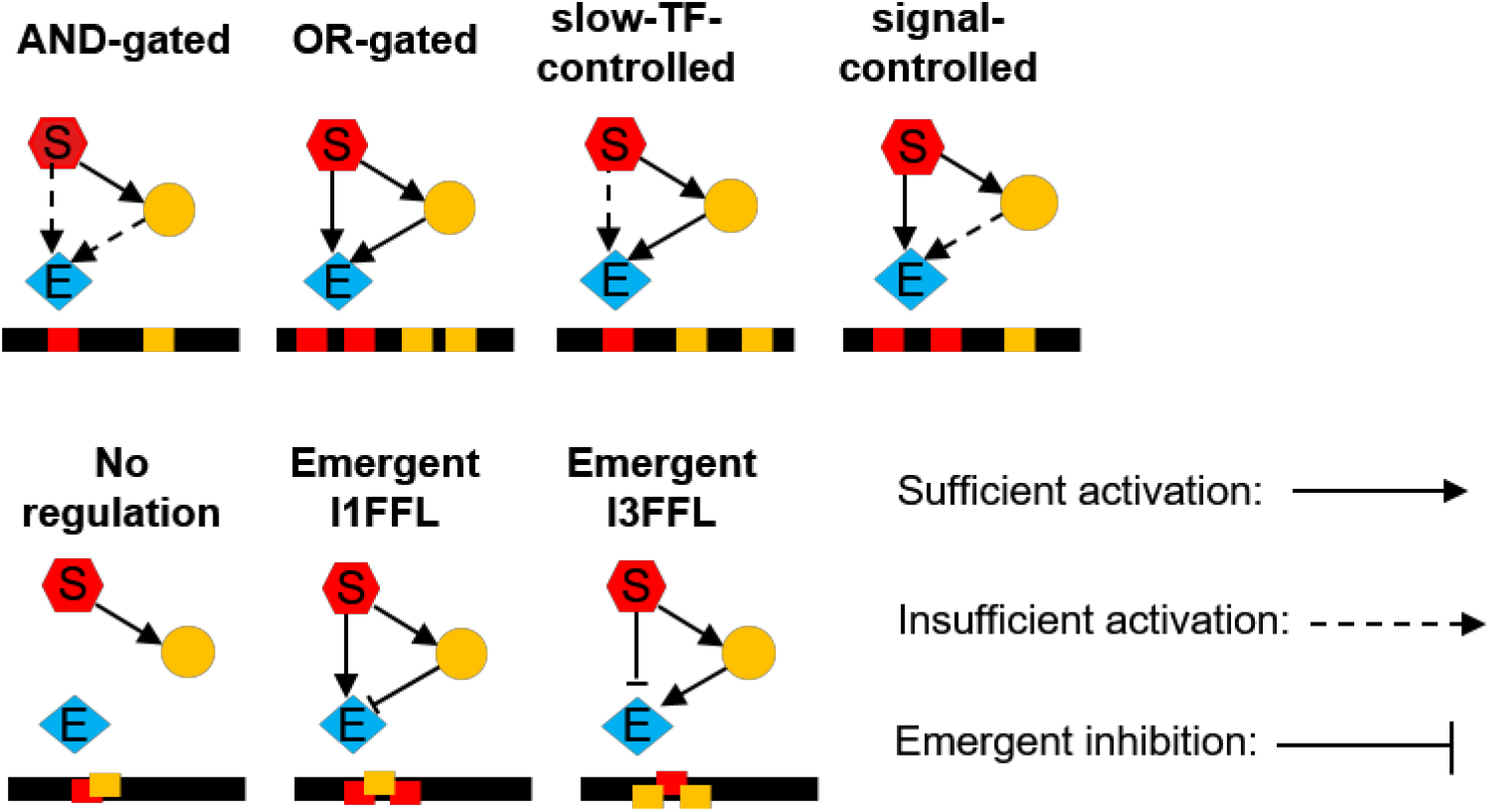
The numbers of TFBSs, and any hindrance between them, determine the regulatory logic of effector expression. We use the pattern of TFBSs (red and yellow bars along black cis-regulatory sequences) to classify the regulatory logic of the effector gene. C1-FFLs are classified first by whether or not they are capable of simultaneously binding the signal and the TF (top vs bottom). Further classification is based on whether either the signal or the TF has multiple non-overlapping TFBSs, allowing it to activate the effector without help from the other (solid arrow). The three subtypes on the bottom (where the signal and TF cannot bind simultaneously) are rarely seen; they are unless otherwise indicated included in “Any logic” and “non-AND-gated” tallies, but are not analyzed separately. Two of them involve emergent repression, creating “incoherent” feed-forward loops (see **Fig. S1** for full FFL naming scheme). Emergent repression occurs when the binding of one activator to its only TFBS prevents the other activator from binding to either of its two TFBSs, hence preventing simultaneous binding of two activators.

Noise in yeast gene expression is well described by a two step process of transcriptional activation^30, 31^, e.g. nucleosome disassembly followed by transcription machinery assembly. We denote the three corresponding possible states of the transcription start site as Repressed, Intermediate, and Active (**Fig. 1A**). Transitions between the states depend on the numbers of activator and repressor TFs bound (e.g. via recruitment of histone-modifying enzymes^32, 33^). We make conversion from Repressed to Intermediate a linear function of *P*_*A*_, ranging from the background rate 0.15 min^-1^ of histone acetylation^34^ (presumed to be followed by nucleosome disassembly), to the rate of nucleosome disassembly 0.92 min^-1^ for the constitutively active PHO5 promoter^30^:

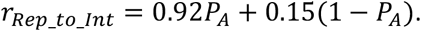

We make conversion from Intermediate to Repressed a linear function of *P*_*R*,_ ranging from a background histone de-acetylation rate of 0.67 min^-1 [34]^, up to a maximum of 4.11 min^-1^ (the latter chosen so as to keep a similar maximum:basal rate ratio as that of *r*_*Rep_to_Int*_):

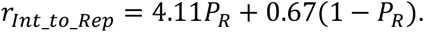

We assume that repressors disrupt the assembly of transcription machinery^35^ to such a degree that conversion from Intermediate to Active does not occur if even a single repressor is bound. In the absence of repressors, activators facilitate the assembly of transcription machinery^36^. Brown et al.^30^ reported that the rate of transcription machinery assembly is 3.3 min^-1^ for a constitutively active PHO5 promoter, and 0.025 min^-1^ when the Pho4 activator of the PHO5 promoter is knocked out. We use this range to set

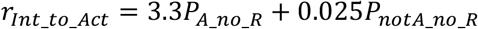

where *P*_*A_no_R*_ is the probability of having no repressors and either one (for an AND-gate-incapable gene) or two (for an AND-gate-capable gene) activators bound, and *P*_*notA_no_R*_ is the probability of having no TFs bound (for AND-gate-incapable genes) or having no repressors and not more than one activator bound (for AND-gate-capable genes).

The promoter sequence not only determines which specific TFBSs are present, but also influences non-specific components of the transcriptional machinery^37, 38^. We capture this via gene-specific but TF-binding-independent rates *r*_*Act_to_Int*_ with which the machinery disassembles and a burst of transcription ends. In other words, we let TF binding regulate the frequency of “bursts” of transcription, while other properties of the cis-regulatory region regulate their duration. For example, the yeast transcription factor Pho4 regulates the frequency but not duration of bursts of PHO5 expression, by regulating the rates of nucleosome removal and of transition to but not from a transcriptionally active state^30^. Parameterization of *r*_*Act_to_Int*_ is described in Supplementary Text Section 3.

### mRNA and protein dynamics

All genes in the Active state initiate new transcripts stochastically at rate *r*_*max_transc_init*_ = 6.75 mRNA/min^30^, while the time for completing transcription depends on gene length (see Supplementary Text Section 4 for parameterization of gene length and associated delay times). We model a second delay before a newly completed transcript produces the first protein, which we assume is dominated by translation initiation (length-independent) plus elongation (length-dependent) and not splicing or mRNA export (see Supplementary Text Section 5). After the second delay, we model protein production as continuous at a gene-specific rate *r*_*protein_syn*_ (see Supplementary Text Section 5).

Protein transport into the nucleus is rapid^39^ and is approximated as instantaneous and complete, so that the newly produced protein molecules immediately increase the probability of TF binding. Each gene has its own mRNA and protein decay rates, initialized from distributions taken from data (see Supplementary Text Section 6).

All the rates regarding transcription and translation are listed in **Table S1**, including distributions estimated from data, and hard bounds imposed to prevent unrealistic values arising during evolutionary simulations.

### Developmental simulation

Our algorithm is part stochastic, part deterministic. We use a Gillespie algorithm^40^ to simulate stochastic transitions between Repressed, Intermediate, and Active chromatin states, and to simulate transcription initiation and mRNA decay events. Fixed (i.e. deterministic) delay times are simulated between transcription initiation and completion, and between transcript completion and the production of the first protein. Protein production and degradation are described deterministically with ODEs, and updated frequently in order to recalculate TF concentrations and hence chromatin transition rates. Details of our simulation algorithm are given in the Supplementary Text Section 7. We initialize developmental simulations with no mRNA or protein, and all genes in the Repressed state.

### Selection

Filtering out short spurious signals is a special case of signal recognition. In environment 1, expressing the effector is beneficial, and in environment 2 it is deleterious. We select for TRNs that take information from the signal and correctly decide whether to express the effector. Fitness is a weighted average across separate developmental simulations in the two environments, one with a signal and one without. In both cases, we begin each developmental simulation with no signal. To ensure that gene expression changes in response to the signal, and not via an internal timer, we simulate a burn-in phase with duration drawn from an exponential distributed truncated at 30 minutes, with un-truncated mean of 10 minutes. By having no fitness effects of gene expression during the burn-in, we eliminate a significant source of noise in fitness estimation due to variable burn-in duration. In our control condition, at the end of the burn-in, the signal suddenly switches to a constant “on” level in environment 1, and remains off in environment 2. In our test condition (**Fig. 3**), the signal is turned on in the same way in environment 1 but is also briefly turned on (for the first 10 minutes after the burn-in) in environment 2 – selection is to ignore this short spurious signal. The signal is treated as though it were an activating TF whose concentration is controlled externally, with an “off” concentration of zero and an “on” concentration of 1,000 molecules per cell, which is the typical per-cell number of a yeast TF^41^.

**Figure 3.**
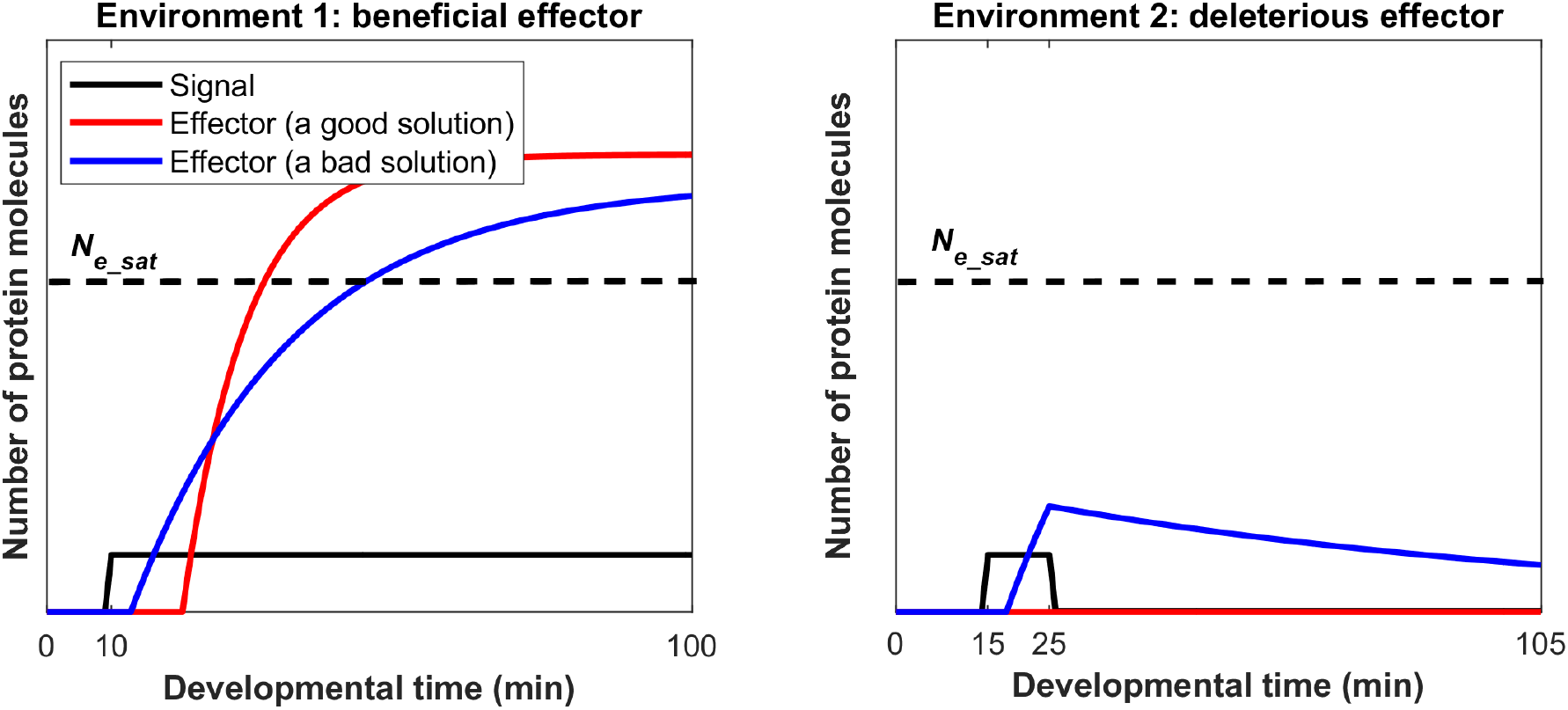
Selection for filtering out short spurious signals. Each selection condition averages fitness across simulations in two environments. The effectors have different fitness effects in the two environments, and the signal also behaves differently in the two environments. Simulations begin with zero mRNA and protein, and all genes at the Repressed state. Each simulation is burned in for a randomly sampled length of time in the absence of signal (shown here as 10 minutes in environment 1, and 15 minutes in environment 2), and continues for another 90 minutes after the burn-in. The signal is shown in black. Red illustrates a good solution in which the effector responds appropriately in each of the environments, while blue shows an inferior solution. See **Fig. S2** for examples of high-fitness and low-fitness evolved phenotypes, where, as shown in this schematic, high-fitness solutions have longer delays followed by more rapid responses thereafter.

We make fitness quantitative in terms of a “benefit” *B*(*t*) as a function of the amount of effector protein *N*_*e*_(*t*) at developmental time *t*. Our motivation is a scenario in which the effector protein is responsible for directing resources from a metabolic program favored in environment 2 to a metabolic program favored in environment 1. In environment 1, where the effector produces benefits,

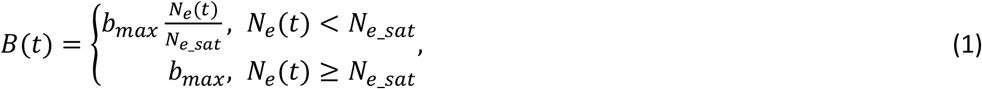

where *b*_*max*_ is the maximum benefit if all resources were redirected, and *N*_*e_sat*_ is the minimum amount of effector protein needed to achieve this. Similarly, in environment 2

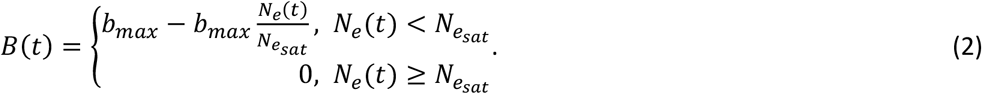

We set *N*_*e_sat*_ to 10,000 molecules, which is about the average number of molecules of a metabolism-associated protein per cell in yeast^41^. Without loss of generality given that fitness is relative, we set *b*_*max*_ to 1.

A second contribution to fitness comes from the cost of gene expression *C*(*t*) (**Fig. 1B, middle**). We make this cost proportional to the total protein production rate. We estimate a fitness cost of gene expression of 2×10^-6^ per protein molecule translated per minute, based on the cost of expressing a non-toxic protein in yeast^42^ (see Supplementary Text Section 7 for details).

We simulate gene expression for 90 minutes plus the duration of the burn-in (**Fig. 3**). A “cellular fitness” in a given environment is calculated as the average instantaneous fitness *B*(*t*)-*C*(*t*) over the 90 minutes. We consider environment 2 to be twice as common as environment 1 (a “signal” should be for an uncommon event rather than the default), and take the corresponding weighted average.

### Evolutionary simulation

We simulate a novel version of origin-fixation (weak-mutation-strong-selection) evolutionary dynamics, i.e. the population contains only one resident genotype at any time, and mutant genotypes are either rejected or chosen to be the next resident (Fig. 1C). Despite the fact that our mutant acceptance rule (see below) was chosen to maximize computational efficiency, our model usually takes 10 CPUs 1-3 days to complete an evolutionary simulation; modeling a heterogeneous population is clearly out of the question. We note that genetic homogeneity entails ignoring some important population genetic phenomena. First, if there were recombination, heterogeneity would favor mutations that combine well with a range of other genotypes. Second, clonal interference would shift evolution toward beneficial mutations of larger effect^43^ (an effect we can mimic by modifying the value 10^-8^ in the equation below). Third, polymorphic populations would evolve mutational robustness^44^. None of these three effects seems *a priori* likely to change our conclusions, although the possibility cannot be ruled out.

Estimators 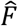 of genotype fitness are averages of the cellular fitness values of 200 developmental replicates per environment in the case of the mutant, plus an additional 800 should it be chosen to be the next resident. The mutant replaces the resident if

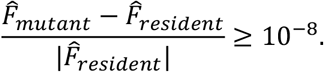

This differs from Kimura’s^45^ equation for fixation probability, but captures the flavor of genetic drift. Genetic drift allows slightly deleterious mutations to occasionally fix, and beneficial mutations to sometimes fail to do so, even as the probability of fixation is monotonic with fitness. This is also achieved by our procedure, because of stochastic deviations of 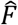 from true genotype fitness. The number of developmental replicates captures the flavor of effective population size.

Note that it is possible, especially at the beginning of an evolutionary simulation, for relative fitness to be paradoxically negative. This occurs when a randomly initialized genotype does not express the effector (garnering no fitness benefit), but does express other genes (accruing a cost of expression); this combination makes fitness negative. In this rare case, for simplicity, we use the absolute value of 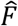 on the denominator.

If 2,000 successive mutants are all rejected, the simulation is terminated; upon inspection, we found that these resident genotypes had evolved to not express the effector in either environment. We refer to each change in resident genotype as an evolutionary step. We stop the simulation after 50,000 evolutionary steps; at this time, most replicate simulations seem to have reached a fitness plateau (**Fig. S3**); we analyze all replicates except those terminated early. To reduce the frequency of early termination in the case where the signal was not allowed to directly regulate the effector, we used a burn-in phase selecting on a more accessible intermediate phenotype (see Supplementary Text Section 10). In this case, burn-in occurred for 1,000 evolutionary steps, followed by the usual 50,000 evolutionary steps with selection for the phenotype of interest (**Fig. S3**, right panels). Most replicates found a stable fitness plateau within 10,000 evolutionary steps, although some replicates were temporarily trapped at a low fitness plateau (**Fig. S3**).

### Genotype Initialization

We initialize genotypes with 3 activator genes, 3 repressor genes, and 1 effector gene. Cis-regulatory sequences and consensus binding sequences contain As, Cs, Gs, and Ts sampled with equal probability. Rate constants associated with the expression of each gene are sampled from the distributions summarized in **Table S1**.

### Mutation

A genotype is subjected to 5 broad classes of mutation, at rates summarized in **Table S2** and justified in Supplementary Text Section 9. First are single nucleotide substitutions in the cis-regulatory sequence; the resident nucleotide mutates into one of the other three types of nucleotides with equal probability. Second are single nucleotide changes to the consensus binding sequence of a TF, with the resident nucleotide mutated into recognizing one of the other three types with equal probability. Both of these types of mutation can affect the number and strength of TFBSs.

Third are gene duplications or deletions. Because computational cost scales steeply (and non-linearly) with network size, we do not allow effector genes to duplicate once there are 5 copies, nor TF genes to duplicate once the total number of TF gene copies is 19. We also do not allow the signal, the last effector gene, nor the last TF gene to be deleted.

Fourth are mutations to gene-specific expression parameters. Most of these (*L, r*_*Act_to_Int*_, *r*_*protein_syn*_, *r*_*mRNA_deg*_, and *r*_*protein_deg*_) apply to both TFs and effector genes, while mutations to the gene-specific values of *K*_*d*_(0) apply only to TFs. Each mutation to *L* increases or decreases it by 1 codon, with equal probability unless *L* is at the upper or lower bound. Effect sizes of mutations to the other five parameters are modeled in such a way that mutation would maintain specified log-normal stationary distributions for these values, in the absence of selection or arbitrary bounds (see Supplementary Text Section 9 for details). Upper and lower bounds (Supplementary Text Section 9) are used to ensure that selection never drives these parameters to unrealistic values.

Fifth is conversion of a TF from being an activator to being a repressor, and vice versa. The signal is always an activator, and does not evolve.

Importantly, this scheme allows for divergence following gene duplication. When duplicates differ due only to mutations of class 4, i.e. protein function is unchanged, we refer to them as “copies” of the same gene, encoding “protein variants”. Mutations in classes 2 and 5 can create a new protein.

**Table S3** summarizes the tendencies of different mutation types to be accepted, and to contribute to evolution. Acceptance rates are high, indicative of substantial nearly neutral evolution, in which slightly deleterious mutations are fixed and subsequently compensated for.

## Results

### Functional AND-gated C1-FFLs evolve readily under selection for filtering out a short spurious signal

We begin by simulating the easiest case we can devise to allow the evolution of C1-FFLs for their purported function of filtering out short spurious signals. The signal is allowed to act directly on the effector, after which all that needs to evolve is a single activating TF between the two, as well as AND-logic for the effector (**Fig. 2**, top left; see “Transcriptional regulation” in the Model Overview for how AND-logic evolution is handled). We score network motifs at the end of a set period of evolution (see Supplemental Text Section 11 for details), further classifying evolved C1-FFLs into subtypes based on the presence of non-overlapping TFBSs (**Fig. 2**). The adaptive hypothesis predicts the evolution of the C1-FFL subtype with AND-regulatory logic, which requires the effector to be stimulated both by the signal and by the slow TF. While all evolutionary replicates show large increases in fitness, the extent of improvement varies dramatically, indicating whether or not the replicate was successful at evolving the phenotype of interest rather than becoming stuck at an alternative locally optimal phenotype (**Fig. 4A**). AND-gated C1-FFLs frequently evolve in replicates that reach high fitness outcomes, but not replicates that reach lower fitness (**Fig. 4B**).

**Figure 4.**
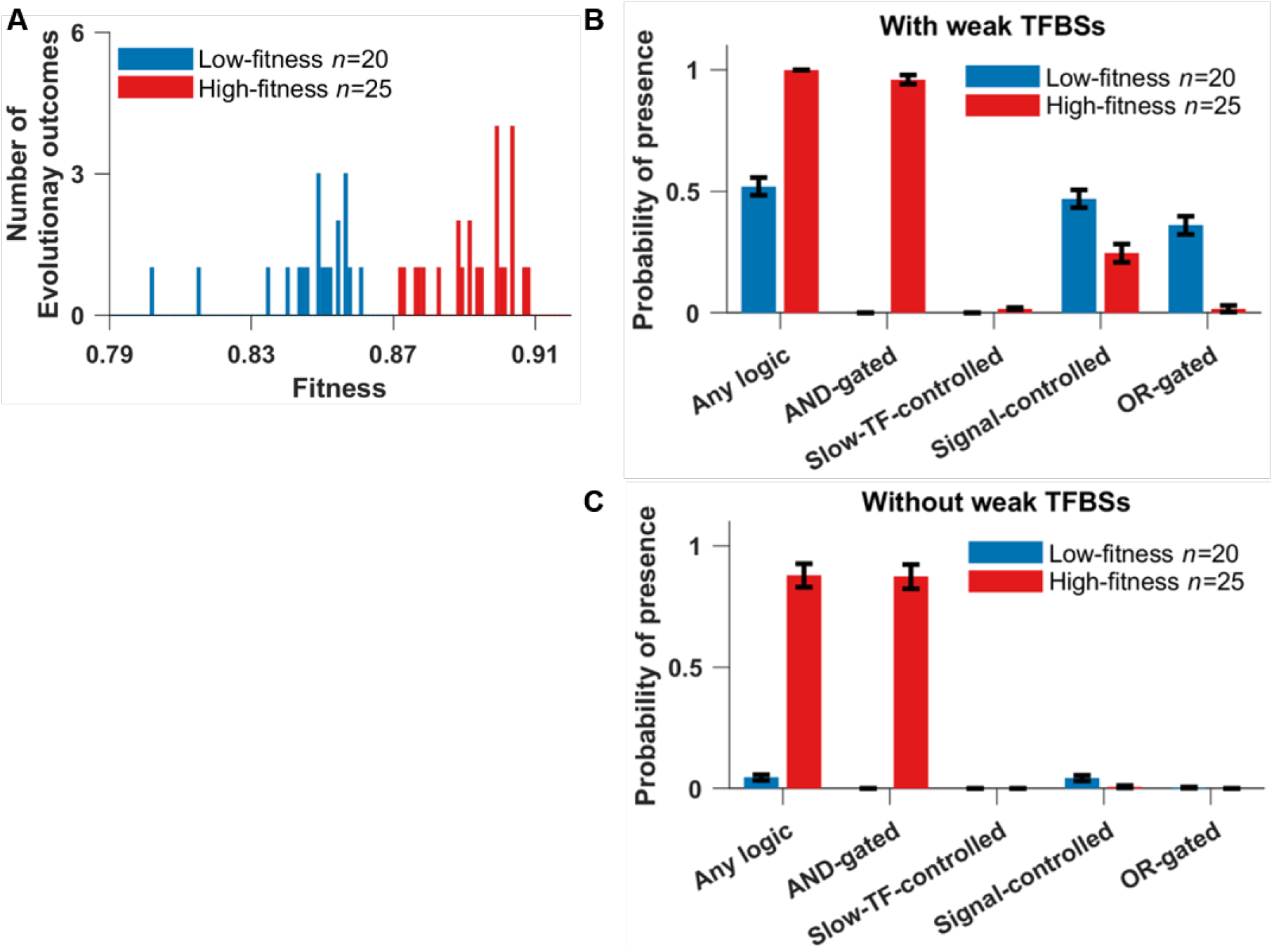
AND-gated C1-FFLs are associated with a successful response to selection for filtering out short spurious signals. **(A)** Distribution of fitness outcomes across replicate simulations, calculated as the average fitness over the last 10,000 steps of the evolutionary simulation. We divide genotypes into a low-fitness group (blue) and a high-fitness group (red) using as a threshold an observed gap in the distribution. **(B)** High fitness replicates are characterized by the presence of an AND-gated C1-FFL. “Any logic” counts the presence of any of the seven subtypes shown in **Fig. 2B**. Because one TRN can contain multiple C1-FFLs of different subtypes, each of which are scored, the sum of the occurrences of all seven subtypes will generally be more than “Any logic”. See Supplementary Text Section 11 for details on the calculation of the y-axis. **(C)** The over-representation of AND-gated C1-FFLs becomes even more pronounced relative to alternative logic-gating when weak (two-mismatch) TFBSs are excluded while scoring motifs. Data are shown as mean±SE of the occurrence over replicate evolution simulations.

We also see C1-FFLs that, contrary to expectations, are not AND-gated. Non-AND-gated motifs are found more often in low fitness than high fitness replicates (**Fig. 4B**), indicating that the preference for AND-gates is associated with adaptation rather than mutation bias. However, some non-AND-gated motifs are still found even in the high fitness replicates. This is because motifs and their logic gates are scored on the basis of all TFBSs, even those with two mismatches and hence low binding affinity. Unless these weak TFBSs are deleterious, they will appear quite often by chance alone. A random 8-bp sequence has probability 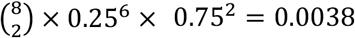 of being a two-mismatch binding site for a given TF. In our model, a TF has the potential to recognize 137 different sites in a 150-bp cis-regulatory sequence (taking into account steric hindrance at the edges), each with 2 orientations. Thus, by chance alone a given TF will have 0.0038 × 137 × 2 ≈ 1 two-mismatch binding sites in a given cis-regulatory sequence (ignoring palindromes for simplicity), compared to only ∼0.1 one-mismatch TFBSs. Non-AND-gated C1-FFLs mostly disappear when two-mismatch TFBSs are excluded, but the AND-gated C1-FFLs found in high fitness replicates do not (**Fig. 4C**).

To confirm the functionality of these AND-gated C1-FFLs, we mutated the evolved genotype in two different ways (**Fig. 5A**) to remove the AND regulatory logic. As expected, this lowers fitness in the presence of the short spurious signal but increases fitness in the presence of constant signal, with a net reduction in fitness (**Fig. 5B**). This is consistent with AND-gated C1-FFLs representing a tradeoff, by which a more rapid response to a true signal is sacrificed in favor of the greater reliability of filtering out short spurious signals.

**Figure 5.**
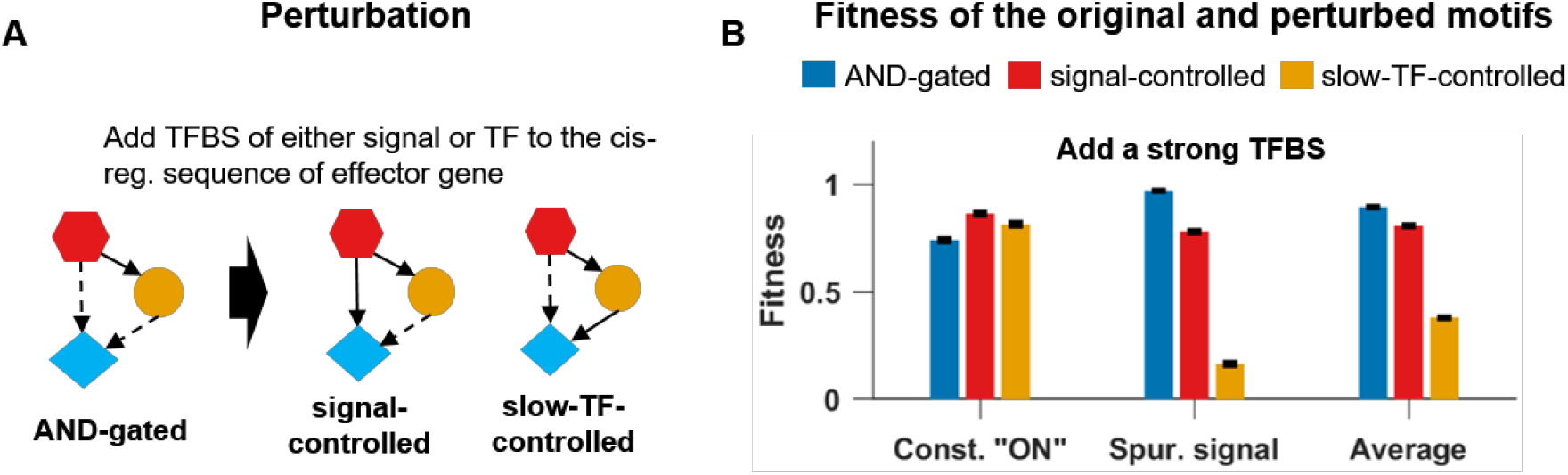
Destroying the AND-logic of a C1-FFL removes its ability to filter out short spurious signals. (**A**) For each of the *n* = 25 replicates in the high fitness group in **Fig. 4**, we perturbed the AND-logic in two ways, by adding one binding site of either the signal or the slow TF to the cis-regulatory sequence of the effector gene. **(B)** For each replicate, the fitness of the original motif (blue) or of the perturbed motif (red or orange) was averaged across the subset of evolutionary steps with an AND-gated C1-FFL and lacking other potentially confounding motifs (see Supplementary Text Section 11 for details). Destroying the AND-logic slightly increases the ability to respond to the signal, but leads to a larger loss of fitness when short spurious signals are responded to. Fitness is shown as mean±SE over replicate evolutionary simulations.

Adaptive motifs are constrained not only in their topology and regulatory logic, but also in the parameter space of their component genes. In particular, there is selection for rapid synthesis of both effector and TF proteins, as well as rapid degradation of effector mRNA and protein (**Table S4**). Fast effector degradation reduces the transient expression induced by the short spurious signal (**Fig. S2**).

To test the extent to which AND-gated C1-FFLs are a specific response to selection to filter out short spurious signals, we simulated evolution under three negative control conditions: 1) no selection, i.e. all mutations are accepted to become the new resident genotype; 2) no spurious signal, i.e. selection to express the effector under a constant “ON” signal and not under a constant “OFF” signal; 3) harmless spurious signal, i.e. selection to express the effector under a constant “ON” environment whereas effector expression in the “OFF” environment with short spurious signals is neither punished nor rewarded beyond the cost of unnecessary gene expression. AND-gated C1-FFLs evolve much less often under all three negative control conditions (**Fig. 6**), showing that their prevalence is a consequence of selection for filtering out short spurious signals, rather than a consequence of mutational bias and/or simpler forms of selection. C1-FFLs that do evolve under control conditions tend not to be AND-gated (**Fig. 6A**), and mostly disappear when weak TFBSs are excluded during motif scoring (**Fig. 6B**).

**Figure 6.**
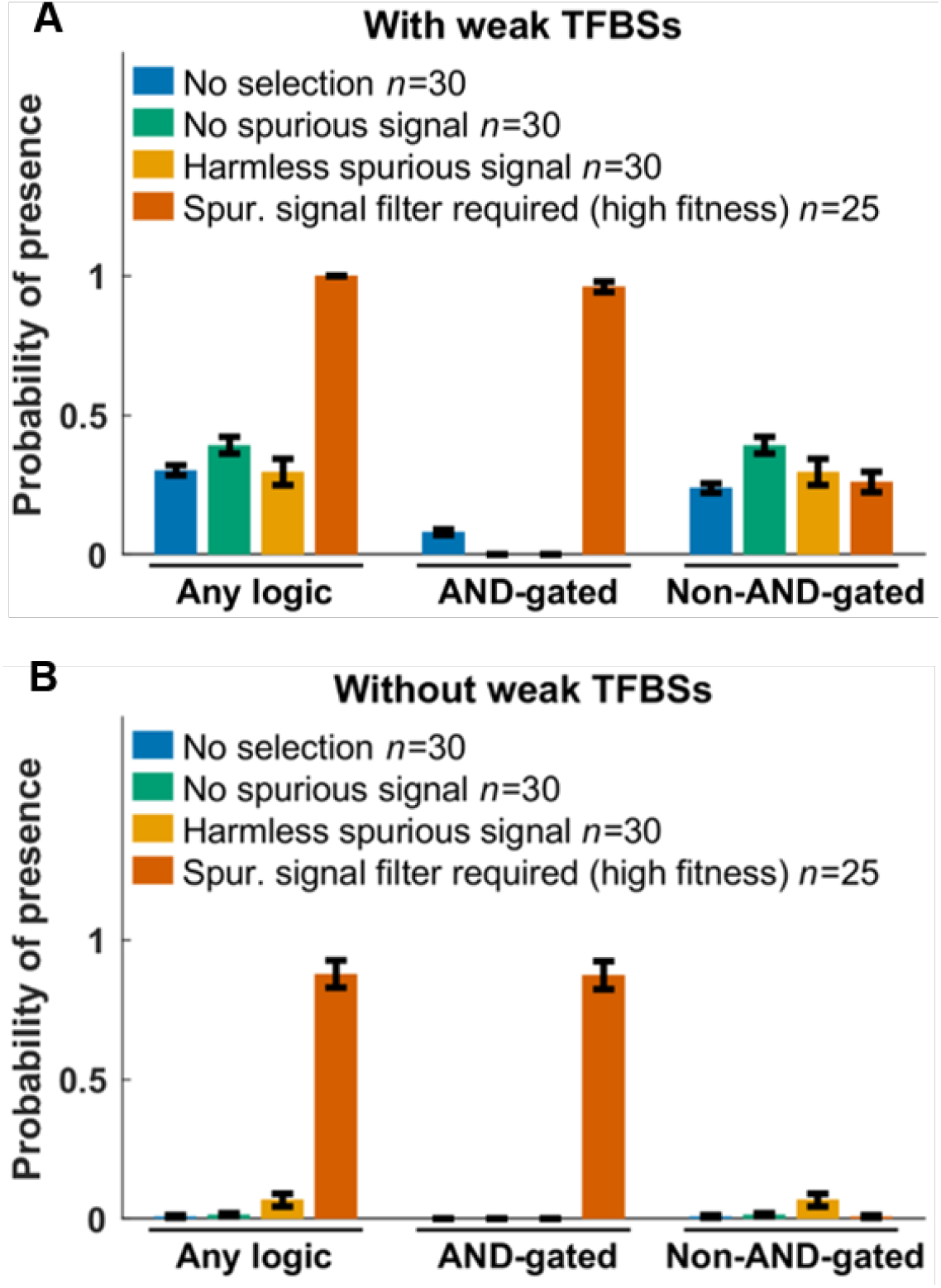
Selection for filtering out short spurious signals is the primary cause of C1-FFLs. TRNs are evolved under different selection conditions, and we score the probability that at least one C1-FFL is present (Supplementary Text Section 11). Weak (two-mismatch) TFBSs are included **(A)** or excluded **(B)** during motif scoring. Data are shown as mean±SE over evolutionary replicates. C1-FFL occurrence is similar for high-fitness and low-fitness outcomes in control selective conditions (**Fig. S4**), and so all evolutionary outcomes were combined. “Spurious signal filter required (high fitness)” uses the same data as in **Fig. 4**.

### Diamond motifs are an alternative adaptation in more complex networks

In real biological situations, sometimes the source signal will not be able to directly regulate an effector, and must instead operate via a longer regulatory pathway involving intermediate TFs^46^. In this case, even if the signal itself takes the idealized form shown in **Fig. 3**, its shape after propagation may become distorted by the intrinsic processes of transcription. Motifs are under selection to handle this distortion.

To enforce indirect regulation, we ran simulations in which the signal was only allowed to bind to the cis-regulatory sequences of TFs and not of effector genes. The fitness distribution of the evolutionary replicates has no obvious gaps (**Fig. S5**), so we compared the highest fitness, lowest fitness, and median fitness replicates. In agreement with results when direct regulation is allowed, genotypes of low and medium fitness contain few AND-gated C1-FFLs, while high fitness genotypes contain many more (**Fig. 7B, left and right**).

**Figure 7.**
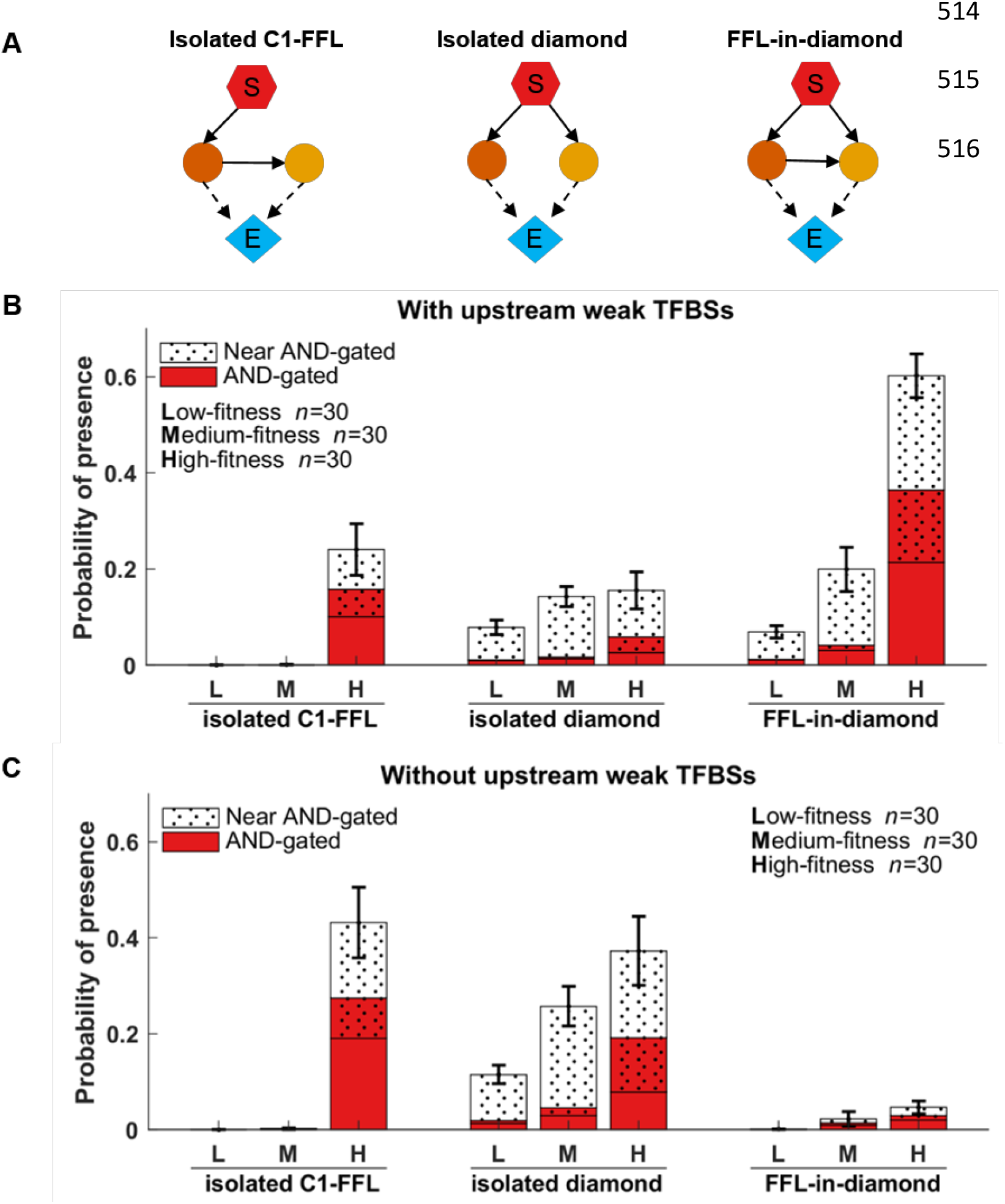
Both AND-gated C1-FFLs and AND-gated diamonds (A) are associated with high fitness in complex networks under selection to filter out short spurious signals. Out of 160 simulations (**Fig. S5**), we took the 30 with the highest fitness (H), the 30 with the lowest fitness (L), and 30 of around median fitness (M). AND-gated motifs are scored while including weak TFBSs in the effectors’ cis-regulatory regions, near-AND-gated motifs are those scored only when these weak TFBSs are excluded. It is possible for the same genotype to contain one of each, resulting in overlap between the red AND-gated columns and the dotted near-AND-gated columns. Weak TFBSs upstream in the TRN, i.e. not in the effector, are shown both included (**B**) and excluded (**C**). See Supplementary Text Section 11 for y-axis calculation details. Error bars show mean±SE of the proportion of evolutionary steps containing the motif in question, across replicate evolutionary simulations.

While visually examining the network context of these C1-FFLs, we discovered that many were embedded within AND-gated “diamonds”. In a diamond, the signal activates the expression of two genes that encode different TFs, and the two TFs activate the expression of an effector gene (**Fig. 7A middle**). When one of the two TF genes activates the other, then a C1-FFL is also present among the same set of genes; we call this topology a “FFL-in-diamond” (**Fig. 7A right**), and the prevalence of this configuration drew our attention toward diamonds. This led us to discover that AND-gated diamonds also occurred frequently without AND-gated C1-FFLs, in the configuration we call “isolated diamonds” (**Fig. 7A middle**). Note that it is in theory possible, but in practice uncommon, for diamonds to be part of more complex conjugates. Systematically scoring the AND-gated isolated diamond motif confirmed its high occurrence (**Fig. 7B and C, middle**).

An AND-gated C1-FFL integrates information from a short/fast regulatory pathway with information from a long/slow pathway, in order to filter out short spurious signals. A diamond achieves the same end of integrating fast and slowly transmitted information via differences in the gene expression dynamics of the two regulatory pathways, rather than via topological length (**Fig. 8**). The fast and slow pathways could be distinguished in a number of ways, e.g. by the slope at which the transcription factor concentration increases or the time at which it exceeds a threshold or plateaus. We found it convenient to identify the “fast TF” as the one with the higher protein degradation rate. Specifically, we use the geometric mean of the protein degradation rate over gene copies of a TF in order to differentiate the two TFs. The parameter values of the fast TF are more evolutionarily constrained than those of the slow TF (**Table S5**). In particular, there is selection for rapid degradation of the fast TF protein and mRNA (**Table S5**). Isolated AND-gated C1-FFLs also show pronounced selection for the TF in the fast pathway to have rapid protein degradation (**Table S6**).

**Figure 8.**
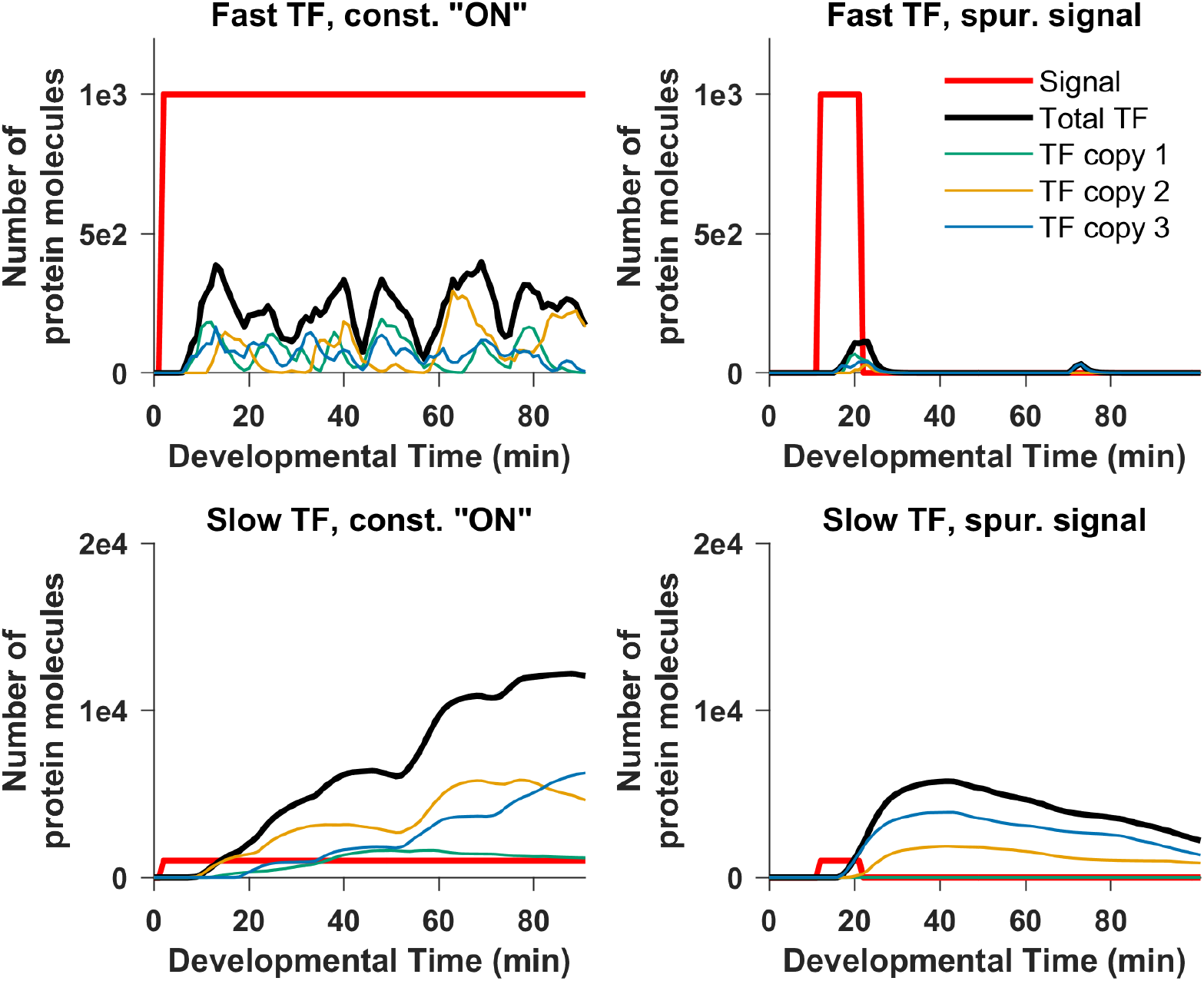
The two intermediate TFs in an AND-gated “diamond” motif have different expression dynamics and propagate the signal at different speeds. Expression of the two TFs in one representative genotype from the one high-fitness evolutionary replicate in **Fig. 7B** that evolved an AND-gated isolated diamond is shown. Each TF is a different protein, and each is encoded by 3 gene copies, shown separately in color, with the total in thick black. The expression of one TF plateaus faster than that of the other; this is characteristic of the AND-gated diamond motif, and leads to the same functionality as the AND-gated C1-FFL.

But mutational biases make it difficult to evolve very fast-degrading mRNA and protein. And even when they do evolve, fast degradation keeps the fast TF at low concentrations. To compensate, the fast TF must overcome mutational bias to also evolve high binding affinity and rapid protein synthesis (**Table S5**, **Table S6**).

Note that a simple transcriptional cascade, signal -> TF -> effector, has also been found experimentally to filter out short spurious signals when the intermediate TF is rapidly degraded, dampening the effect of a brief signal^47^. Two such transcriptional cascades involving different intermediate TFs form a diamond, so the utility of a single cascade is a potential explanation for the high prevalence of double-cascade diamonds. However, in this case we would have no reason to expect marked differences in expression dynamics between the two TFs, as illustrated in **Fig. 8** and **Table S5**. Enrichment for AND-gates (**Fig. 7**) indicates selection to integrate information from the two cascades. On the other hand, we do find some non-AND-gated diamonds, and these might best be considered as cascades. Inspection of their parameter values reveals that in these diamonds, both TFs have fast-degrading mRNAs and proteins so that both TFs shut down rapidly once signal is turned off. This makes such diamonds less vulnerable to spurious signals, reducing the need for the AND gate. The difficulty of evolving not just one but two fast-degrading high-affinity TFs likely explains why non-AND-gated diamonds are rare. As we will see in the next section, these non-AND-gated diamonds are nevertheless scored as AND-gated when weak TFBSs are excluded.

### Weak TFBSs can change how adaptive motifs are scored even when they do not change function

Results depend on whether we include weak TFBSs when scoring motifs. Weak TFBSs can either be in the effector’s cis-regulatory region, affecting how the regulatory logic is scored, or in TFs upstream in the TRN, affecting only the presence or absence of motifs. When a motif is scored as AND-gated only when two-mismatch TFBSs in the effector are excluded, we call it a “near-AND-gated” motif. Recall from **Fig. 2** that effector expression requires two TFs to be bound, with only one TFBS of each type creating an AND-gate. When a second, two-mismatch TFBS of the same type is present, we have a near-AND-gate. TFs may bind so rarely to this weak affinity TFBS that its presence changes little, making the regulatory logic still effectively AND-gated. A near-AND-gated motif may therefore evolve for the same adaptive reasons as an AND-gated one. **Fig. 7B** and **C** shows that both AND-gated and near-AND-gated motifs are enriched in the higher fitness genotypes.

When we exclude upstream weak TFBSs while scoring motifs, FFL-in-diamonds are no longer found, while the occurrence of isolated C1-FFLs and diamonds increases (**Fig. 7C**). This makes sense, because adding one weak TFBS, which can easily happen by chance alone, can convert an isolated diamond or C1-FFL into a FFL-in-diamond (added between intermediate TFs, or from signal to slow TF, respectively).

AND-gated isolated C1-FFLs appear mainly in the highest fitness outcomes, while AND-gated isolated diamonds appear in all fitness groups (**Fig. 7C**), suggesting that diamonds are easier to evolve. 25 out of 30 high-fitness evolutionary replicates are scored as having a putatively adaptive AND-gated or near-AND-gated motif in at least 50% of their evolutionary steps when upstream weak TFBSs are ignored (close to addition of bars in **Fig. 7C**, because these two AND-gated motifs rarely coexist in a high-fitness genotype).

Just as for the AND-gated C1-FFLs evolved under direct regulation and analyzed in **Fig. 5**, perturbation analysis supports an adaptive function for AND-gated C1-FFLs and diamonds evolved under indirect regulation (**Fig. 9A.i, 9B.i**). Breaking the AND-gate logic of these motifs by adding a (strong) TFBS to the effector cis-regulatory region reduces the fitness under the spurious signal but increases it under the constant “ON” beneficial signal, resulting in a net decrease in the overall fitness.

**Figure 9.**
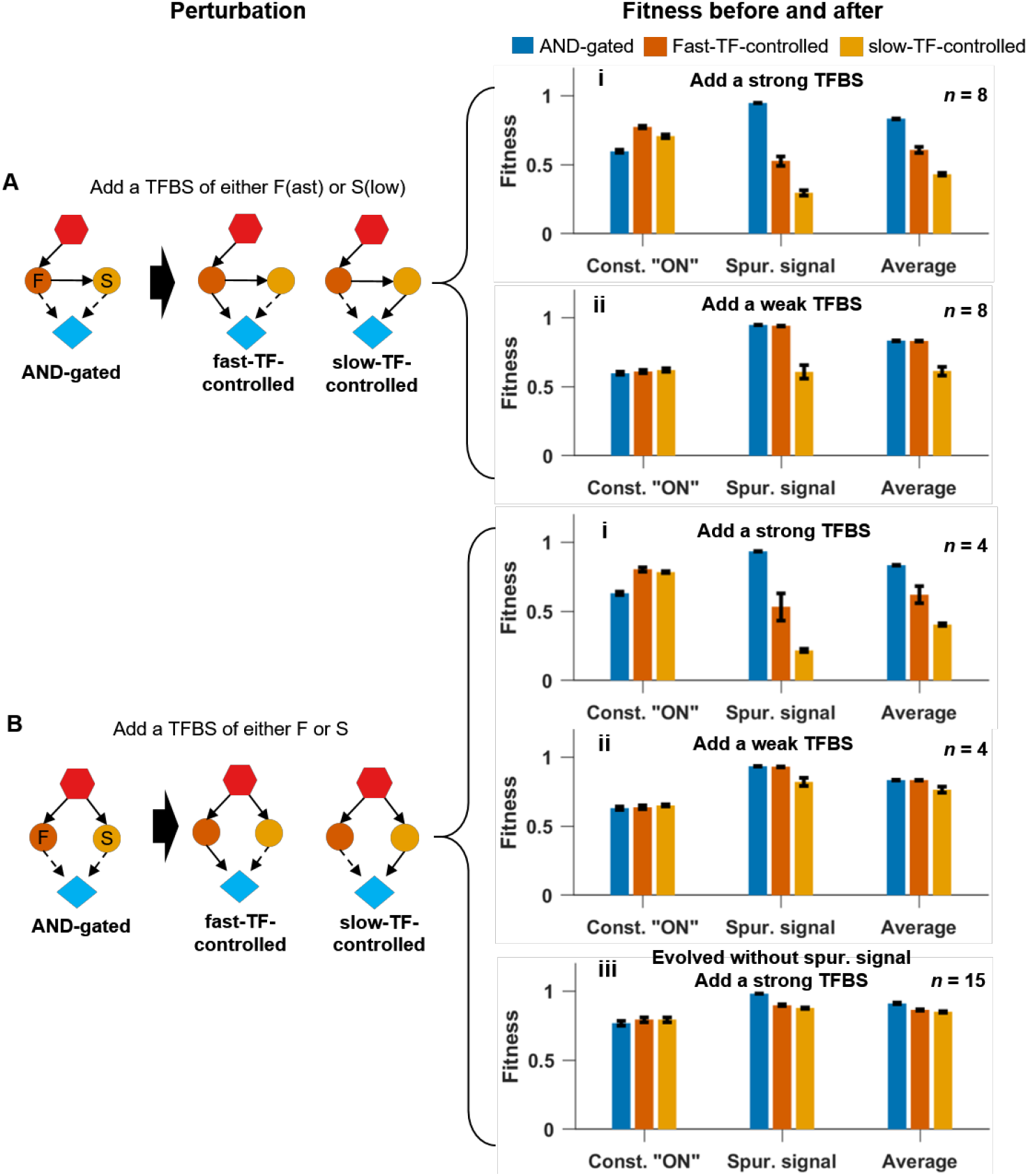
Perturbation analysis shows that AND-gated C1-FFLs (A) and diamonds (B) filter out short spurious signals. We add a strong TFBS **(i)** or a two-mismatch TFBS **(ii)** or **(iii)**; the latter creates near-AND-gated motifs. Allowing the effector to respond to the slow TF alone slightly increases the ability to respond to the signal, but leads to a larger loss of fitness when effector expression is undesirable. Allowing the effector to respond to the fast TF alone does so only when the conversion uses a strong TFBS not a two-mismatch TFBS. **(A)** We perform the perturbation on 8 of the 18 high-fitness replicates from **Fig.7B** that evolved an AND-gated C1-FFL. **(B)** (**i)** and (**ii**) are based on 4 of the 26 high-fitness replicates that evolved an AND-gated diamond in **Fig. 7B**, (**iii**) is based on 15 of the 37 replicates that evolved an AND-gated diamond in response to selection for signal recognition in the absence of an external spurious signal (**Fig. 10B**). Replicate exclusion was based on the co-occurrence of other motifs with the potential to confound results (see Supplementary Text Section 12 for details). Fitness is shown as mean±SE of over replicate evolutionary simulations, calculated as described for **Fig. 5**.

If we add a weak (two-mismatch) TFBS instead, this converts an AND-gated motif to a near-AND-gated motif. This lowers fitness only when the extra link is from the slow TF to the effector, and not when the extra link is from the fast TF to the effector (**Fig. 9A.ii, 9B.ii)**.

Indeed, these extra links are tolerated during evolution too. If we take the 16 high-fitness replicates that contain a near-AND-gated C1-FFL in at least 1% of the evolutionary steps, then for 15 replicates of the 16, at least 88% of the near-AND-gated C1-FFLs in each of the 15 replicates are only near-AND-gated because of extra weak TFBSs for the fast TF. In the remaining 1 replicate, 93% of the near-AND-gated C1-FFLs have extra weak TFBSs specific for each of the TFs (and are therefore scored as OR-gated). In this last replicate, the two TFs in these OR-gated C1-FFLs have high and similar protein degradation rates, reducing the need for an AND gate for reasons discussed earlier. We similarly examine high-fitness replicates that, when upstream weak TFBSs are excluded, contain a near-AND-gated diamond in at least 1% of the evolutionary steps. In 15 of these 24 evolutionary replicates, the near-AND regulatory logic is in most evolutionary steps due to an extra weak TFBS of the fast TF, in 8 replicates (all of them OR-gated, like the OR-gated C1-FFL already discussed) it is due to weak TFBSs for each of the TFs, and in only 1 replicate is it due to an extra TFBS for the slow TF. For the latter two categories, both TFs in near-AND-gated diamonds have high and similar protein degradation rates. By chance alone, fast and slow TF should be equally likely to contribute the weak TFBS that makes a motif near-AND-gated rather than AND-gated. This expected 50:50 ratio can be rejected from our observed 15:0 and 15:1 ratios with *p* = 3 × 10^-5^ and *p* = 3 × 10^-4^, respectively (cumulative binomial distribution, one-sided test). This non-random occurrence of weak TFBSs creating near-AND-gates illustrates how even weak TFBSs can be shaped by selection against some (but not all) motif-breaking links.

### AND-gated isolated diamonds also evolve in the absence of external spurious signals

We simulated evolution under the same three control conditions as before, this time without allowing the signal to directly regulate the effector. In the “no spurious signal” and “harmless spurious signal” control conditions, motif frequencies are similar between low and high fitness genotypes (**Fig. S6, Fig. S7)**, and so our analysis includes all evolutionary replicates. When weak (two-mismatch) TFBSs are excluded, AND-gated isolated C1-FFLs are seen only after selection for filtering out a spurious signal, and not under other selection conditions (**Fig. 10A**). However, AND-gated isolated diamonds also evolve in the absence of spurious signals, indeed at even higher frequency (**Fig. 10B**). Results including weak TFBSs are similar (**Fig. S8**).

**Figure 10.**
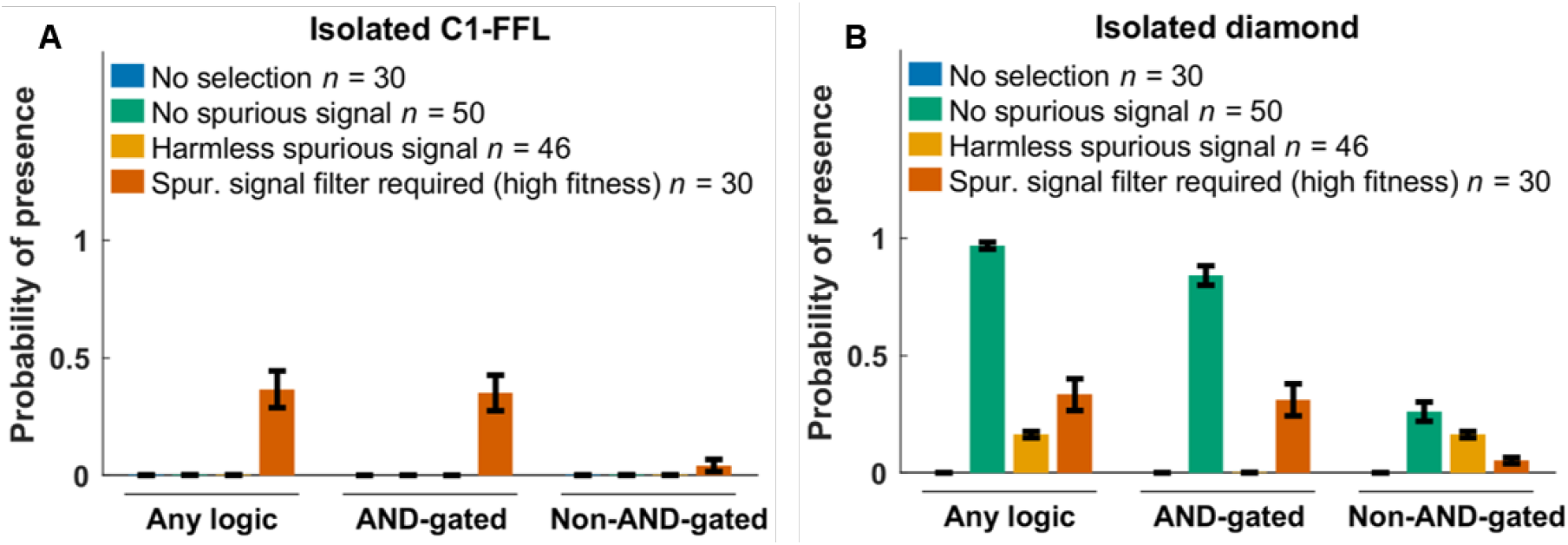
Selection for filtering out a short spurious signal is the primary way to evolve AND-gated isolated C1-FFLs (A), but AND-gated isolated diamonds also evolve in the absence of spurious signals (B). The selection conditions are the same as in **Fig. 6**, but we do not allow the signal to directly regulate the effector. When scoring motifs, we exclude all two-mismatch TFBSs; more comprehensive results are shown in **Fig. S8**. Many non-AND-gated diamonds have the “no regulation” logic in **Fig. 2**, perhaps as an artifact created by the duplication and divergence of intermediate TFs; we excluded them from the “Any logic” and “Non-AND-gated” tallies in (B). See Supplementary Text Section 11 for the calculation of y-axis. Data are shown as mean±SE over evolutionary replicates. We reused data from **Fig. 7** for “Spurious signal filter required (high fitness)”.

Perturbing the AND-gate logic in these isolated diamonds reduces fitness via effects in the environment where expressing the effector is deleterious (**Fig. 9B.iii**). Even in the absence of external short spurious signals, the stochastic expression of intermediate TFs might effectively create short spurious signals when the external signal is set to “OFF”. It seems that AND-gated diamonds evolve to mitigate this risk, but that AND-gated C1-FFLs do not. The duration of internally generated spurious signals has an exponential distribution, which means that the optimal filter would be one that does not delay gene expression^48^. The two TFs in an AND-gated diamond can be activated simultaneously, but they must be activated sequentially in an AND-gated C1-FFL; the shorter delays possible with AND-gated diamonds might explain why only diamonds and not FFLs evolve to filter out intrinsic noise in gene expression.

## Discussion

### Adaptive nature of AND-gated C1-FFLs

There has never been sufficient evidence to satisfy evolutionary biologists that motifs in TRNs represent adaptations for particular functions. Critiques by evolutionary biologists to this effect^13-23^ have been neglected, rather than answered, until now. While C1-FFLs can be conserved across different species^49-52^, this does not imply that specific “just-so” stories about their function are correct. In this work, we study the evolution of AND-gated C1-FFLs, which are hypothesized to be adaptations for filtering out short spurious signals^3^. Using a novel and more mechanistic computational model to simulate TRN evolution, we found that AND-gated C1-FFLs evolve readily under selection for filtering out a short spurious signal, and not under control conditions. Our results support the adaptive hypothesis about C1-FFLs.

AND-gated C1-FFLs express an effector after a noise-filtering delay when the signal is turned on, but shut down expression immediately when the signal is turned off, giving rise to a “sign-sensitive delay”^3, 7^. Rapidly switching off has been hypothesized to be part of their selective advantage, above and beyond the function of filtering out short spurious signals^48^. We intended to select only for filtering out a short spurious signal, and not for fast turn-off; specifically, we expected effector expression to evolve a delay equal to the duration of the spurious signal. However, evolved solutions still expressed the effector in the presence of short spurious signals (**Fig. S2**), and thus benefitted from rapidly turning off this spurious expression. In other words, we effectively selected for both delayed turn-on and rapid turn-off, despite our intent to only select for the former.

It is difficult to distinguish adaptations from “spandrels”^8^. Standard procedure is to look for motifs that are more frequent than expected from some randomized version of a TRN^2, 53^. For this method to work, this randomization must control for all confounding factors that are non-adaptive with respect to the function in question, from patterns of mutation to a general tendency to hierarchy – a near-impossible task. Our approach to a null model is not to randomize, but to evolve with and without selection for the specific function of interest. This meets the standards of evolutionary biology for inferring the adaptive nature of a motif^13-23^.

### Technical lessons learned

Previous studies have also attempted to evolve adaptive motifs in a computational TRN, successfully under selection for circadian rhythm and for multiple steady states^54^, and unsuccessfully under selection to produce a sine wave in response to a periodic pulse^23^. Other studies have evolved adaptive motifs in a mixed network of transcriptional regulation and protein-protein interaction^55-57^. Our successful simulation might offer some methodological lessons, especially a focus on high-fitness evolutionary replicates, which was done by us and by Burda et al.^54^ but not by Knabe et al.^23^.

Knabe et al.^23^ suggested that including a cost for gene expression may suppress unnecessary links and thus make it easier to score motifs. However, when we removed the cost of gene expression term (*C*(*t*) = 0 in Supplementary Section 8), AND-gated C1-FFLs still evolved in the high-fitness genotypes under selection for filtering out a spurious signal (**Fig. S9**). In our model, removing the cost of gene expression did not, via permitting unnecessary links, conceal motifs.

While simplified relative to reality, our model is undeniably complicated. An important question is which complications are important for what. One complication is our nucleotide-sequence-level model of cis-regulatory sequences. This has the advantage of capturing weak TFBSs, realistic turnover, and other mutational biases. The disadvantage is that calculating the probabilities of TF binding is computationally expensive and scales badly with network size. Future work might design a more schematic model of cis-regulatory sequences to improve computation while still capturing realistic mutation biases. A second complication of our approach is the stochastic simulation of gene expression. This is essential for our question, because intrinsic noise in gene expression can mimic the effects of a spurious signal, but may be less important in other scenarios, e.g. where the focus is on steady state behavior.

### The ubiquity of weak TFBSs complicates motif scoring

Our model, while complex for a model and hence capable of capturing intrinsic noise, is inevitably less complex than the biological reality. However, we hope to have captured key phenomena, albeit in simplified form. One key phenomenon is that TFBSs are not simply present vs. absent but can be strong or weak, i.e. the TRN is not just a directed graph, but its connections vary in strength. Our model, like that of Burda et al.^54^ in the context of circadian rhythms, captures this fact by basing TF binding affinity on the number of mismatch deviations from a consensus TFBS sequence. While in reality, the strength of TF binding is determined by additional factors, such as broader nucleic context and cooperative behavior between TFs (reviewed in Inukai et al.^58^), these complications are unlikely to change the basic dynamics of frequent appearance of weak TFBSs and greater mutational accessibility of strong TFBSs from weak TFBSs than de novo. Similarly, AND-gating can be quantitative rather than qualitative^59^, a phenomenon that weak TFBSs in our model provide a simplified version of.

Core links in adaptive motifs almost always involve strong not weak TFBSs. However, weak (two-mismatch) TFBSs can create additional links that prevent an adaptive motif from being scored as such. Some potential additional links are neutral while others are deleterious; the observed links are thus shaped by this selective filter, without being adaptive. Note that there have been experimental reports that even weak TFBSs can be functionally important^60, 61^; these might, however, better correspond to 1-mismatch TFBSs in our model than two-mismatch TFBSs. Ramos et al.^61^ and Crocker et al.^60^ identified their “weak” TFBSs in comparison to the strongest possible TFBS, not in comparison to the weakest still showing affinity above baseline.

### Different solutions for filtering out short spurious signals

A striking and unexpected finding of our study was that AND-gated diamonds evolved as an alternative motif for filtering out short spurious external signals, and that these, unlike FFLs, were also effective at filtering out intrinsic noise. Multiple motifs have previously been found capable of generating the same steady state expression pattern^21^; here we find multiple motifs for a much more complex function.

Diamonds are not overrepresented in the TRNs of bacteria^2^ or yeast^62^, but are overrepresented in signaling networks (in which post-translational modification plays a larger role)^63^, and in neuronal networks^1^. In our model, we treated the external signal as though it were a transcription factor, simply as a matter of modeling convenience. In reality, signals external to a TRN are by definition not TFs (although they might be modifiers of TFs). This means that our indirect regulation case, in which the signal is not allowed to directly turn on the effector, is the most appropriate one to analyze if our interest is in TRN motifs that mediate contact between the two. Note that if under indirect regulation we were to score the signal as not itself a TF, we would observe adaptive C1-FFLs but not diamonds, in agreement with the TRN data. However, this TRN data might miss functional diamond motifs that spanned levels of regulatory organization, i.e. that included both transcriptional and other forms of regulation. The greatest chance of finding diamonds within TRNs alone come from complex and multi-layered developmental cascades, rather than bacterial or yeast^64^. Multiple interwoven diamonds are hypothesized to be embedded with multi-layer perceptrons that are adaptations for complex computation in signaling networks^65^.

Previous work has also identified alternatives to AND-gated C1-FFLs. Specifically, in mixed networks of transcriptional regulation and protein-protein interactions, FFLs did not evolve under selection for delayed turn-on (as well as rapid turn-off)^57^. Indeed, even when a FFL topology was enforced, with only the parameters allowed to evolve, two alternative motifs remained superior^57^. However, one alternative motif, which the authors called “positive feedback” is essentially still an AND-gated C1-FFL, specifically one in which the intermediate TF expression is also AND-gated, requiring both itself and the signal for upregulation. The other is a cascade in which the signal inhibits the expression of an intermediate TF protein that represses the expression of the effector. The cost of constitutive expression of the intermediate TF in the absence of the signal was not modeled^57^, giving this cascade an unrealistic advantage.

### The importance of dynamics and intrinsic noise

Most previous research on C1-FFLs has used an idealized implementation (e.g. a square wave) of what a short spurious signal entails^4, 48, 66^. In real networks, noise arises intrinsically in a greater diversity of forms, which our model does more to capture. Even when a “clean” form of noise enters a TRN, it subsequently gets distorted with the addition of intrinsic noise^67^. Intrinsic noise is ubiquitous and dealing with it is an omnipresent challenge for selection. Indeed, we see adaptive diamonds evolve to suppress intrinsic noise, even when we select in the absence of extrinsic spurious signals.

The function of a motif relies ultimately on its dynamic behavior, with topology merely a means to that end. To create two pathways that regulate the effector in different speeds, the C1-FFL motif uses a pair of short and long pathways, but these also correspond to fast-degrading and slow-degrading TFs. This same function was achieved entirely non-topologically in our adaptively evolved diamond motifs. This agrees with other studies showing that topology alone is not enough to infer activities such as spurious signal filtering from network motifs ^68-70^.

## Acknowledgements

Work was supported by the University of Arizona and by a Pew Scholarship to JM, John Templeton Foundation grant 39667 to JM, and by National Institutes of Health grants R35GM118170 to MLS and R01GM076041 to JM. We thank Hinrich Boeger for helpful discussions and careful reading of the manuscript, Jasmin Uribe for early work on this project, and the high-performance computing center at the University of Arizona for generous allocations.

## Author Contributions

K.X. and J.M. designed the simulations, analyzed the results, and wrote the manuscript. K.X. performed the simulations and statistical analyses. K.X., A.L., and J.M. wrote the simulation code. M.L.S. and J.M. conceptualized the initial design of the simulations.

## Supporting Text

### 1 TF binding

Transcription of each gene is controlled by TFBSs present within a 150-bp cis-regulatory region, corresponding to a typical yeast nucleosome-free region within a promoter (Yuan et al. 2005). The perfect TFBS for a typical yeast TF has information content equivalent to 13.8 bits (Wunderlich & Mirny 2009); this means that in a simplified model of binding where only one of the four nucleotides is a good match at each site, ∼7 bp are recognized as an optimal consensus binding site. Maerkl & Quake (2007) reported that the TFBSs of two yeast TFs, Pho4p and Cbf1p, can have up to 2 mismatched sites within their 6 bp consensus binding sequence, while still binding the TF above background levels (Maerkl & Quake 2007). Our model therefore tracks TFBSs with up to 2 mismatches. This low information content implies a higher density of TFBSs within our cis-regulatory regions than our algorithm was able to handle, so we instead assigned each TF an 8-bp consensus sequence. Two TFs cannot simultaneously occupy overlapping stretches (Fig. S10), which we assume extend beyond the recognition sequence to occupy a total of 14 bp (Zhu & Zhang 1999); this captures competitive binding. The consequences of hindrance between TFBSs for the regulation of effector gene expression are shown in **Fig. 2**.

We denote the dissociation constant of a TFBS with *m* mismatches as *K*_*d*_(*m*). Sites with *m*>3 mismatches are assumed to still bind at a background rate equal to *m*=3 mismatches, with dissociation constant *K*_*d*_(3) = 10^-5^ mole/liter (Maerkl & Quake 2007) for all TFs. We assume that each of the last three base pairs makes an equal and independent additive contribution *ΔG*_*bp*_ < 0 to the binding energy (Benos et al. 2002): although not always true, this approximates average behavior well (Maerkl & Quake 2007). We ignore cooperativity in binding. Dissociation constants of eukaryotic TFs for perfect TFBSs can range from 10^-5^ mole/liter (Park et al. 2004) to 10^-11^ mole/liter (Nalefski et al. 2006). We initialize each TF with its own value of log_10_(*K*_*d*_(0)) sampled from a uniform distribution between −6 and −9, with mutation capable of further expanding this range, subject to *K*_*d*_(0) < 10^-5^ mole/liter. Substituting *m*=0 and *m*=3 into

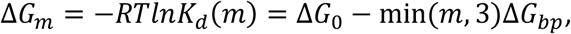

we can solve for *ΔG*_*bp*_ and Δ*G*_0_, and thus obtain *K*_*d*_(1) and *K*_*d*_(2) (the dissociation constants for TFBS with one and two mismatches, respectively).

Because TFs bind non-specifically to DNA at a high background rate, each nucleosome-free stretch of 14 bp can be considered to be a non-specific binding site (NSBS). A haploid *S. cerevisiae* genome is 12 Mb, 80% of which is wrapped in nucleosomes (Lee et al. 2007), yielding approximately 10^6^ potential non-specific binding sites (NSBSs). In a yeast nucleus of volume 3×10^-15^ liters, the NSBS concentration is of order 10^-4^ mole/liter. To find the concentration of free TF [TF] in the nucleus given a total nucleic TF concentration of *C*_*TF*_, we consider

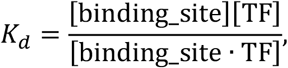

in the context of NSBSs, substitute [TF·NSBS] with *C*_*TF*_ - [TF], and solve for

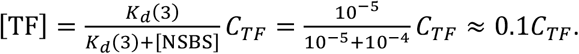

Thus, about 90% of total TFs are bound non-specifically, leaving about 10% free. The relatively small number of specific TFBSs is not enough to significantly perturb the proportion of free TFs, and so for the specific TFBSs with *m*<3 that are of interest in our model, we simply use 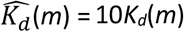 to account for the reduction in the amount of available TF due to non-specific binding. We also convert 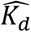 from the units of mole/liter in which *K*_*d*_ is estimated empirically to the more convenient molecules/nucleus. The rescaling factor *r* for which 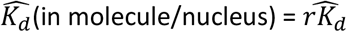 (in mole/liter) is 3×10^-15^ liter/nucleus × 6.02×10^23^ molecule/mole = 1.8×10^9^ molecule cell^-1^ liter mole^-1^. Taken together, 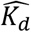 (molecule/nucleus) = 10*rK*_*d*_ (mole/liter), where the factor 10 accounts for non-specific TF binding.

### 2 TF occupancy

Here we calculate the probability that there are *A* activators and *R* repressors bound to a given cis-regulatory region at a given moment in developmental time. First we note that if we consider TF *i* binding to TFBS *j* in isolation from all other TFs and TFBSs, Eq. S1 gives us probability of being bound:

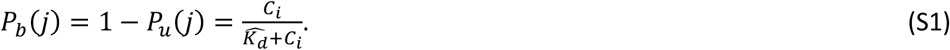

Let 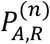 be a term proportional (for a given value of *R*) to the combined probability of all binding configurations in which exactly *A* activators and *R* repressors are bound to the first *n* binding sites along the cis-regulatory sequence. We calculate 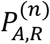 recursively, considering one additional TFBS at each step. Note that if two different TFs bind to exactly the same location on a cis-regulatory region, we treat this as two TFBSs, not as one, and treat first one and then the other in our recursive algorithm.

Consider the case where the (*n*+1)^th^ binding site belongs to an activator. The case where this activator is not bound contributes 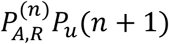. If it is bound, then we must also take into account that the (*n*+1)^th^ binding site overlaps (partially or completely) with the last *H* ≥ 0 sites, and so contributes 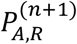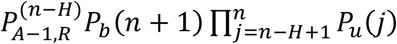. Taken together,

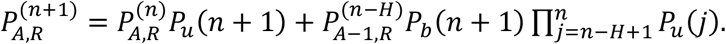

Similarly, if the (*n*+1)^th^ site belongs to a repressor, we have

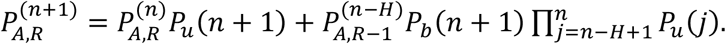

By definition, 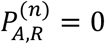 for binding configurations that are impossible, e.g. those with negative *A* or negative *R*. We initialize the recursion at *n* = 0, where the only valid binding configuration is for *A* = *R* = 0, i.e. 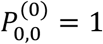. At 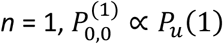, and if the binding site belongs to an activator, 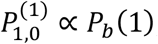; otherwise, 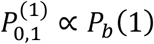. For *N* = 1, the two probabilities sum to 1 and normalization is unnecessary. For higher values of *N= N*_*A*_*+N*_*R*_ TFBSs, we normalize 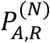 at the end of the recursion by dividing by 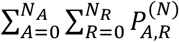 to get the probability of binding configurations that include exactly *A* activators and *R* repressors.

### 3. r_Act_to_Int_

Transcription initiation over an interval of time *r*_*transc_init*_ is proportional to the proportion of time spent in the Active state. Assuming a steady state between <u>Rep</u>ressed, <u>Int</u>ermediate, and <u>Act</u>ive states, as a function of current TF concentrations, we have:

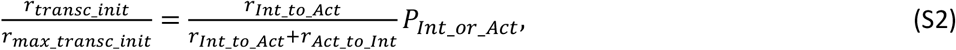

where *P*_*Int_or_Act*_ is the probability a gene is at Intermediate or Active. We set *r*_*max_transc_init*_ (the rate of transcription given 100% Active state) to 6.75 min^-1^, based on the corresponding rate when a model of the *PHO5* promoter is fit to data (Brown et al. 2013). In this model fit, the constitutively expressed *PHO5* promoter is free of nucleosomes 80% of the time, i.e. *P*_*Int_or_Act*_ = 0.8. We take these two values as universal for constitutively expressed genes, and assume that variation in *r*_*Act_to_Int*_ is responsible for variation in *r*_*transc_init*_. To identify a set of constitutively expressed genes, we identified 225 genes that have mRNA production rate of at least 0.5 molecule min^-1^ from genome-wide measurements (Pelechano et al. 2010); this threshold corresponds to low H2A.Z occupancy (Guillemette et al. 2005). We set *r*_*transc_init*_ to the production rate of mRNA of these 225 genes, and solve for gene-specific *r*_*Act_to_Int*_ from Eq. S2. We fit the solutions to a log-normal distribution and arrive at 10^*N*(1.27, 0.226)^ min^-1^.

To initialize values of *r*_*Act_to_Int*_ for each gene, we sample from this distribution. We also set lower and upper bounds for allowable values; if either the initial sample or subsequent mutation put *r*_*Act_to_Int*_ beyond these bounds, we set the value of *r*_*Act_to_Int*_ to equal to boundary value. We set the lower bound for *r*_*Act_to_Int*_ at 0.59 min^-1^, half the minimum of the values inferred from the set of 225 genes. To set an upper bound, we use the low H2A.Z occupancy bound of *r*_*transc_init*_ = 0.5, which gives a solution of 32.34 min^-1^; we double this to set the upper bound as 64.7 min^-1^.

### 4 Transcription delay times

Yeast protein lengths fit a log-normal distribution of 10^*N*(2.568, 0.34)^ amino acids (from the Saccharomyces Genome Database (SGD Project), excluding mitochondrial proteins). We sample ORF length *L* from this distribution. To constrain the values of *L*, we set a lower bound of 50 amino acids and an upper bound of 5000 amino acids; the longest protein in SGD is 4910 amino acids. If either initialization or mutation put *L* beyond these bounds, we set the value of *L* to the boundary value.

With an mRNA elongation rate of 600 codon/min (Larson et al. 2011; Hocine et al. 2013), it takes *L* / 600 minutes to transcribe the ORF of an mRNA. Also including time for transcribing UTRs and for transcription termination, and ignoring introns for simplicity, it takes 290 seconds to complete transcription of the yeast *GLT1* gene (Larson et al. 2011), whose ORF is 6.4kb. Putting the two together, we infer that transcribing the UTRs and terminating transcription takes around 1 minute for *GLT1*. Generalizing to assume that transcribing UTRs and terminating transcription takes exactly 1 minute for all genes, producing an mRNA from a gene of length *L* takes 1 + *L* / 600 minutes.

### 5 Translation delay times and *r*_*protein_syn*_

We model a second delay between the completion of a transcript and the production of the first protein from it. The delay comes from a combination of translation initiation and elongation; it ends when the mRNA is fully loaded with ribosomes all the way through to the stop codon and the first protein is produced. We ignore the time required for mRNA splicing; introns are rare in yeast (Dujon 1996). mRNA transportation from nucleus to cytosol, which is likely diffusion-limited (Niño et al. 2013; Smith et al. 2015), is fast even in mammalian cells (Mor et al. 2010) let alone much smaller yeast cells, and the time it takes is also ignored. The median time in yeast for initiating translation is 0.5 minute (Table 1 in Siwiak et al. 2010), and the genomic average peptide elongation rate is 330 codon/min (Siwiak et al. 2010). After an mRNA is produced, we therefore wait for 0.5 + *L* / 330 minutes, and then model protein production as continuous at a gene-specific rate *r*_*protein_syn*_.

To calculate *r*_*protein_syn*_, we combine the gene-specific ribosome densities *D* along the mRNAs and the gene-specific peptide elongation rates *E*, both measured in yeast (Siwiak et al. 2010). The values of *DE* across yeast genes fit the log-normal distribution 10^*N*(0.322, 0.416)^ molecule mRNA^-1^ min^-1^; we initialize *r*_*protein_syn*_ for each gene by sampling from this distribution. We set the lower bound for *r*_*protein_syn*_ at half the minimum observed value of *DE* (4.5×10^-3^ molecule mRNA^-1^ min^-1^). The upper bound corresponds to an mRNA fully occupied by rapidly moving ribosomes. Each ribosome occupies about 10 codons (Siwiak et al. 2010), and the peptide elongation rate can be as high as 614 codon/min (Waldron et al. 1977). If ribosomes are packed closely together at 10 codons apart, a protein comes off the end of production in the time taken to elongate 10 codons, i.e. proteins are produced at 61.4 molecules per minute. If either initialization or mutation put *r*_*protein_syn*_ beyond these bounds, we set the value of *r*_*protein_syn*_ to the boundary value.

### 6 mRNA and protein decay rates

We fit the log-normal distribution 10^*N*(−1.49, 0.267)^ min^-1^ to yeast mRNA degradation rates (Wang et al. 2002), and initialize the mRNA degradation rate *r*_*mRNA_deg*_ for each gene by sampling from this distribution. We set lower and upper bounds for *r*_*mRNA_deg*_ at half the minimum and twice the maximum observed values (7.5×10^-4^ min^-1^ and 0.54 min^-1^), respectively. If either initialization or mutation put *r*_*mRNA_deg*_ beyond these bounds, we set the value of *r*_*mRNA_deg*_ to the boundary value.

Expressing the estimated half-lives of yeast proteins (Belle et al. 2006) in terms of protein degradation rates, they fit the log-normal distribution 10^*N*(−1.88, 0.56)^ min^-1^; we initialize gene-specific protein degradation rates *r*_*protein_deg*_ by sampling from this distribution. We ignore the additional reduction in protein concentration due to dilution as the cell grows and thus increases in volume. We set lower and upper bounds for *r*_*protein_deg*_ at half the minimum and twice the maximum observed degradation rate (3.0 × 10^-6^ min^-1^ and 0.69 min^-1^), respectively. If either initialization or mutation put *r*_*protein_deg*_ beyond these bounds, we set the value of *r*_*protein_deg*_ to the boundary value.

### 7 Simulation of gene expression

Our algorithm is part-stochastic, part-deterministic. We use a Gillespie algorithm (Gillespie 1977) to simulate stochastic transitions between <u>Rep</u>ressed, <u>Int</u>ermediate, and <u>Act</u>ive chromatin states, and to simulate transcription initiation and mRNA decay events. We refer to these as “Gillespie events”. The completion of transcription to produce a complete mRNA, and subsequent ribosomal loading onto the mRNA, are referred to as “fixed events” (they require fixed times of 1 *+ L* / 600 minutes and 0.5 + *L* / 330 minutes, respectively). Scheduled changes in the strength of the external signal are also fixed events. Protein production and degradation are described deterministically with ODEs, and updated frequently in order to recalculate TF concentrations and hence chromatic transition rates. Updates occur at the time of Gillespie and fixed events, and also in between.

The total rate of all Gillespie events is

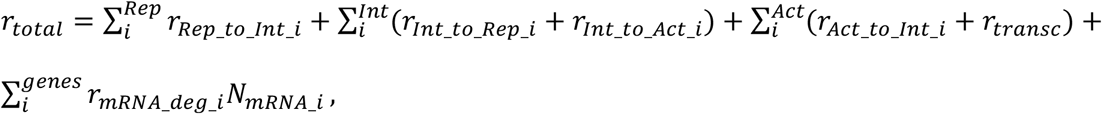

where *Rep, Int*, and *Act* are the numbers of gene copies in our haploid model that are in the Repressed, Intermediate, and Active chromatin states, respectively, and *N*_*mRNA_i*_ is the number of completely transcribed mRNA molecules from gene *i*. We only simulate degradation of full transcribed mRNA, and not that of mRNA that are still being transcribed, because the latter are already captured implicitly by *r*_*max_transc_init*_, which is based on mRNAs that complete transcription (Brown et al. 2013). Once an mRNA finishes transcription, it is subjected to degradation regardless of whether ribosome loading is complete.

The waiting time Δ*t* before the next Gillespie event is

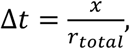

where *x* is random number drawn from an exponential distribution with mean 1. Which Gillespie event takes place next is sampled only if a different update does not happen first. If a fixed event is scheduled to happen first at *Δt*_*1*_ < *Δt*, we advance time by *Δt*_*1*_, update the state of the cell, and calculate a new *r*_*total*_*’.* Since the cellular activity has been going on with the old rate *r*_*total*_ for *Δt*_*1*_, the remaining “labor” required to trigger the Gillespie event planned earlier is reduced. The new waiting time, *Δt’*, to trigger the planned Gillespie event is

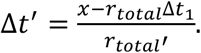

Gene duplication creates *R* ≥ 1 genes producing the same protein, where each copy *i* might have diverged in their production rate *r*_*protein_syn_i*_ and degradation rate *r*_*protein_deg_i*_. Complete proteins are produced continuously once an mRNA molecule is fully loaded with ribosomes, which occurs 0.5 + *L* / 330 minutes after transcription is complete – the concentration of such molecules is denoted *N*_*mRNA_aft_delay_i*_(*t*). Total protein concentration obeys:

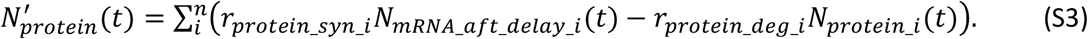

Protein concentrations are updated using a closed-form integral of Eq. S3

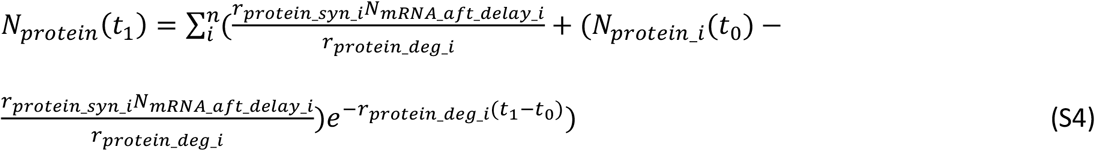

with this expression updated every time a Gillespie or fixed event at time *t*_*1*_ changes the value of *N*_*mRNA_aft_delay_i*_.

In between updates, values of *P*_*A*_, *P*_*R*_, *P*_*A_no_R*_, and *P*_*notA_no_R*_, and hence chromatin transition rates, are calculated under the approximation of constant *N*_*protein*_. Additional updates, above and beyond fixed and Gillespie events, are performed in order to ensure that chromatin transition rates do not change too dramatically from one update to the next. We use a target of *D* = 0.01 for the amount of change tolerated in the values of *P*_*A*_, *P*_*R*_, *P*_*A_no_R*_, and *P*_*notA_no_R*_, in order to schedule updates after time Δ*t*\*, which are triggered when neither a Gillespie event nor a fixed event occurs before this time has elapsed, i.e. when Δ*t*\* < Δ*t*_1_ and Δ*t*\* < Δ*t*.

There is the greatest potential for large changes after an update that changes the value of *N*_*mRNA_aft_delay_i*_. In this case, we use Eq. S1 to solve for the time interval for which the probability that TF *i* would be bound to a single perfect and non-overlapping TFBS would change by *D*, by choosing *Δt\** > 0 that satisfies

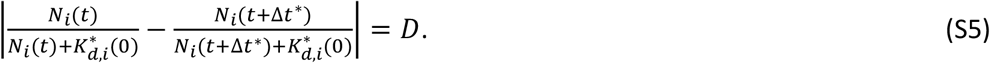

A solution for *Δt* may not exist, e.g. if the concentration of TF *i* is decreasing but *P*_*b*_(*t*_*2*_) < *D*. In such cases, we set Δ*t*\* to infinity.

When the previous update does not change any *N*_*mRNA_aft_delay_i*_ values, then we modify Δ*t*\* adaptively. Let *d* be the maximum of *ΔP*_*A*_, *ΔP*_*R*_, *ΔP*_*A_no_R*_, and *ΔP*_*notA_no_R*_ during the last update. We then schedule an update at

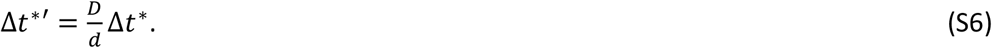

After an update that changes the value of *N*_*mRNA_aft_delay_i*_, we use the smaller value from Eqs. S5 and S6. These additional update times are discarded and recalculated when a Gillespie or fixed event occurs first.

In **Fig. S11**, we see that simulations rarely exceed our target of *D*=0.01, and do so only modestly.

### 8 Cost of gene expression

The cost of gene expression comes from some combination of the act of expression and from the presence of the resulting gene product. Yeast cells with plasmids carrying fast-degrading GFP had as much growth impairment as those carrying wild-type GFP (Fig. 3 of Kafri et al. 2016), suggesting that the former cost dominates. Universal costs stemming from the act of gene expression include the consumption of energy (Wagner 2005; Wagner 2007) and the opportunity cost of not using ribosomes to make other gene products (Scott et al. 2014). While some costs arise from transcription (Kafri et al. 2016), we simplify our model by attributing all of the cost of expression to the act of translation.

Kafri et al. (2016) reported that, in rich media, the growth rate of haploid yeast is reduced by about 1% when mCherry is expressed to about 2% of proteome. With *b*_*max*_ = 1 giving the growth rate of the yeast when mCherry is not expressed, we have the cost of gene expression equal to 0.01. Next, we estimate the production rate of mCherry in Kafri et al. (2016) by assuming that mCherry is in steady state between production and dilution due to cell division; fluorescent proteins tend to be stable such that degradation can be ignored (Snapp 2009). Ghaemmaghami et al. (2003) estimated that a haploid yeast cell contains about 5×10^7^ protein molecules, 2% of which are now mCherry. Over a 90 minute cell cycle in Kafri et al. (2016), about 5×10^5^ mCherry molecule per cell need to be expressed in order to double in numbers. This yields a production rate of about 5×10^3^ mCherry molecules per minute per cell. Because the total cost of gene expression is 0.01, the cost at a protein production rate of one mCherry molecule per minute per cell, *c*_*transl*_, is 2×10^-6^. Long genes should be more expensive to express than short ones; for a gene of length *L*, we assume its cost of expression is *c*_*transl*_ *L* / 370, where 370 is the geometric mean length of a yeast protein as described above in Section 4. Results using the length of mCherry instead, i.e. a slightly higher cost of expression of *c*_*transl*_ *L* / 236, are unlikely to be significantly different.

The overall cost of gene expression at time *t, C*(*t*) is:

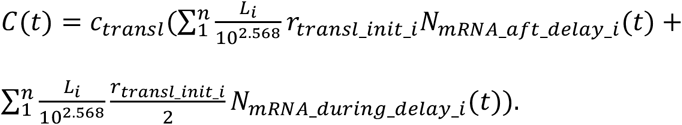

The second term represents transcripts that are on average half-loaded with ribosomes, and hence experiencing on average half the cost of translation. We integrate *C*(*t*) within segments of constant *C*(*t*) to obtain the overall cost of gene expression during a simulation.

### 9 Mutation

Because we use an origin-fixation approach, only the relative and not the absolute values of our mutation rates matter. In *S. cerevisiae*, the rates of small indels and of single nucleotide substitutions have been estimated as 0.2×10^-10^ per base pair and 3.3×10^-10^ per base pair, respectively (Lynch et al. 2008). Thus, cis-regulatory sequences are primarily shaped by single nucleotide substitutions. We do not model small indels in the cis-regulatory sequence, but increase the single nucleotide substitution up to 3.5×10^-10^ per base pair to compensate. This corresponds to a rate of 5.25×10^-8^ per 150 bp cis-regulatory sequence.

Lynch et al. (2008) also report a rate of gene duplication of 1.5×10^-6^ per gene and of deletion of 1.3×10^-6^ per gene (not including non-deletion-based loss of function mutations). These values turned out to swamp the evolution of TFBSs and hence significantly slow down our simulations, so we chose values 10-fold lower, making both gene duplication and gene deletion occur at rate 1.5×10^-7^ per gene. This preserves their numerical excess but reduces its magnitude.

Our model contains 8 gene-specific parameters, namely *L, r*_*Act_to_Int*_, *r*_*protein_deg*_, *r*_*protein_syn*_, *r*_*mRNA_deg*_, the *K*_*d*_(0) of a TF, whether a TF is an activator vs. repressor, and the consensus binding sequence of a TF. We assume mutations to *L* are caused by relatively neutral small indels, which we assume to be 20% of all small indels; mutation to *L* therefore occurs at rate 1.2×10^-11^ per codon, i.e. 1.2×10^-11^*L* for a gene of length *L*. For *r*_*Act_to_Int*_, we assume that it is altered by 10% of all the point mutations (single nucleotide substitution and small indels) to the core promoter of a gene. The length of a core promoter is about 100 bp and is relatively constant among genes (Roy & Singer 2015), yielding a mutation rate of *r*_*Act_to_Int*_ of 3.5×10^-9^ per gene.

The remaining 6 gene-specific parameter mutation rates are parameterized with lower accuracy due to lack to data; the principal decision is which to make dependent vs. independent of gene length. TF binding to DNA depends on particular peptide motifs whose length is likely independent of TF length, therefore we make mutation rates independent of gene length for mutations to *K*_*d*_(0), to the consensus binding sequence of a TF, and to the activating vs repressing identity of a TF. We set the rate of each of the three mutation types to 3.5×10^-9^ per gene. In contrast, because the stability of an mRNA mainly depends on its codon usage (Cheng et al. 2017) and thus more codons means more opportunities for change, we assume the rate of mutation to *r*_*mRNA_deg*_ does depend on gene length, as do mutations to protein stability *r*_*protein_deg*_. *r*_*protein_syn*_ is determined by the density of ribosomes on mRNA and the elongation rate of ribosomes, and therefore is affected both by ribosome loading speed and by slow spots forming queues in the mRNA. Ribosome loading often relies on the 5’UTR of mRNA (Hinnebusch 2011), and 5’UTR length is positively correlated with ORF length (Tuller et al. 2009). Slow-spots in mRNA can be due to secondary structure or to suboptimal codons, therefore are also more likely to appear by mutation to long mRNAs, so we assume the rate of mutation to *r*_*protein_syn*_ depends on gene length. We set the mutation rates of *r*_*protein_deg*_, *r*_*protein_syn*_, and *r*_*mRNA_deg*_ each to 9.5×10^-12^ per codon; in other words, each mutation rate is 3.5×10^-9^ for a yeast gene of average length (on a log-scale) 10^2.568^ = 370 codons.

*r*_*Act_to_Int*_, *r*_*protein_syn*_, *K*_*d*_(0), *r*_*protein_deg*_, and *r*_*mRNA_deg*_ evolve as quantitative traits. They are assumed to have, in the absence of selection, a log-normal stationary distribution with mean *µ* and standard deviation *σ*, with values estimated below and listed in **Table S2**. Denote the values of a parameter as *x* before mutation and *x’* after mutation; mutation takes the form:

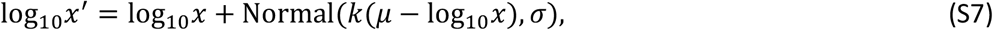

where *k* controls the speed of regressing back to the stationary distribution; we set *k* = 0.5 for all 5 parameters. To set values of *µ*, central tendency estimates of these five values (from **Table S1)** are adjusted according to our expectations about mutation bias. We assume a mutation bias toward faster mRNA degradation *r*_*mRNA_deg*_, faster *r*_*Act_to_Int*_ (Decker & Hinton 2013; Roy & Singer 2015), slower translation initiation *r*_*protein_syn*_ (Hinnebusch 2011), and larger *K*_*d*_(0). We assume that the observed log-normal means of *r*_*mRNA_deg*_, *r*_*protein_syn*_, and *r*_*Act_to_Int*_ differ by 2-fold from the mean expected from mutational bias; for example, the mean of log_10_(*r*_*mRNA_deg*_) is −1.49, so the value of *µ* for *r*_*mRNA_deg*_ is −1.49 + log_10_(2) = −1.19. We assume a larger bias for *K*_*d*_(0), namely that mutation is likely to reduce the affinity of a TF for a TFBS down to non-specific levels. Thus, we set *µ* = log_10_(*K*_*d*_(3)) = −5 for *K*_*d*_(0); note that in this case *µ* is equal to one of the boundary values, which will be hit far more often than during the evolution of other parameters. We assume that the observed central tendency estimate of protein stability does not depart from mutational equilibrium, therefore the value of *µ* for *r*_*protein_deg*_ is the mean of log_10_(*r*_*protein_deg*_) =-1.88.

The value of *σ* controls mutational effect size. We set the value of *σ* such that 1% of mutational changes from *x*=10^*µ*^ go beyond the boundary values, for simplicity approximating by considering only the closer of the two boundary values on a log scale, i.e. we solve Eq. S8 for *σ*:

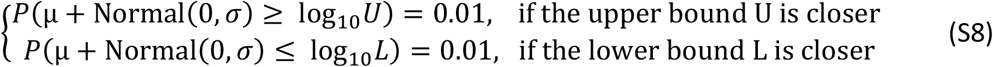

For example, the upper and the lower bounds of *r*_*mRNA_deg*_ are 0.54 min^-1^ and 7.5×10^-4^ min^-1^; on a log-scale, the upper bound is closer to 10^*µ*^ = 10^-1.19^ min^-1^. Plugging these values in Eq. S8 and solving for *σ*, we have *σ* = 0.396. We set the values of *σ* for *r*_*protein_syn*_, and *r*_*protein_deg*_ in the same way. However for *r*_*Act_to_Int*_, *σ* is set according to the lower bound, even though it is the more distant from 10^*µ*^, because otherwise a stable preinitiation complex will evolve too rarely. Under this high mutational variance, evolutionary outcomes at the two bounds are still only observed 5% of the time. For *K*_*d*_(0), because its upper bound is equal to 10^*µ*^, we set *σ* to 0.776, such that 1% of mutations can change the values of *K*_*d*_(0) by 100-fold or more.

Mutant values of *L, r*_*Act_to_Int*_, *r*_*protein_syn*_, *r*_*protein_deg*_, and *r*_*mRNA_deg*_ are constrained by the same bounds that constrain the initial values of these parameters (Sections 3-6). If a mutation increases the value of any of these 5 parameters to beyond the corresponding upper bound, we set the mutant value to the upper bound; similarly for a mutant value that is smaller than the lower bound of the corresponding parameter. For mutation to *K*_*d*_(0), we resample if *x’* ≥ *K*_*d*_(3), because otherwise the mutation effectively “deletes” the TF by reducing its affinity to non-specific levels.

### 10 Burn-in evolutionary simulation conditions

When the signal is not allowed to regulate the effector genes directly, most simulations under selection either to filter out short spurious signals or for simple signal recognition in the absence of spurious signals rapidly found a local optimal solution in which effector genes are never expressed. This local optimum exists in part because we assume that the environment in which the effector is deleterious is twice as common as the environment in which it is beneficial (**Fig. 3**). When the signal is not allowed to directly turn on the effector, then to escape this local optimum, at least one activator must be induced by the signal and then induce the effector. Such activators are rare when genotypes are randomly initialized. Making matters worse, mutation tends to reduce expression after initialization (see Section 9).

To reduce the frequency of this problem, we added a burn-in stage to simulations in which the signal is not allowed to regulate the effector directly. During burn-in, we switch the frequencies of the two environments, so that selection to express the effector is stronger. We also change the mutational bias in *r*_*Act_to_Int*_, *r*_*protein_syn*_, and *K*_*d*_(0) to favor higher expression and stronger binding. For *r*_*Act_to_Int*_ and *r*_*protein_syn*_, we use 0.1 instead of 0.01 as the tolerated fraction of extreme mutations in Eq. S8. For *K*_*d*_(0), we decrease *µ* from −5 to −7.5, biasing mutation toward the mean value at which we initialize (**Table S1**). Evolving an activator that can reliably turn on the effector when the signal is “ON” primarily relies on forming strong binding sites and appropriate kinetic constants in expression, assisted by the change in mutational bias above. To better focus the simulations on sampling appropriate mutations during the burn-in phase, we reduce the rate of gene duplication and the rate of deletion to 5.25 × 10^-9^ per gene, and limit the maximum number of TF genes to 9 and that of effector genes to 2. Each simulation is run under burn-in conditions for 1000 steps, after which normal model settings and selection conditions are restored. The same burn-in mutational settings are used for the control selection conditions (no selection, no spurious signal, and harmless spurious signal).

### 11 Quantifying occurrence of network motifs

Scoring the presence of a C1-FFL motif (e.g. **Fig. 4B**) or diamond motif (e.g. **Fig. 7**) is based on scoring whether TF *x* regulates gene *y*. Gene duplication and divergence complicate this scoring, because different gene copies might encode functionally identical proteins, but one copy of gene *y* might have a TFBS for TF *x* and the other might not. For the purpose of scoring motifs, our algorithm begins by simply treating each gene copy as though it were a unique gene.

Following Milo et al. (2002), a C1-FFL is scored if activating TF A can bind to the cis-regulatory sequence of activating TF B and to the effector, if B can also bind to that of the effector, and if B does not bind to that of A. Auto-regulation is allowed. We exclude C1-FFLs in which A and B encode the same TF or variants of the same TF. In the case of direct regulation, A can be the signal rather than a TF. C1-FFLs can then be subdivided into categories based on overlap between the TFBSs in the cis-regulatory region of the effector (**Fig. 2**).

A diamond is scored if two signal-regulated activating TFs, A and B, do not bind to each other’s cis-regulatory region, but both bind to that of the effector. We allow auto-regulation and require A and B to not encode the same TF or variants of the same TF.

A FFL-in-diamond is scored if one signal-regulated activating TF A binds to the cis-regulatory region of another signal-regulated activating TF B, but B does not bind to that of A, and both A and B bind to that of the effector. Again, auto-regulation is allowed, and A and B must not encode the same TF or variants of the same TF.

Occurrence within one evolutionary replicate is calculated as the fraction of the last 10,000 evolutionary steps in which at least one motif of the type of interest is present. The mean and standard error of this occurrence metric is then calculated across replicates.

### 12 Perturbing network motifs

In **Fig. 5** and **Fig. 9**, we add a TFBS to the cis-regulatory sequence of the effector gene, in order to destroy the AND-gate logic of an isolated C1-FFL or diamond. The new TFBS is chosen such that it does not overlap with any existing TFBSs, and has the same affinity as the strongest TFBS that is already present in the cis-regulatory sequence of the effector gene for the signal/fast TF (to convert from an AND-gate to signal-controlled/fast TF-controlled), or for the slow TF (to convert from an AND-gate to slow TF-controlled).

When a TRN has multiple AND-gated motifs of interest, we convert all of them. A perturbation can also affect the logic of other, potentially non-AND-gated motifs in the same TRN (e.g. **Fig. S12**), making it hard to attribute the fitness effect to the AND-gate logic of the targeted motif. For this reason, we perform the perturbation analysis not on a single potentially problematic genotype, but on the last 10,000 evolutionary steps of an evolutionary simulation. Within those 10,000 related genotypes, we exclude those that also contain other motifs that might influence our results. For simulations where the signal is allowed to directly regulate the effector, this means excluding those with non-AND-gated C1-FFLs. For simulations where the signal is not allowed to directly regulate the effector, we exclude genotypes with either AND-gated or non-AND-gated motifs other those of interest (e.g. if we intend to perturb AND-gated isolated C1-FFLs, we exclude genotypes that also contain either an AND-gated isolated diamond or a non-AND-gated C1-FFL). Both pre-perturbation fitness and post-perturbation fitness are averaged over the remaining genotypes. If no evolutionary step meets our requirement, we exclude the entire evolutionary simulation; this occurs only when the signal cannot directly regulate the effector genes.

## Supplementary Figures

**Fig. S1.**
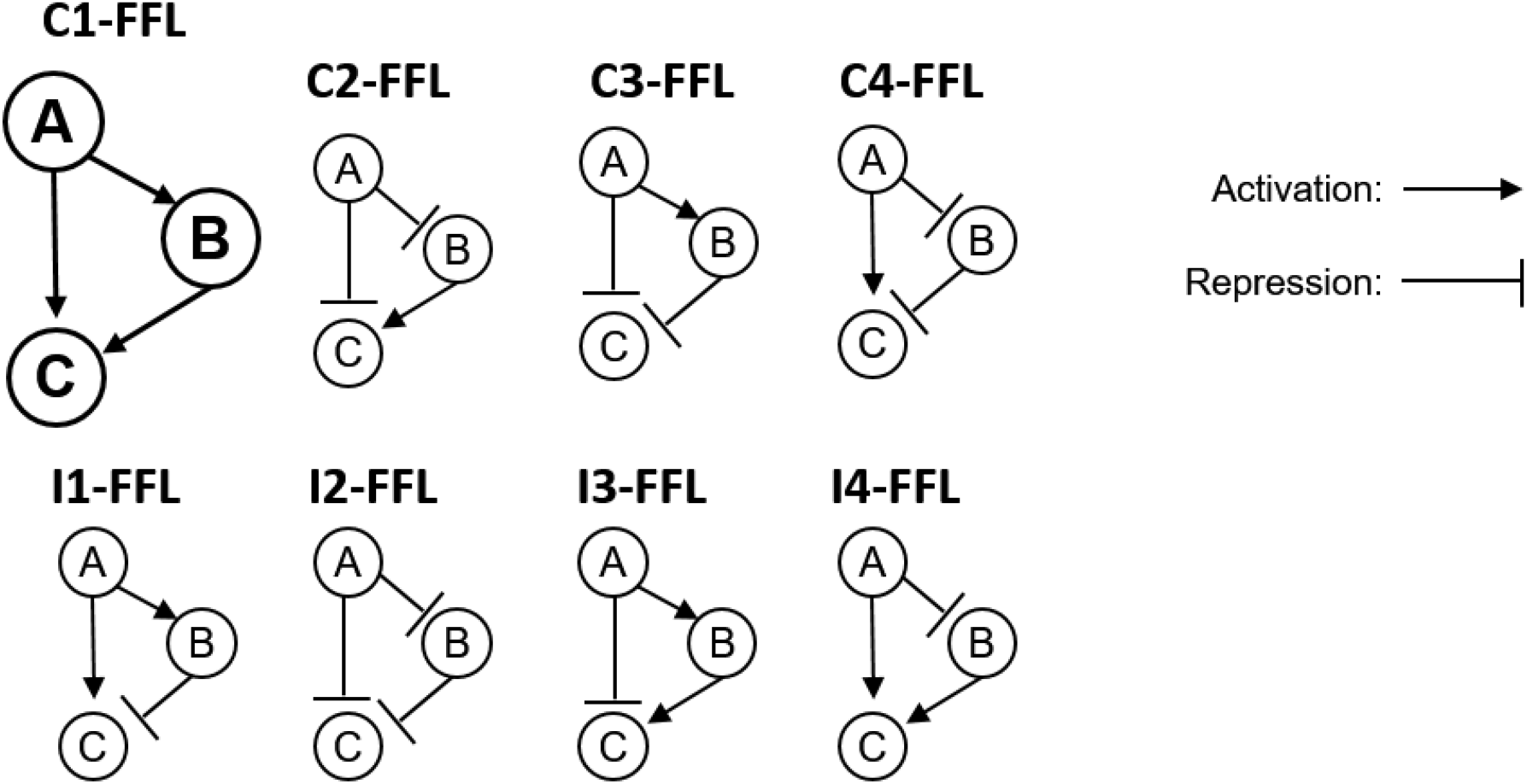
Feed-forward loops come in eight subtypes. TF A and TF B can activate (indicated by arrows) or repress (indicated by bars) expression of the effector C as well as other TFs. Auto-regulation is allowed, but not shown. Following Milo et al. (2002), we exclude the case in which A and B regulate one another, rather than treating this case as two overlapping FFLs. C stands for coherent and I for incoherent.

**Fig. S2.**
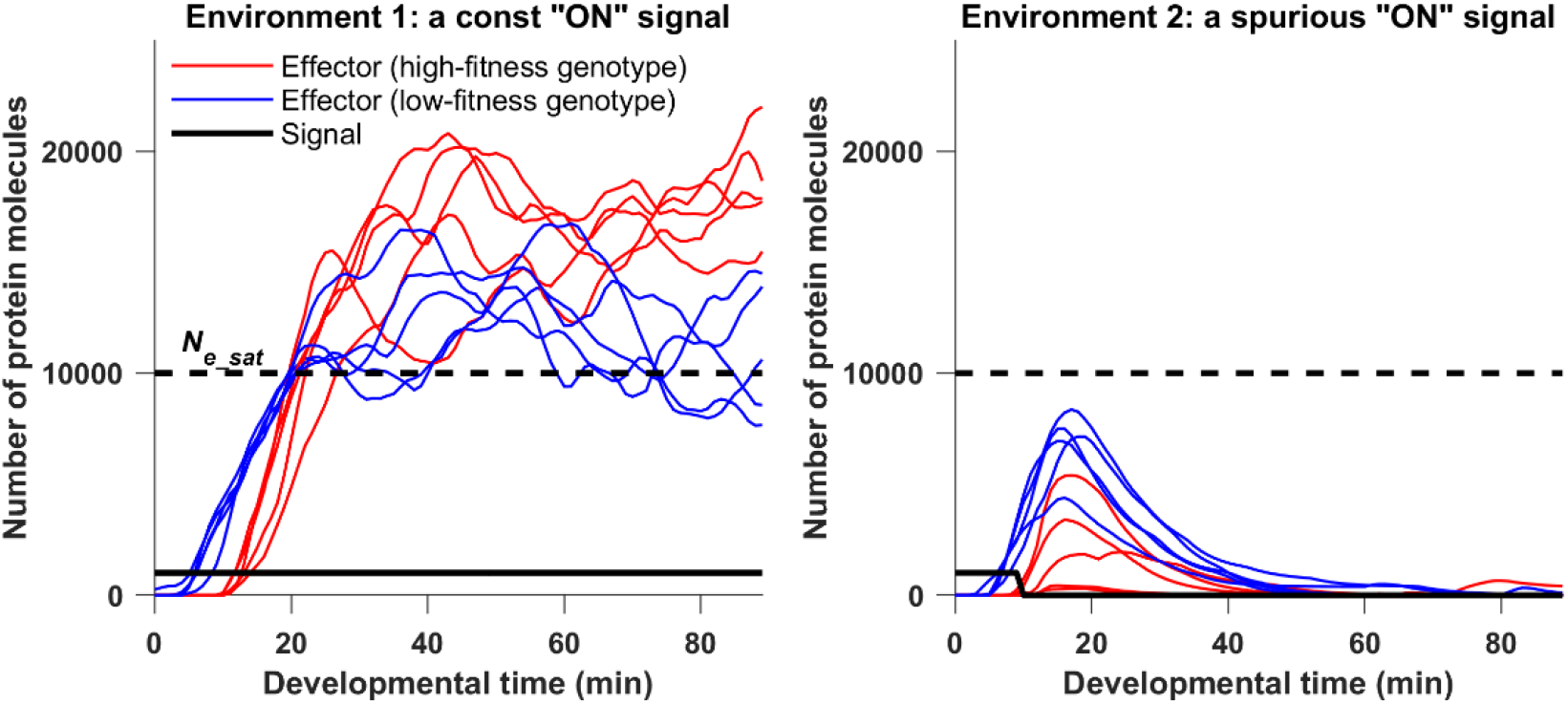
Examples of evolved phenotypes under selection for filtering out a short spurious signal. The figure shows trajectories of the effector protein in one randomly chosen high-fitness replicate (red) and one randomly chosen low-fitness replicate (blue), as defined in **Fig. 4A**. The genotype of the final evolutionary step is used, and other genotypes were confirmed to behave similarly. Each genotype is illustrated by 5 replicate developmental simulations in each of the two environments. The high-fitness genotype has a longer delay followed by more rapid response given a consistent signal, with this longer delay reducing but not eliminating effector expression given a short spurious signal. The signal is allowed to directly regulate the effector in these simulations. The burn-in period is not shown, with developmental time zero corresponding to the moment the signal is turned on. Among developmental replicates of the same genotype, the concentration at a given time usually has an approximately log-normal distribution, but in environment 2 the distribution has two modes after the spurious signal turns off. One mode corresponds to expression at the basal rate, the other to a burst of expression that has yet to turn off. Because of this bimodality, we plot sample trajectories rather than mean concentration over many replicates.

**Fig. S3.**
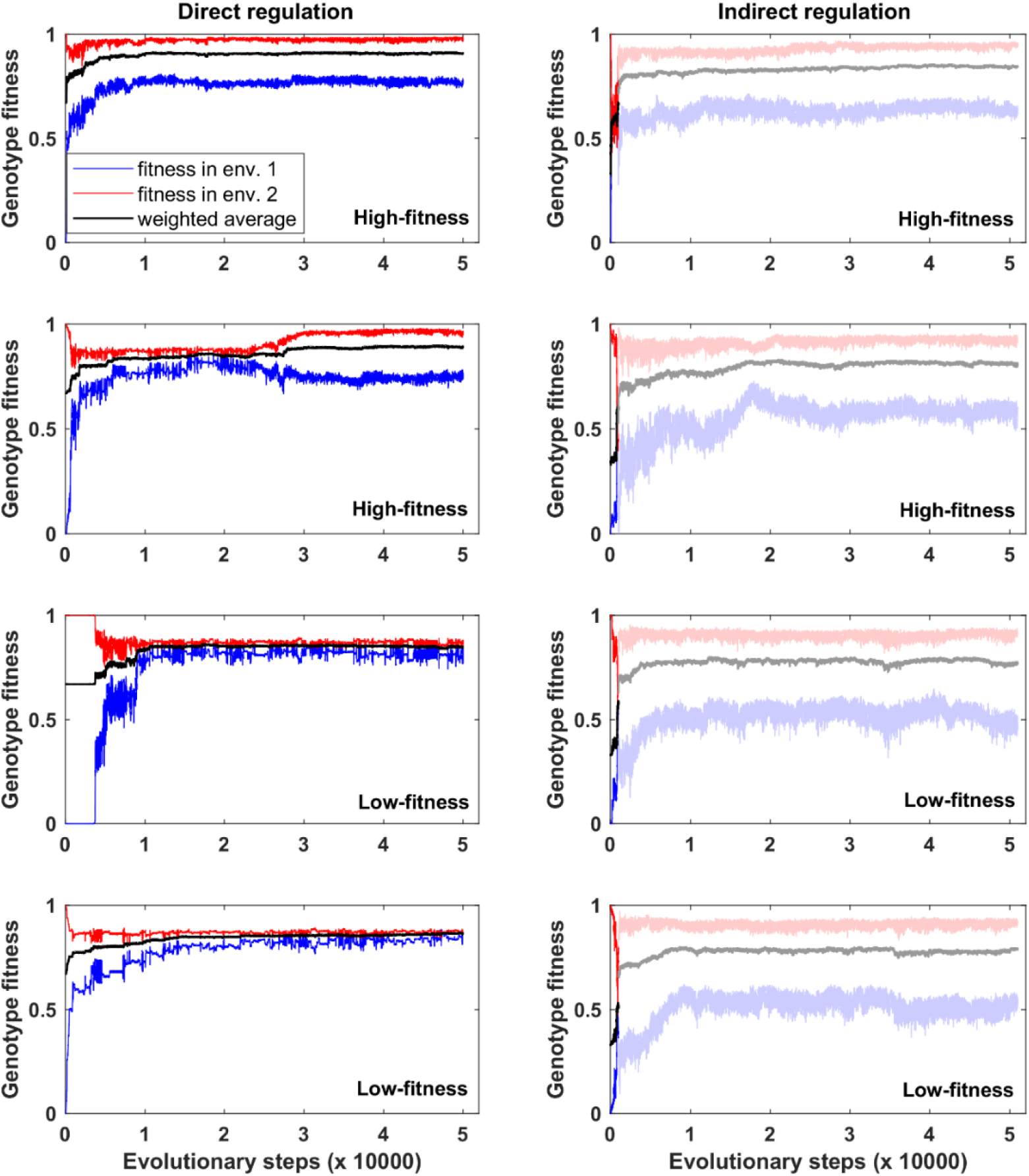
Representative fitness trajectories under selection to filter out short spurious signals. Left panels: The signal is allowed to directly regulate the effector genes. Panels 1 and 3 correspond to the two genotypes shown in **Fig. S2**. Right panels: the signal cannot directly regulate the effector genes. Average fitness (black) is a weighted average of the blue and red trajectories, with environment 2 (where the signal is spurious) being considered twice as common as environment 1 (where the signal is sustained and real). When the signal cannot directly regulate the effector genes, evolutionary simulations begins with a burn-in phase that lasts 1000 evolutionary steps (see Evolutionary Simulation in the Main Text). We show the burn-in phase in undiluted color, and dilute color after burn-in. Most replicates quickly reach a stable fitness plateau (first and third rows). Certain replicates can be temporarily trapped at a low fitness plateau (second and third rows on the left).

**Fig. S4.**
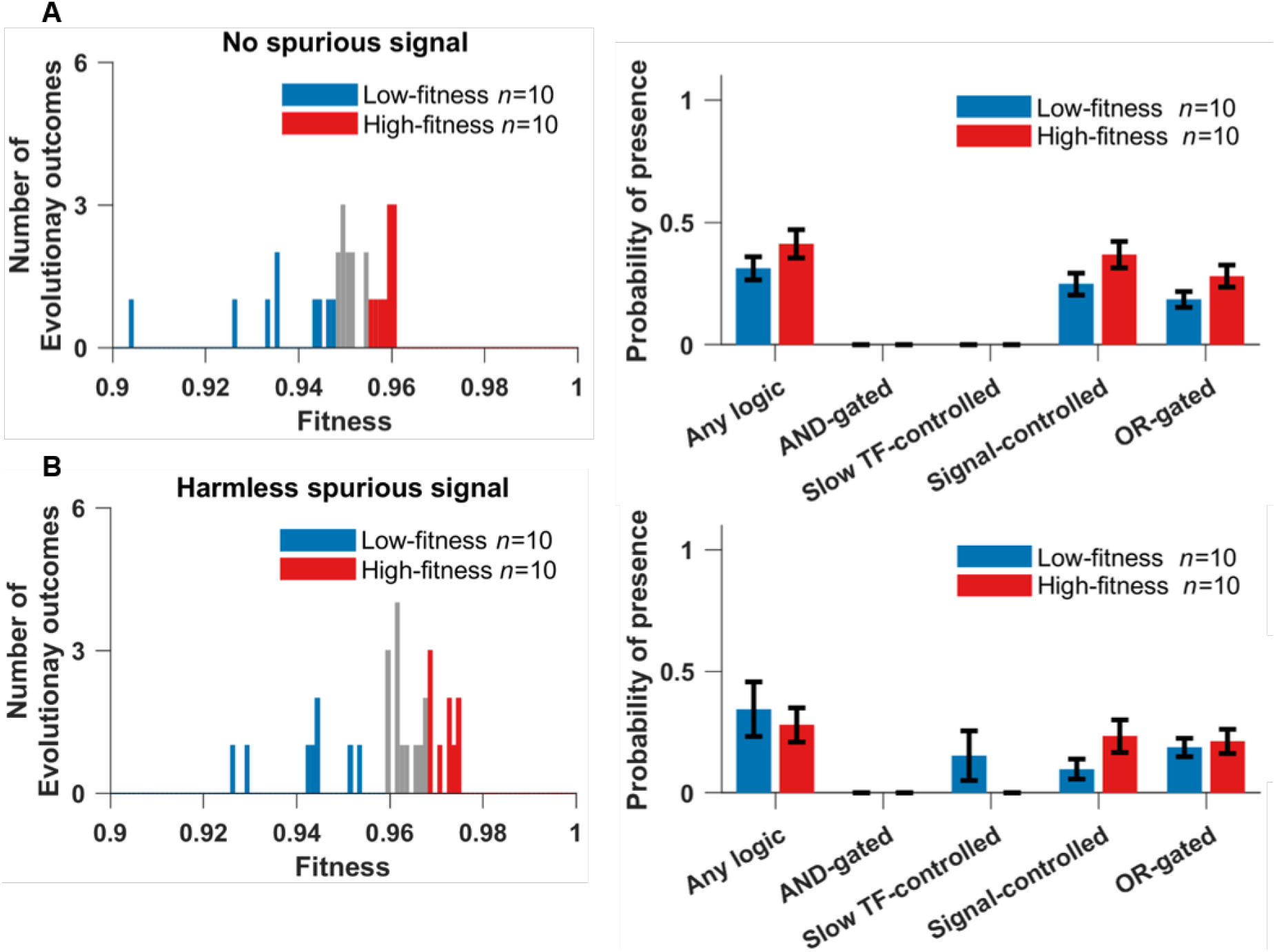
Genotypes evolved under control selective conditions: (A) “no spurious signal”, and (B) “harmless spurious signal”. There is no clear evidence of a multimodal distribution of fitness outcomes among replicates (left), and C1-FFLs occur equally in the 10 genotypes of the highest fitness vs. the 10 genotypes of the lowest fitness (right), and so the entire distribution (left) was used to produce **Fig. 6**. Data are shown as mean±SE over evolutionary replicates.

**Fig. S5.**
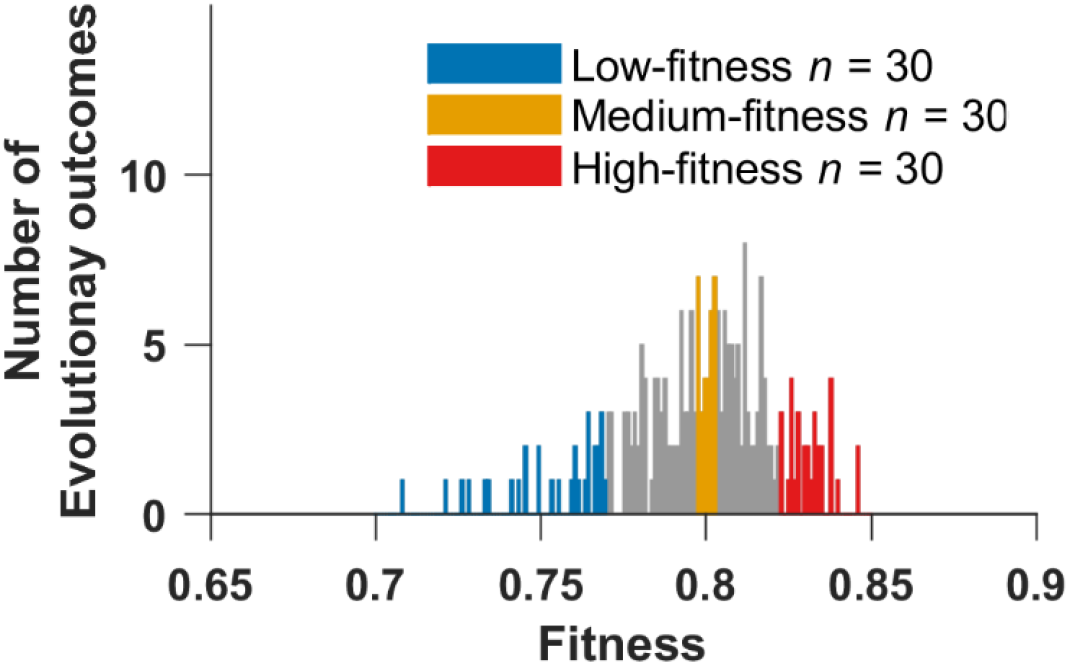
Fitness distribution of 258 evolutionary replicates under selection for filtering out short spurious signals, when the signal cannot directly regulate the effector. The fitness of a replicate is the average genotype fitness over the last 10,000 evolutionary steps. Colors indicate replicates analyzed elsewhere.

**Fig. S6.**
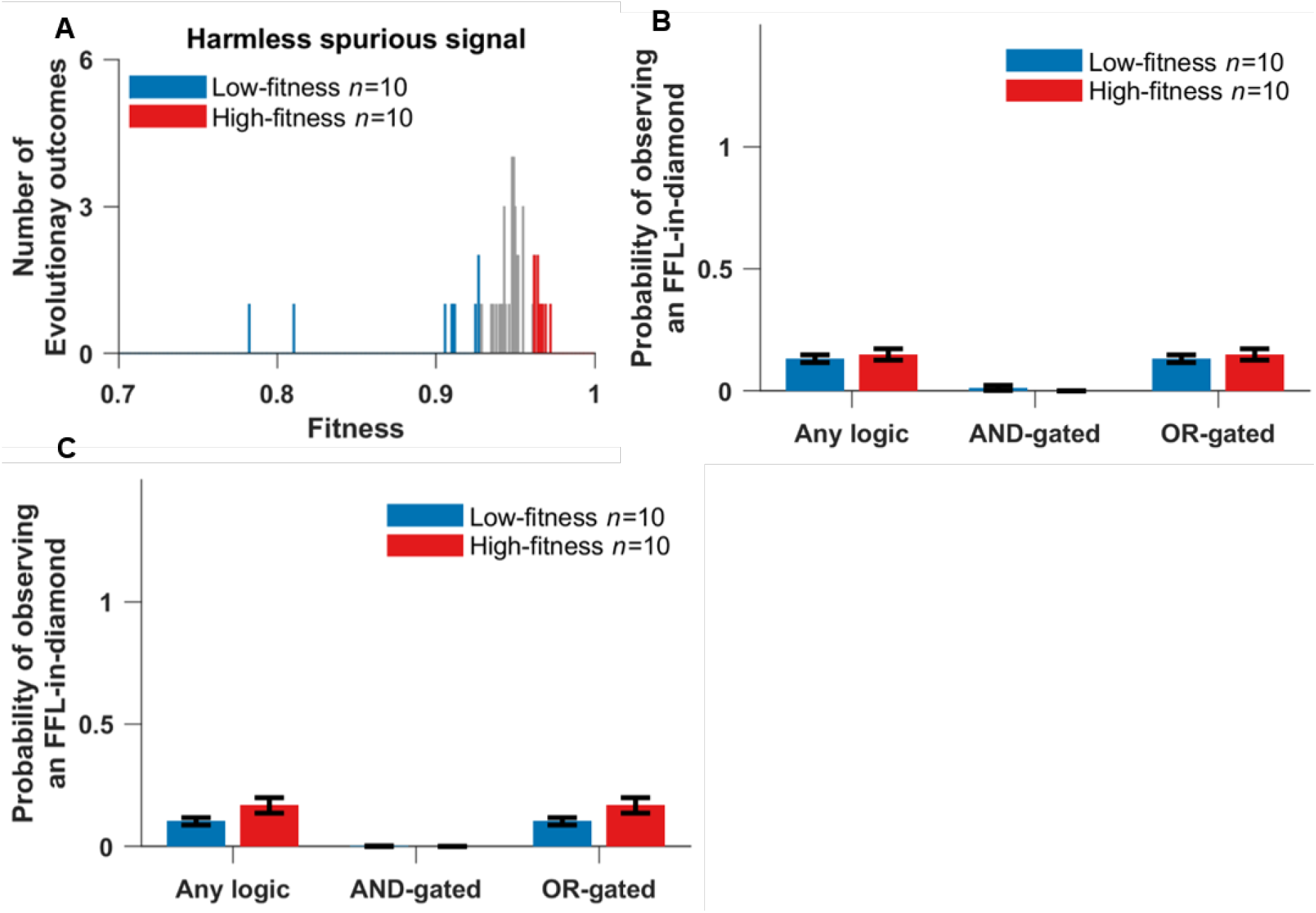
Evolution when responding to a spurious signal is harmless, when the signal is not allowed to directly regulate the effector. **(A)** Fitness distribution of 50 replicate simulations. The occurrence of both **(B)** FFL-in-diamonds and **(C)** isolated diamonds were similar in the 10 genotypes with the highest fitness vs. in 10 genotypes with the lowest fitness. Weak (two-mismatch) TFBSs are included when scoring motifs. Data are shown as mean±SE over replicates. Isolated C1-FFLs rarely evolve under this condition, therefore their occurrence is not plotted.

**Fig. S7.**
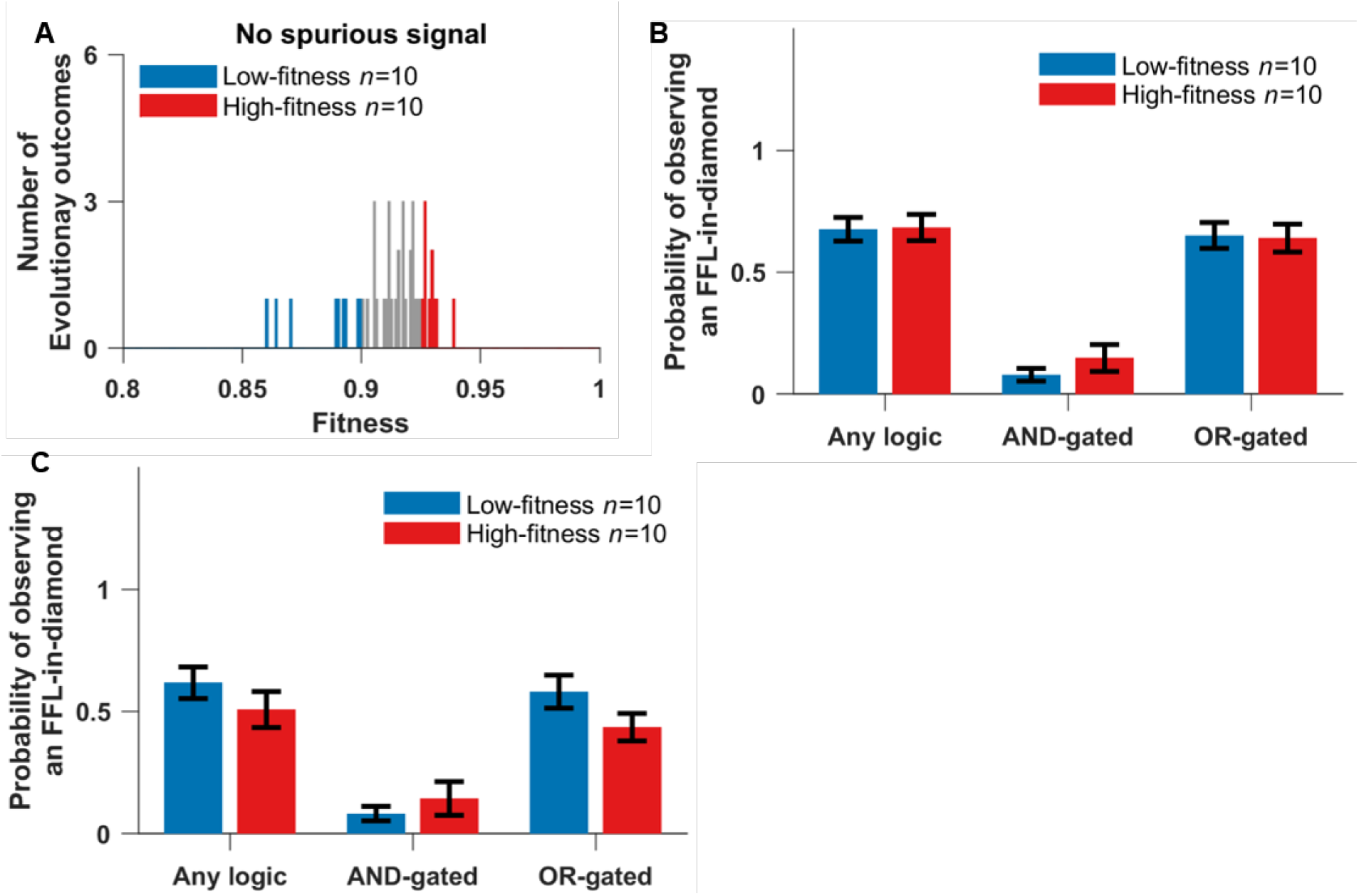
Evolution when there is no spurious signal, when the signal is not allowed to directly regulate the effector. **(A)** Fitness distribution of 46 replicate simulations. The occurrence of both **(B)** FFL-in-diamonds and **(C)** isolated diamonds were similar in the 10 genotypes with the highest fitness vs. in the 10 genotypes with the lowest fitness. Weak (two-mismatch) TFBSs are included when scoring motifs. Data are shown as mean±SE over replicates. Isolated C1-FFLs rarely evolve under this condition, therefore their occurrence is not plotted.

**Fig. S8.**
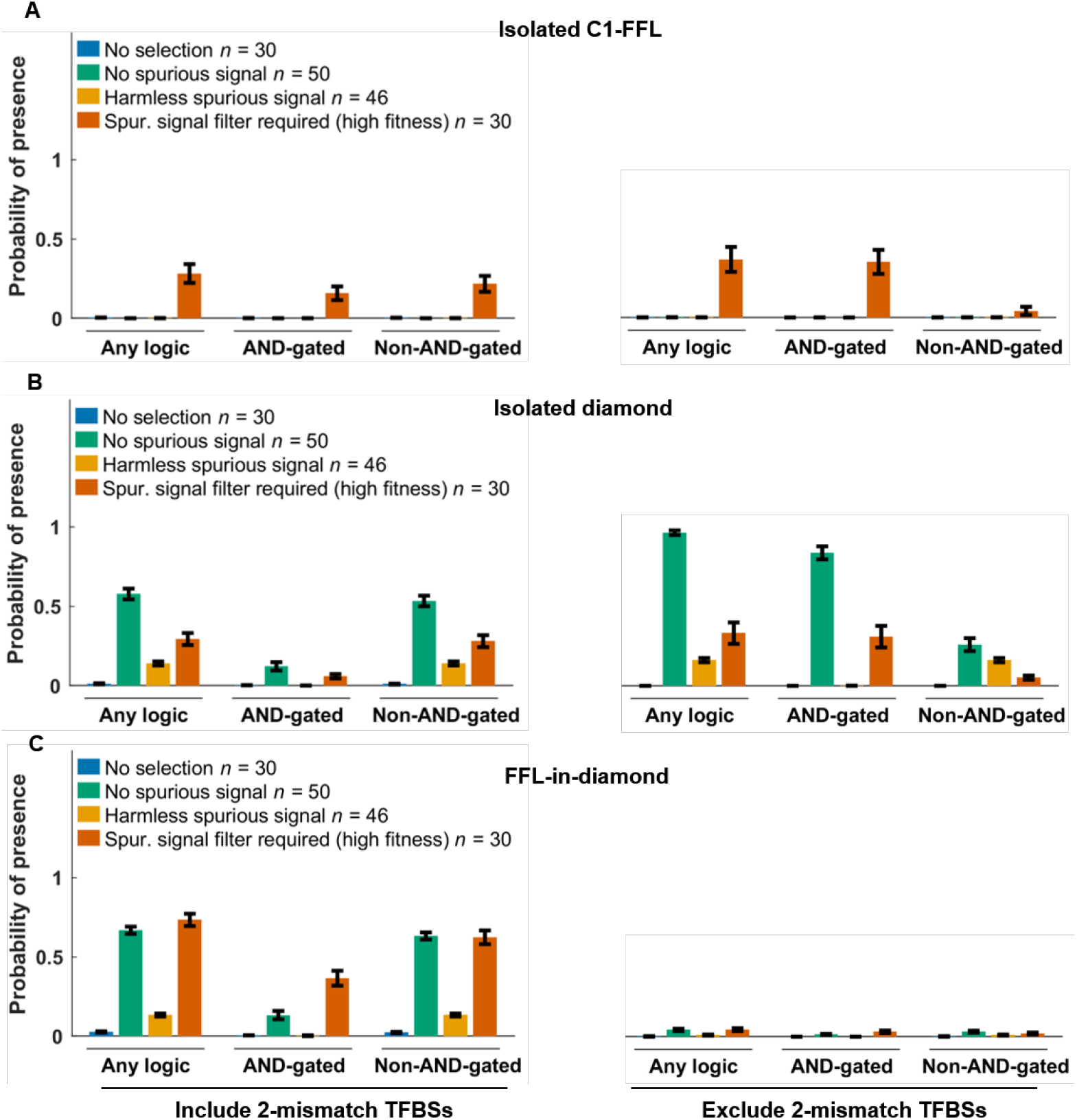
Selection for filtering out a short spurious signal is the primary way to evolve AND-gated C1-FFLs (A), but AND-gated isolated diamonds also evolve in the absence of spurious signals (B). The signal is not allowed to directly regulate the effector, and the right panels of **(A)** and **(B)** are identical to **Fig. 10**. When scoring motifs, we either include (left) or exclude (right) all two-mismatch TFBSs in the cis-regulatory sequences of intermediate TF genes and effector genes. We excluded “no regulation” (**Fig. 2**) diamonds from the “Any logic” and “Non-AND-gated” tallies in **(B)**; this was necessary because of their high occurrence due to duplication and divergence of intermediate TFs. See Section 11 for the calculation of y-axis. Data are shown as mean±SE over evolutionary replicates.

**Fig. S9.**
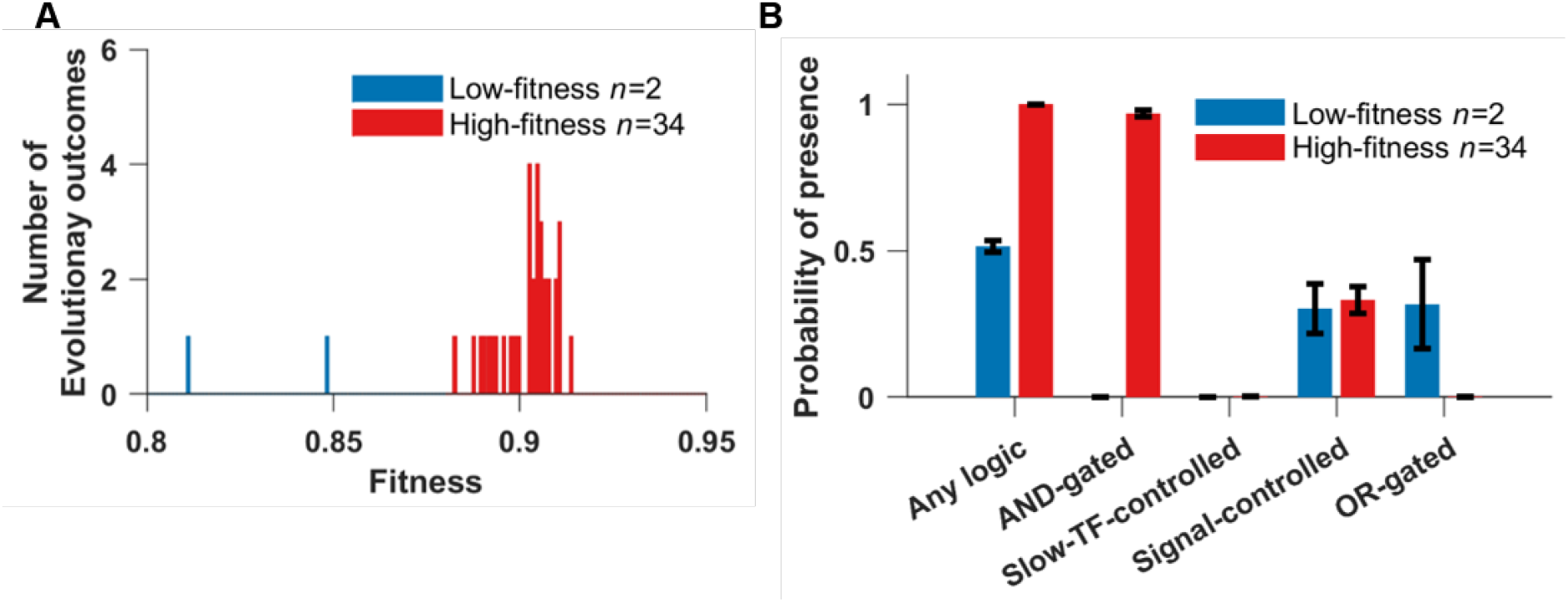
After removing cost of gene expression, AND-gated C1-FFLs are still associated with a successful response to selection for filtering out a short spurious signal. The signal can directly regulate the effector genes. **(A)** We arbitrarily divide the 36 replicate simulations into high-fitness (red) and low-fitness (blue) groups. **(B)** The high-fitness replicates still evolve AND-gated C1-FFLs. Bars are mean±SE of the occurrence over replicate evolutionary simulations.

**Fig. S10.**
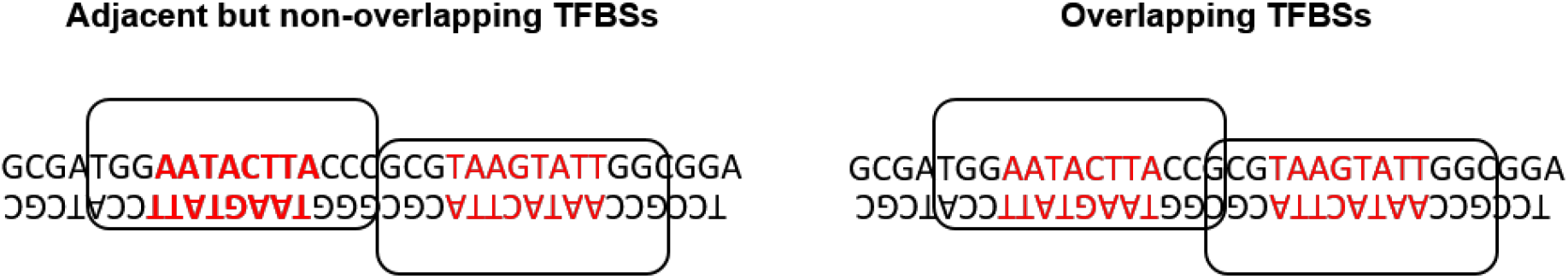
TFs (white boxes) recognize 8 bp (red) sites while occupying and thus excluding other TFs from a 14 bp long space. TFs are assumed to bind in either orientations (Sharon et al. 2012). The sequence on the left allows simultaneous binding but that on the right does not.

**Fig. S11.**
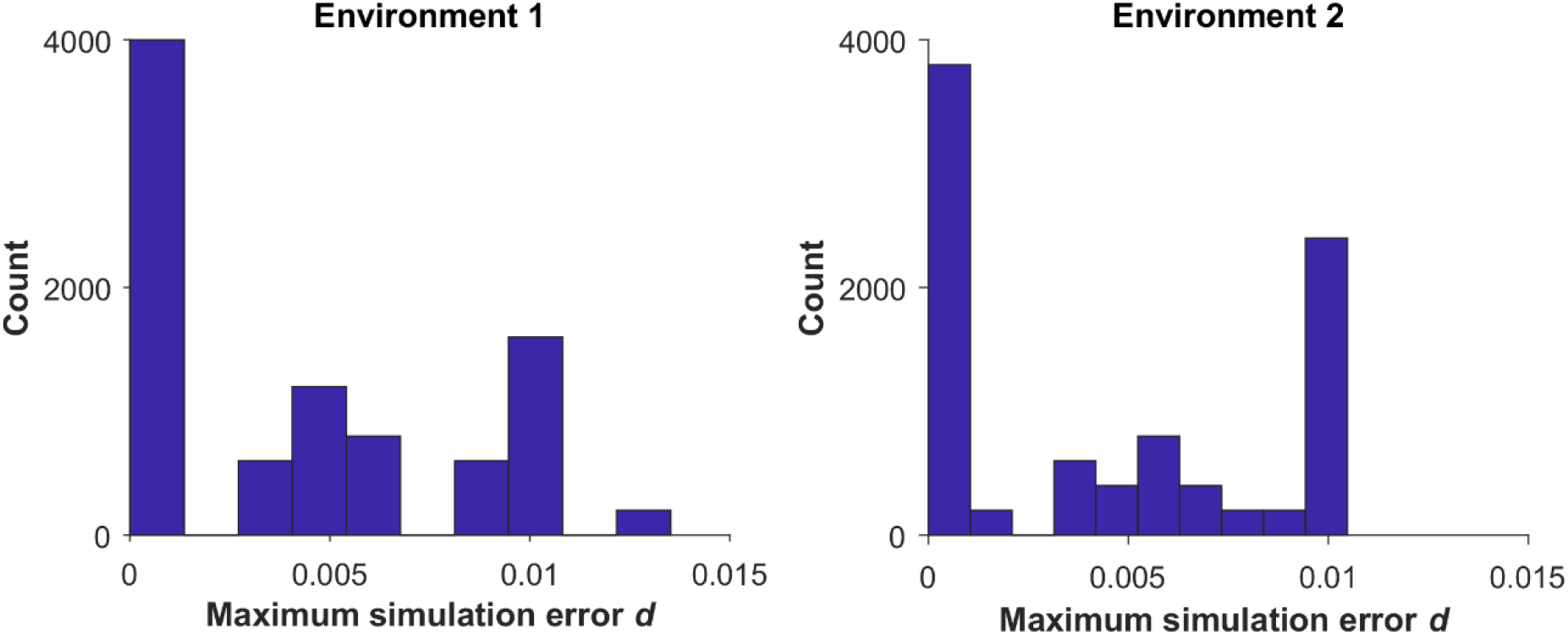
Our updating algorithm is able to limit simulation errors. The distribution across 9,000 simulations of the maximum value of *d* over the course of development. For each of the 45 evolutionary replicates in **Fig. 4**, we run 200 simulations of development of the final evolved genotype. These genotypes were the outcome of evolution under selection for filtering out short spurious signals, in which direct regulation of the effector by the signal is not allowed. In environment 1 a genotype responds to a constant “ON” signal and in environment 2 it responds to a short spurious signal (**Fig. 3**).

**Fig. S12.**
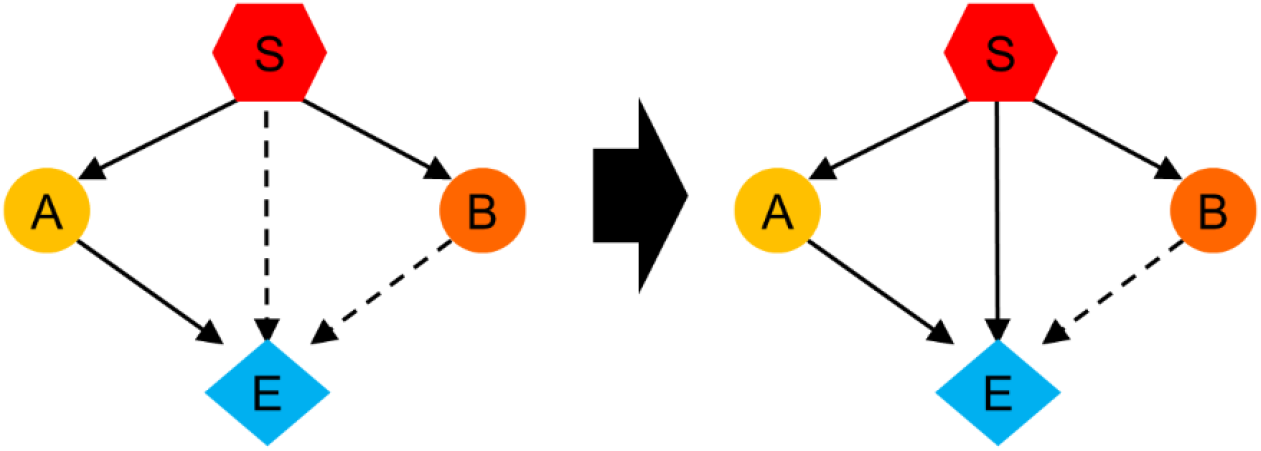
Examples of confounding motifs in perturbation analysis. The TRN on the left contains a slow TF-controlled C1-FFL (S-B-E) and an AND-gated C1-FFL (S-C-E). To convert S-C-E into a signal-controlled C1-FFL, we need to add one TFBS for the signal to the cis-regulatory sequence of E. However, this change also makes S-B-E OR-gated, making it difficult to conclude whether it is the AND gate logic of S-B-E that matters for fitness.

**Table S1.**
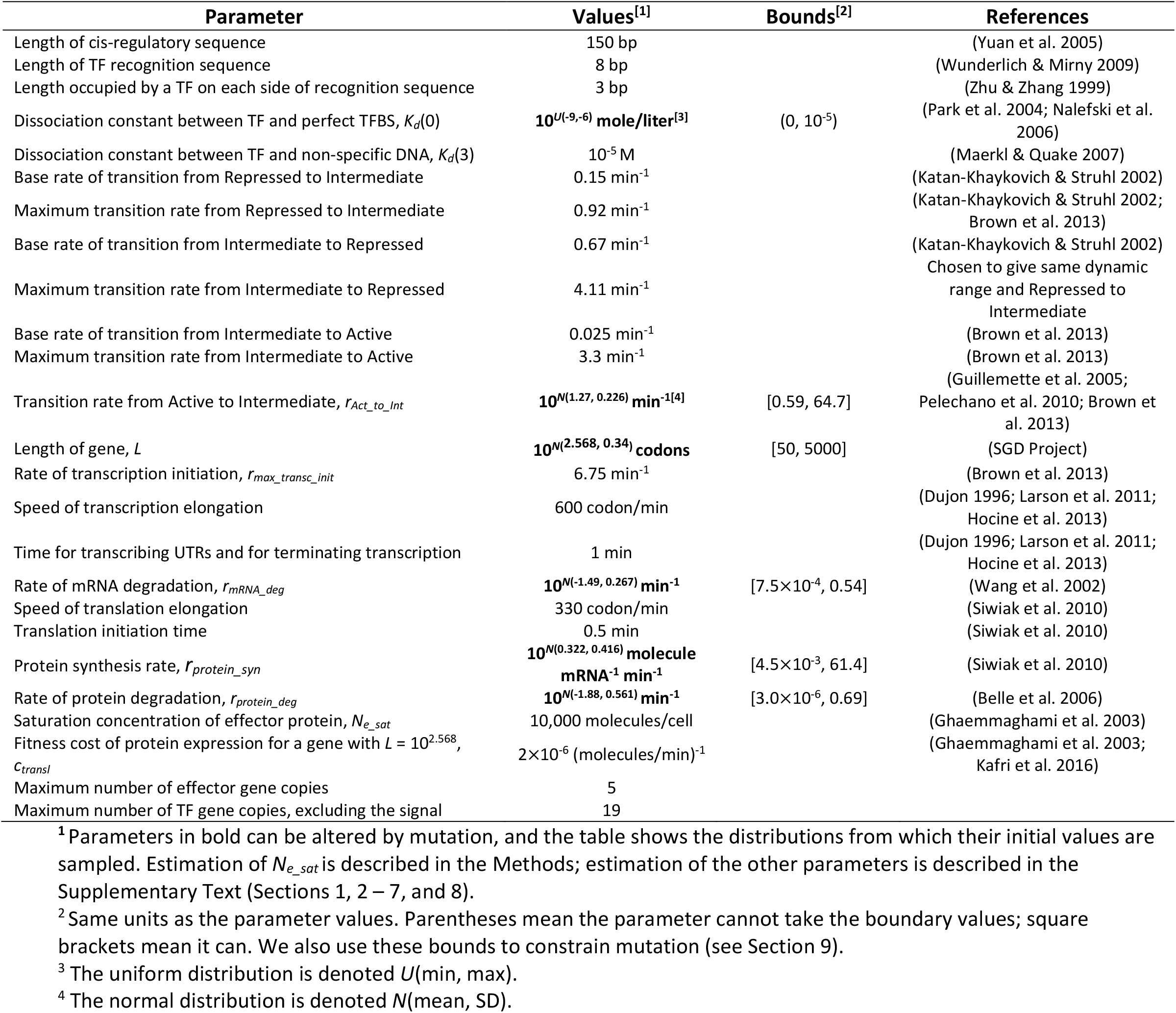
Major model parameters

**Table S2.**
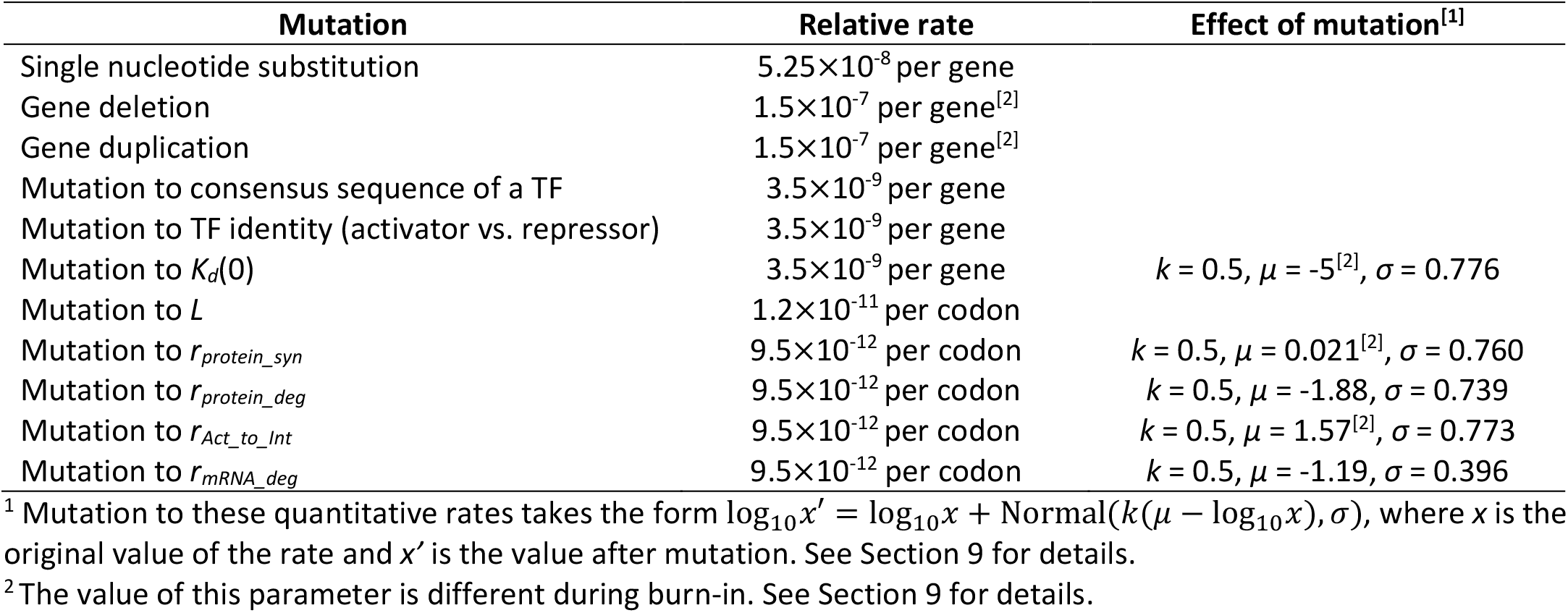
Mutation rates and effect sizes

**Table S3.**
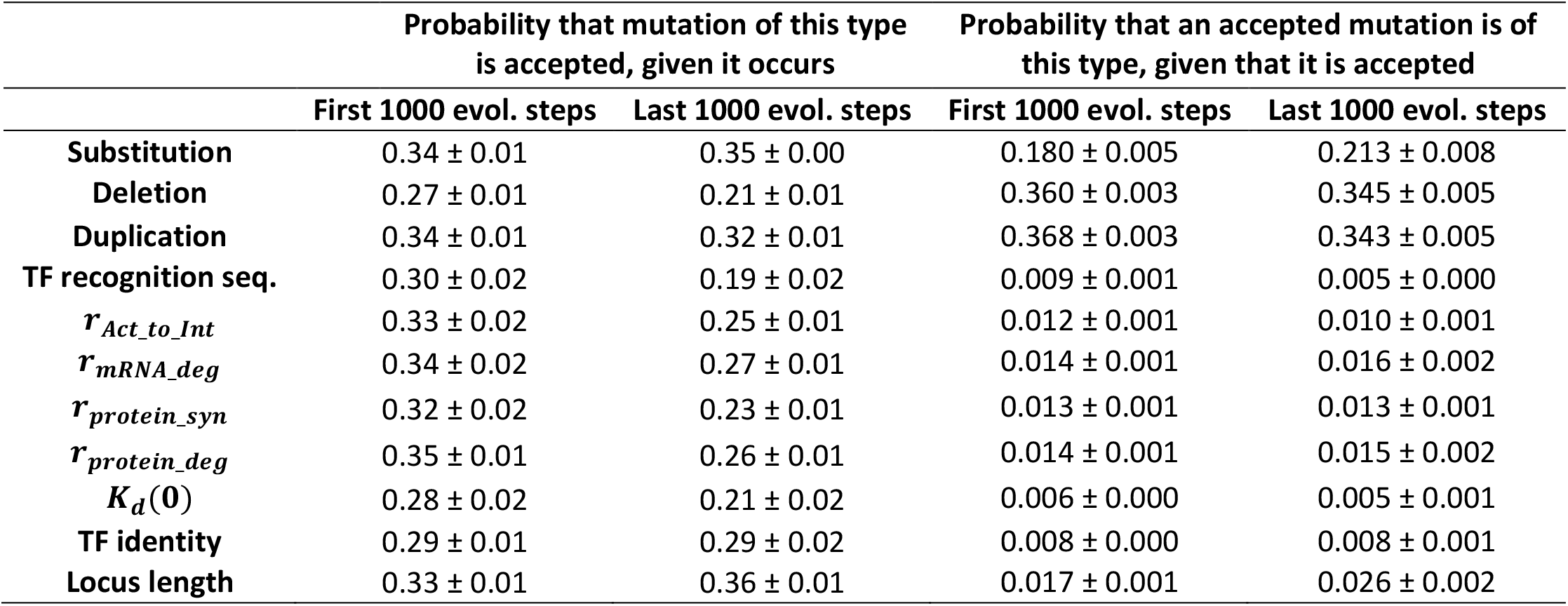
Summary of mutations that replaced the resident genotype. Data is shown as mean ± SE over the 45 evolutionary replicates under selection for filtering out a spurious signal, with the signal allowed to regulate the effector directly. Without selection, each mutation would have probability 50% of replacing the resident; selection reduces this to around one in three at the beginning of the simulation, down to around one in four at the end. This high rate of accepting mutations after fitness has plateaued suggests significant nearly neutral evolution, i.e. that slightly deleterious mutations fix and are then compensated for. The estimated selection coefficient need only be 10^-8^ for a mutant to replace the resident, which can be easily occur for a slightly deleterious mutation through the error in fitness estimation (see Evolution Simulation in the main text). Single nucleotide substitutions are particularly prone to nearly neutral evolution, whereas changes to the consensus sequence recognized by a TF are under stronger stabilizing selection. Deletion and duplication mutations are the most common forms of substitution not because they are more likely to be accepted, but because they occur at higher mutation rates.

**Table S4.**
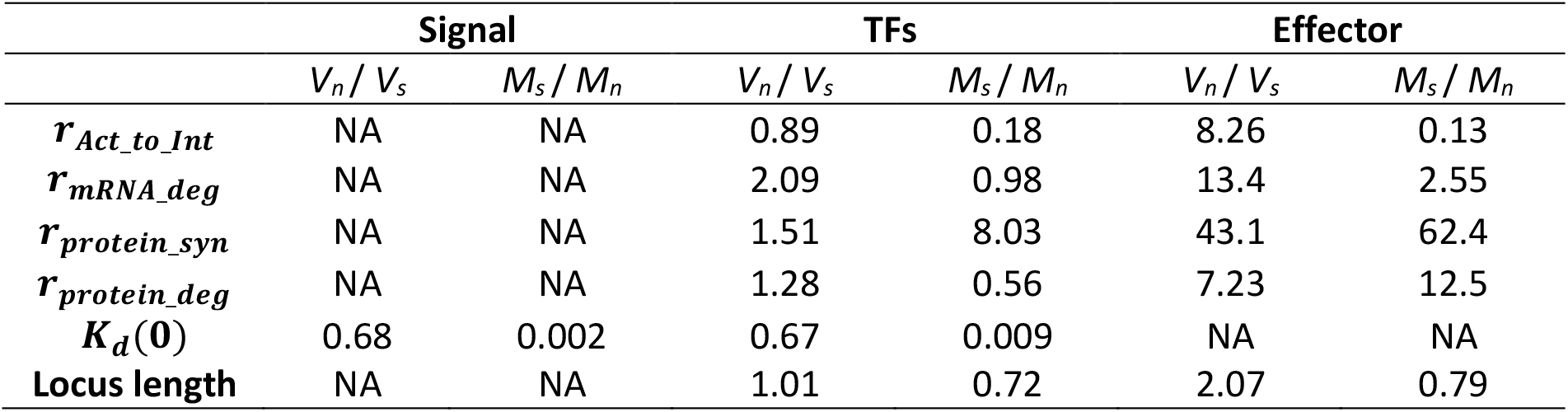
Evolutionary constraint on parameters in AND-gated C1-FFLs. Adaptive AND-gated C1-FFLs are taken from the 25 high-fitness replicates evolved for filtering out a spurious signal, where the signal directly regulates the effector. For each replicate, we sample one of the last 10,000 evolutionary time steps, and then sample one AND-gated C1-FFL in that genotype, should there be more than one (or resample a time step for that replicate, if there are none). We then take the variance *V*_*s*_ of each C1-FFL parameter value across the 25 replicates. We repeat this sampling process 100 times (using the same 25 replicates) and take the mean in order to obtain a better estimator of the variance in each parameter value. We compare this by a comparable variance *V*_*n*_ given no selection. We obtain these from 30 evolutionary replicates under no selection (from **Fig. 6**), sampling parameter values from the signal, from one TF gene copy, and from one effector gene, without the requirement for C1-FFL presence. Variances are calculated for log-transformed parameter values, except for locus length. For locus length, we use the coefficient of variation rather than variance, i.e. we divide both variances by the square of the average locus length. The table also shows the how the parameter values *M*_*s*_ in adaptive AND-gated C1-FFLs differ from the expected value *M*_*n*_ given no selection. *M*_*s*_ and *M*_*n*_ are calculated as arithmetic means for locus length and as geometric means for all other parameters. The variance ratio is greater than 1 (indicating constraint), for all parameters except *K*_*d*_(0), where the ratio of mean parameter values indicates that *K*_*d*_(0) is nevertheless subject to strong directional selection. Effectors are more constrained than TFs, likely because the former are less redundant, having evolved fewer gene copies (4.7 on average for effectors vs. 8.6 for TFs). High degradation rates of effector mRNA and protein suggest selection to shorten the impact of transient expression in response to a short spurious signal (**Fig. S2**). High degradation rates of effector mRNA and protein are also seen in **Tables S5** and **S6**.

**Table S5.**
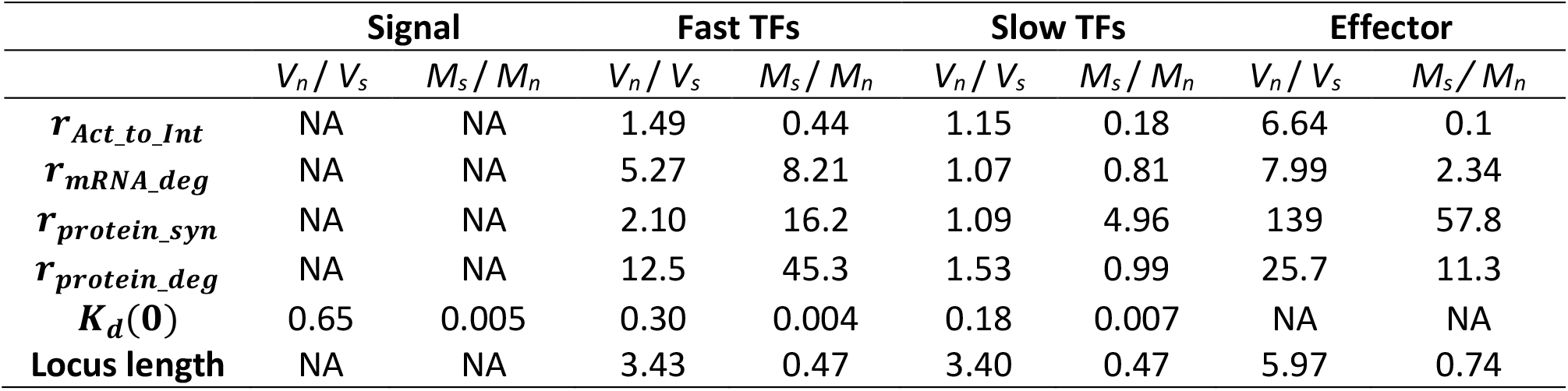
Evolutionary constraint on parameters in isolated AND-gated diamonds. *V*_*n*_, *V*_*s*_, *M*_*n*_, and *M*_*s*_ are defined in the same way as in **Table S4**, and are calculated from 18 high-fitness evolutionary replicates (**Fig. 7B**) in which isolated AND-gated diamonds occur in at least 100 of the last 10,000 evolutionary steps. Because they occur at low rates, we sample 50 times per evolutionary replicate, instead of 100 times as in **Tables S4** and **S6**. There is more constraint on fast TFs than on slow TFs. The fast TFs usually have more gene copies than the slow TFs, therefore redundancy is not the reason for this difference in constraint. As seen for the C1-FFLs in **Table S4**, effectors are more constrained than either TF, *K*_*d*_(0) shows strong selection for high affinity combined with high variance, and effectors evolve rapid degradation. Fast TFs exhibit not just fast protein degradation (which was used for their identification), but also fast mRNA degradation.

**Table S6.**
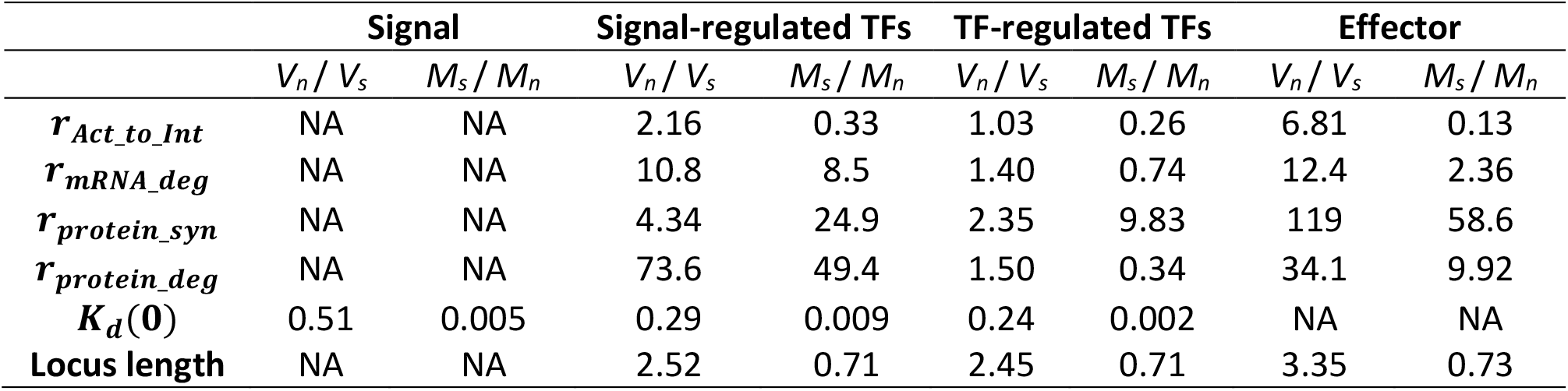
Evolutionary constraint on parameters in isolated AND-gated C1-FFLs. *V*_*n*_, *V*_*s*_, *M*_*n*_, and *M*_*s*_ are defined in the same way as in **Table S4**, and are calculated from 12 high-fitness evolutionary replicates (**Fig. 7B**) evolved when the signal cannot directly regulate the effector, and in which isolated AND-gated C1-FFLs occur in at least 1,000 out of the last 10,000 evolutionary steps. Note that the signal-regulated TFs, which are identified via network topology, also have high protein degradation rates, as is used to identify their fast TF counterparts in diamonds – they can thus be seen as a kind of fast TF. Consistent with results on C1-FFLs when direct regulation is allowed (**Table S4**) and results on isolated AND-gated diamonds (**Table S5**), effectors are more constrained than signal-regulated (fast) TFs, which are more constrained than TF-regulated (slow) TFs, despite an opposite trend in gene copy number. Note that selection promotes fast mRNA and protein degradation in fast TFs, but promotes slow degradation of slow TFs; this result is also found more weakly in **Table S5**.

