## Supplementary materials for "Feed-forward regulation adaptively evolves via dynamics rather than topology when there is intrinsic noise"

#### Table of contents


Supplementary Tables S1-S4 2

Supplementary Figures S1-S9 6

Supplementary Text Section 1: TF binding 17

Supplementary Text Section 2: TF occupancy 19

Supplementary Text Section 3: $r_{Act\_to\_Int}$ 21

Supplementary Text Section 4: Transcription delay times 22

Supplementary Text Section 5: Translation delay times and $r_{protein\_syn}$ 23

Supplementary Text Section 6: mRNA and protein decay rates 24

Supplementary Text Section 7: Simulation of gene expression 24

Supplementary Text Section 8: Cost of gene expression 29

Supplementary Text Section 9: Mutation 30

Supplementary Text Section 10: Burn-in evolutionary simulation conditions 34

Supplementary Text Section 11: Quantifying occurrence of network motifs 35

Supplementary Text Section 12: Perturbing network motifs 36

References 38

22 **Table S1. Major model parameters**

| Parameter | Values[1] | Bounds[2] | References |
|---|---|---|---|
| Length of cis-regulatory sequence | 150 bp | | (Yuan et al. 2005) |
| Length of TF recognition sequence | 8 bp | | (Wunderlich & Mirny 2009) |
| Length occupied by a TF on each side of recognition sequence | 3 bp | | (Zhu & Zhang 1999) |
| Dissociation constant between TF and perfect TFBS, $K_d(0)$ | **$10^{U(-9,-6)}$ mole/liter[3]** | $(0, 10^{-5})$ | (Park et al. 2004; Nalefski et al. 2006) |
| Dissociation constant between TF and non-specific DNA, $K_d(3)$ | $10^{-5}$ M | | (Maerkl & Quake 2007) |
| Base rate of transition from Repressed to Intermediate | 0.15 $min^{-1}$ | | (Katan-Khaykovich & Struhl 2002) |
| Maximum transition rate from Repressed to Intermediate | 0.92 $min^{-1}$ | | (Katan-Khaykovich & Struhl 2002; Brown et al. 2013) |
| Base rate of transition from Intermediate to Repressed | 0.67 $min^{-1}$ | | (Katan-Khaykovich & Struhl 2002) |
| Maximum transition rate from Intermediate to Repressed | 4.11 $min^{-1}$ | | Chosen to give same dynamic range and Repressed to Intermediate |
| Base rate of transition from Intermediate to Active | 0.025 $min^{-1}$ | | (Brown et al. 2013) |
| Maximum transition rate from Intermediate to Active | 3.3 $min^{-1}$ | | (Brown et al. 2013) |
| Transition rate from Active to Intermediate, $r_{Act\_to\_Int}$ | **$10^{N(1.27,\ 0.226)}$ $min^{-1}$[4]** | [0.59, 64.7] | (Guillemette et al. 2005; Pelechano et al. 2010; Brown et al. 2013) |
| Length of gene, $L$ | **$10^{N(2.568,\ 0.34)}$ codons** | [50, 5000] | (SGD Project) |
| Rate of transcription initiation, $r_{max\_transc\_init}$ | 6.75 $min^{-1}$ | | (Brown et al. 2013) |
| Speed of transcription elongation | 600 codon/min | | (Dujon 1996; Larson et al. 2011; Hocine et al. 2013) |
| Time for transcribing UTRs and for terminating transcription | 1 min | | (Dujon 1996; Larson et al. 2011; Hocine et al. 2013) |
| Rate of mRNA degradation, $r_{mRNA\_deg}$ | **$10^{N(-1.49,\ 0.267)}$ $min^{-1}$** | $[7.5\times10^{-4}, 0.54]$ | (Wang et al. 2002) |
| Speed of translation elongation | 330 codon/min | | (Siwiak et al. 2010) |
| Translation initiation time | 0.5 min | | (Siwiak et al. 2010) |
| Protein synthesis rate, $r_{protein\_syn}$ | **$10^{N(0.322,\ 0.416)}$ molecule $mRNA^{-1}$ $min^{-1}$** | $[4.5\times10^{-3}, 61.4]$ | (Siwiak et al. 2010) |
| Rate of protein degradation, $r_{protein\_deg}$ | **$10^{N(-1.88,\ 0.561)}$ $min^{-1}$** | $[3.0\times10^{-6}, 0.69]$ | (Belle et al. 2006) |
| Saturation concentration of effector protein, $N_{e\_sat}$ | 10,000 molecules/cell | | (Ghaemmaghami et al. 2003) |
| Fitness cost of protein expression for a gene with $L = 10^{2.568}$, $c_{transl}$ | $2\times10^{-6}$ $(molecules/min)^{-1}$ | | (Ghaemmaghami et al. 2003; Kafri et al. 2016) |
| Maximum number of effector gene copies | 5 | | |
| Maximum number of TF gene copies, excluding the signal | 19 | | |

[1] Parameters in bold can be altered by mutation, and the table shows the distributions from which their initial values are sampled. Estimation of $N_{e\_sat}$ is described in the Methods; estimation of the other parameters is described in the Supplementary Text (Sections 1, 2 – 7, and 8).

[2] Same units as the parameter values. Parentheses mean the parameter cannot take the boundary values; square brackets mean it can. We also use these bounds to constrain mutation (see Section 9).

[3] The uniform distribution is denoted $U$(min, max).

[4] The normal distribution is denoted $N$(mean, SD).

**Table S2. Mutation rates and effect sizes**

| Mutation | Relative rate | Effect of mutation[1] |
|---|---|---|
| Single nucleotide substitution | $5.25\times10^{-8}$ per gene | |
| Gene deletion | $1.5\times10^{-7}$ per gene[2] | |
| Gene duplication | $1.5\times10^{-7}$ per gene[2] | |
| Mutation to consensus sequence of a TF | $3.5\times10^{-9}$ per gene | |
| Mutation to TF identity (activator vs. repressor) | $3.5\times10^{-9}$ per gene | |
| Mutation to $K_d(0)$ | $3.5\times10^{-9}$ per gene | $k$ = 0.5, $\mu$ = -5[2], $\sigma$ = 0.776 |
| Mutation to $L$ | $1.2\times10^{-11}$ per codon | |
| Mutation to $r_{protein\_syn}$ | $9.5\times10^{-12}$ per codon | $k$ = 0.5, $\mu$ = 0.021[2], $\sigma$ = 0.760 |
| Mutation to $r_{protein\_deg}$ | $9.5\times10^{-12}$ per codon | $k$ = 0.5, $\mu$ = -1.88, $\sigma$ = 0.739 |
| Mutation to $r_{Act\_to\_Int}$ | $9.5\times10^{-12}$ per codon | $k$ = 0.5, $\mu$ = 1.57[2], $\sigma$ = 0.773 |
| Mutation to $r_{mRNA\_deg}$ | $9.5\times10^{-12}$ per codon | $k$ = 0.5, $\mu$ = -1.19, $\sigma$ = 0.396 |

[1] Mutation to these quantitative rates takes the form $\log_{10}x' = \log_{10}x + \mathrm{Normal}(k(\mu - \log_{10}x), \sigma)$, where $x$ is the original value of the rate and $x'$ is the value after mutation. See Section 9 for details.

[2] The value of this parameter is different during burn-in. See Section 9 for details.

| | Probability that mutation of this type is accepted, given it occurs | | Probability that an accepted mutation is of this type, given that it is accepted | |
|---|---|---|---|---|
| | **First 1000 evol. steps** | **Last 1000 evol. steps** | **First 1000 evol. steps** | **Last 1000 evol. steps** |
| **Substitution** | 0.34 ± 0.01 | 0.35 ± 0.00 | 0.180 ± 0.005 | 0.213 ± 0.008 |
| **Deletion** | 0.27 ± 0.01 | 0.21 ± 0.01 | 0.360 ± 0.003 | 0.345 ± 0.005 |
| **Duplication** | 0.34 ± 0.01 | 0.32 ± 0.01 | 0.368 ± 0.003 | 0.343 ± 0.005 |
| **TF recognition seq.** | 0.30 ± 0.02 | 0.19 ± 0.02 | 0.009 ± 0.001 | 0.005 ± 0.000 |
| $\boldsymbol{r_{Act\_to\_Int}}$ | 0.33 ± 0.02 | 0.25 ± 0.01 | 0.012 ± 0.001 | 0.010 ± 0.001 |
| $\boldsymbol{r_{mRNA\_deg}}$ | 0.34 ± 0.02 | 0.27 ± 0.01 | 0.014 ± 0.001 | 0.016 ± 0.002 |
| $\boldsymbol{r_{protein\_syn}}$ | 0.32 ± 0.02 | 0.23 ± 0.01 | 0.013 ± 0.001 | 0.013 ± 0.001 |
| $\boldsymbol{r_{protein\_deg}}$ | 0.35 ± 0.01 | 0.26 ± 0.01 | 0.014 ± 0.001 | 0.015 ± 0.002 |
| $\boldsymbol{K_d(0)}$ | 0.28 ± 0.02 | 0.21 ± 0.02 | 0.006 ± 0.000 | 0.005 ± 0.001 |
| **TF identity** | 0.29 ± 0.01 | 0.29 ± 0.02 | 0.008 ± 0.000 | 0.008 ± 0.001 |
| **Locus length** | 0.33 ± 0.01 | 0.36 ± 0.01 | 0.017 ± 0.001 | 0.026 ± 0.002 |

**Table S3. Summary of mutations that replaced the resident genotype.** Data is shown as mean $\pm$ SE over the 45 evolutionary replicates under selection for filtering out a spurious signal, with the signal allowed to regulate the effector directly. Without selection, each mutation would have probability 50% of replacing the resident; selection reduces this to around one in three at the beginning of the simulation, down to around one in four at the end. This high rate of accepting mutations after fitness has plateaued suggests significant nearly neutral evolution, i.e. that slightly deleterious mutations fix and are then compensated for. The estimated selection coefficient need only be $10^{-8}$ for a mutant to replace the resident, which can be easily occur for a slightly deleterious mutation through the error in fitness estimation (see Evolution Simulation in the main text). Single nucleotide substitutions are particularly prone to nearly neutral evolution, whereas changes to the consensus sequence recognized by a TF are under stronger stabilizing selection. Deletion and duplication mutations are the most common forms of substitution not because they are more likely to be accepted, but because they occur at higher mutation rates.

| | Signal | | TFs | | Effector | |
|---|---|---|---|---|---|---|
| | $V_n$ / $V_s$ | $M_s$ / $M_n$ | $V_n$ / $V_s$ | $M_s$ / $M_n$ | $V_n$ / $V_s$ | $M_s$ / $M_n$ |
| $r_{Act\_to\_Int}$ | NA | NA | 0.89 | 0.18 | 8.26 | 0.13 |
| $r_{mRNA\_deg}$ | NA | NA | 2.09 | 0.98 | 13.4 | 2.55 |
| $r_{protein\_syn}$ | NA | NA | 1.51 | 8.03 | 43.1 | 62.4 |
| $r_{protein\_deg}$ | NA | NA | 1.28 | 0.56 | 7.23 | 12.5 |
| $K_d(0)$ | 0.68 | 0.002 | 0.67 | 0.009 | NA | NA |
| **Locus length** | NA | NA | 1.01 | 0.72 | 2.07 | 0.79 |

**Table S4. Evolutionary constraint on parameters in AND-gated C1-FFLs.** Adaptive AND-gated C1-FFLs are taken from the 25 high-fitness replicates evolved for filtering out a spurious signal, where the signal directly regulates the effector. For each replicate, we sample one of the last 10,000 evolutionary time steps, and then sample one AND-gated C1-FFL in that genotype, should there be more than one (or resample a time step for that replicate, if there are none). We then take the variance $V_s$ of each C1-FFL parameter value across the 25 replicates. We repeat this sampling process 100 times (using the same 25 replicates) and take the mean in order to obtain a better estimator of the variance in each parameter value. We compare this by a comparable variance $V_n$ given no selection. We obtain these from 30 evolutionary replicates under no selection (from **Fig. 6**), sampling parameter values from the signal, from one TF gene copy, and from one effector gene, without the requirement for C1-FFL presence. Variances are calculated for log-transformed parameter values, except for locus length. For locus length, we use the coefficient of variation rather than variance, i.e. we divide both variances by the square of the average locus length. The table also shows the how the parameter values $M_s$ in adaptive AND-gated C1-FFLs differ from the expected value $M_n$ given no selection. $M_s$ and $M_n$ are calculated as arithmetic means for locus length and as geometric means for all other parameters. The variance ratio is greater than 1 (indicating constraint), for all parameters except $K_d(0)$, where the ratio of mean parameter values indicates that $K_d(0)$ is nevertheless subject to strong directional selection. Effectors are more constrained than TFs, likely because the former are less redundant, having evolved fewer gene copies (4.7 on average for effectors vs. 8.6 for TFs). High degradation rates of effector mRNA and protein suggest selection to shorten the impact of transient expression in response to a short spurious signal (**Fig. S2**). High degradation rates of effector mRNA and protein are also seen in **Tables S5** and **S6**.

| | Signal | | Fast TFs | | Slow TFs | | Effector | |
|---|---|---|---|---|---|---|---|---|
| | $V_n$ / $V_s$ | $M_s$ / $M_n$ | $V_n$ / $V_s$ | $M_s$ / $M_n$ | $V_n$ / $V_s$ | $M_s$ / $M_n$ | $V_n$ / $V_s$ | $M_s$ / $M_n$ |
| $\boldsymbol{r_{Act\_to\_Int}}$ | NA | NA | 1.49 | 0.44 | 1.15 | 0.18 | 6.64 | 0.1 |
| $\boldsymbol{r_{mRNA\_deg}}$ | NA | NA | 5.27 | 8.21 | 1.07 | 0.81 | 7.99 | 2.34 |
| $\boldsymbol{r_{protein\_syn}}$ | NA | NA | 2.10 | 16.2 | 1.09 | 4.96 | 139 | 57.8 |
| $\boldsymbol{r_{protein\_deg}}$ | NA | NA | 12.5 | 45.3 | 1.53 | 0.99 | 25.7 | 11.3 |
| $\boldsymbol{K_d(0)}$ | 0.65 | 0.005 | 0.30 | 0.004 | 0.18 | 0.007 | NA | NA |
| **Locus length** | NA | NA | 3.43 | 0.47 | 3.40 | 0.47 | 5.97 | 0.74 |

**Table S5. Evolutionary constraint on parameters in isolated AND-gated diamonds.** $V_n$, $V_s$, $M_n$, and $M_s$ are defined in the same way as in **Table S4**, and are calculated from 18 high-fitness evolutionary replicates (**Fig. 7B**) in which isolated AND-gated diamonds occur in at least 100 of the last 10,000 evolutionary steps. Because they occur at low rates, we sample 50 times per evolutionary replicate, instead of 100 times as in **Tables S4** and **S6**. There is more constraint on fast TFs than on slow TFs. The fast TFs usually have more gene copies than the slow TFs, therefore redundancy is not the reason for this difference in constraint. As seen for the C1-FFLs in **Table S4**, effectors are more constrained than either TF, $K_d(0)$ shows strong selection for high affinity combined with high variance, and effectors evolve rapid degradation. Fast TFs exhibit not just fast protein degradation (which was used for their identification), but also fast mRNA degradation.

| | Signal | | Signal-regulated TFs | | TF-regulated TFs | | Effector | |
|---|---|---|---|---|---|---|---|---|
| | $V_n$ / $V_s$ | $M_s$ / $M_n$ | $V_n$ / $V_s$ | $M_s$ / $M_n$ | $V_n$ / $V_s$ | $M_s$ / $M_n$ | $V_n$ / $V_s$ | $M_s$ / $M_n$ |
| $r_{Act\_to\_Int}$ | NA | NA | 2.16 | 0.33 | 1.03 | 0.26 | 6.81 | 0.13 |
| $r_{mRNA\_deg}$ | NA | NA | 10.8 | 8.5 | 1.40 | 0.74 | 12.4 | 2.36 |
| $r_{protein\_syn}$ | NA | NA | 4.34 | 24.9 | 2.35 | 9.83 | 119 | 58.6 |
| $r_{protein\_deg}$ | NA | NA | 73.6 | 49.4 | 1.50 | 0.34 | 34.1 | 9.92 |
| $K_d(0)$ | 0.51 | 0.005 | 0.29 | 0.009 | 0.24 | 0.002 | NA | NA |
| **Locus length** | NA | NA | 2.52 | 0.71 | 2.45 | 0.71 | 3.35 | 0.73 |

**Table S6. Evolutionary constraint on parameters in isolated AND-gated C1-FFLs.** $V_n$, $V_s$, $M_n$, and $M_s$ are defined in the same way as in **Table S4**, and are calculated from 12 high-fitness evolutionary replicates (**Fig. 7B**) evolved when the signal cannot directly regulate the effector, and in which isolated AND-gated C1-FFLs occur in at least 1,000 out of the last 10,000 evolutionary steps. Note that the signal-regulated TFs, which are identified via network topology, also have high protein degradation rates, as is used to identify their fast TF counterparts in diamonds – they can thus be seen as a kind of fast TF. Consistent with results on C1-FFLs when direct regulation is allowed (**Table S4**) and results on isolated AND-gated diamonds (**Table S5**), effectors are more constrained than signal-regulated (fast) TFs, which are more constrained than TF-regulated (slow) TFs, despite an opposite trend in gene copy number. Note that selection promotes fast mRNA and protein degradation in fast TFs, but promotes slow degradation of slow TFs; this result is also found more weakly in **Table S5**.

#### Supplementary Figures

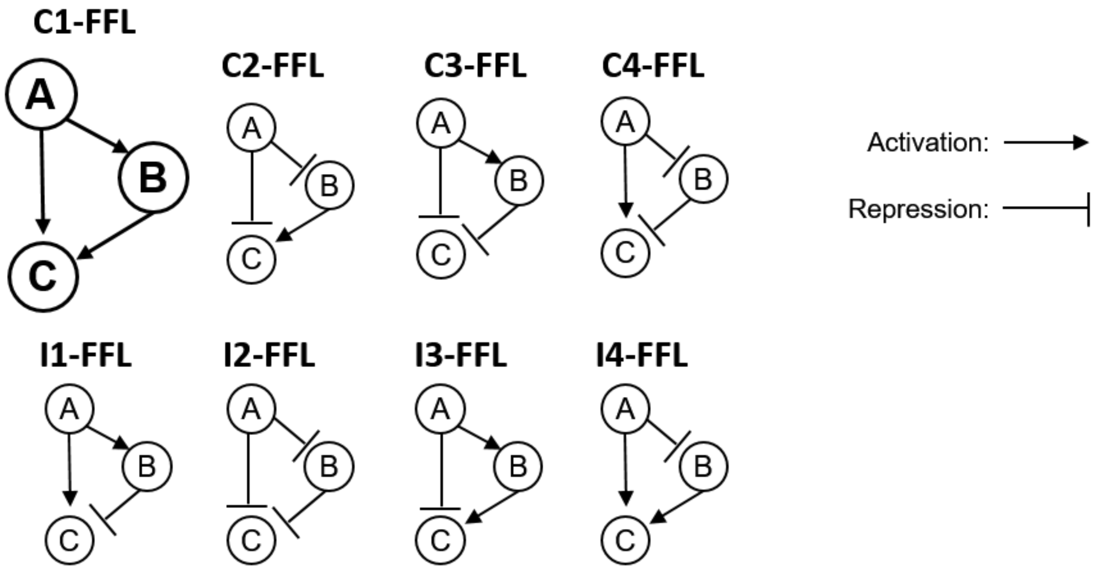


**Fig. S1. Feed-forward loops come in eight subtypes.** TF A and TF B can activate (indicated by arrows) or repress (indicated by bars) expression of the effector C as well as other TFs. Auto-regulation is allowed, but not shown. Following Milo et al. (2002), we exclude the case in which A and B regulate one another, rather than treating this case as two overlapping FFLs. C stands for coherent and I for incoherent.

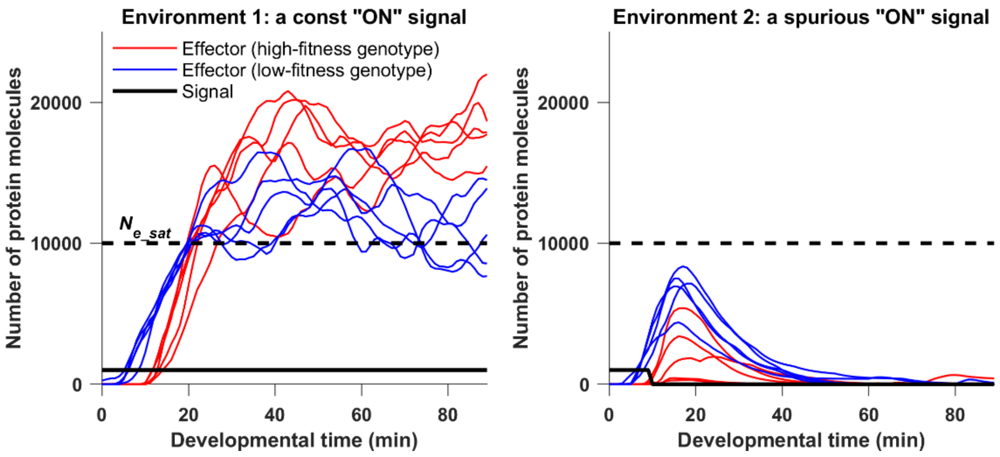


**Fig. S2 Examples of evolved phenotypes under selection for filtering out a short spurious signal.** The figure shows trajectories of the effector protein in one randomly chosen high-fitness replicate (red) and one randomly chosen low-fitness replicate (blue), as defined in **Fig. 4A**. The genotype of the final evolutionary step is used, and other genotypes were confirmed to behave similarly. Each genotype is illustrated by 5 replicate developmental simulations in each of the two environments. The high-fitness genotype has a longer delay followed by more rapid response given a consistent signal, with this longer delay reducing but not eliminating effector expression given a short spurious signal. The signal is allowed to directly regulate the effector in these simulations. The burn-in period is not shown, with developmental time zero corresponding to the moment the signal is turned on. Among developmental replicates of the same genotype, the concentration at a given time usually has an approximately log-normal distribution, but in environment 2 the distribution has two modes after the spurious signal turns off. One mode corresponds to expression at the basal rate, the other to a burst of expression that has yet to turn off. Because of this bimodality, we plot sample trajectories rather than mean concentration over many replicates.

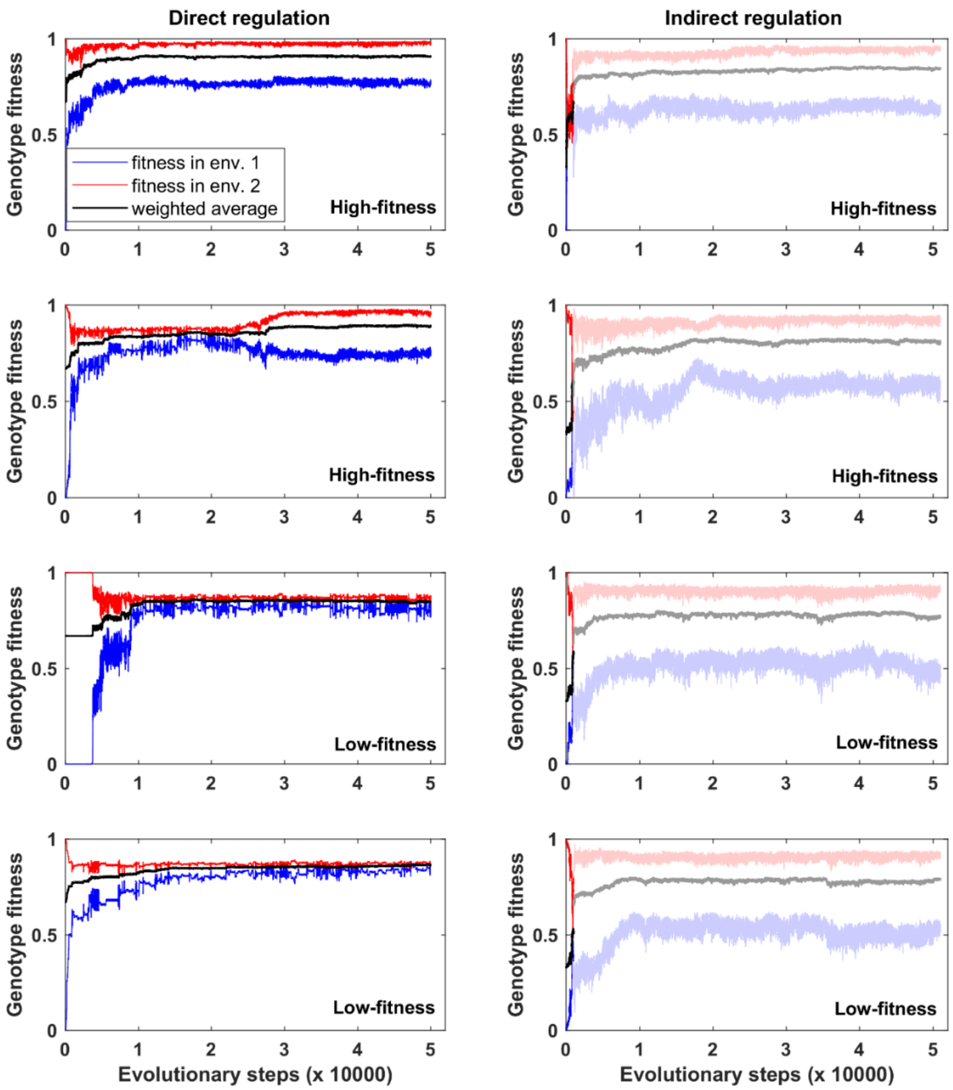


**Fig. S3 Representative fitness trajectories under selection to filter out short spurious signals.** Left panels: The signal is allowed to directly regulate the effector genes. Panels 1 and 3 correspond to the two genotypes shown in **Fig. S2**. Right panels: the signal cannot directly regulate the effector genes. Average fitness (black) is a weighted average of the blue and red trajectories, with environment 2 (where the signal is spurious) being considered twice as common as environment 1 (where the signal is sustained and real). When the signal cannot directly regulate the effector genes, evolutionary simulations begins with a burn-in phase that lasts 1000 evolutionary steps (see Evolutionary Simulation in the Main Text). We show the burn-in phase in undiluted color, and dilute color after burn-in. Most replicates quickly reach a stable fitness plateau (first and third rows). Certain replicates can be temporarily trapped at a low fitness plateau (second and third rows on the left).

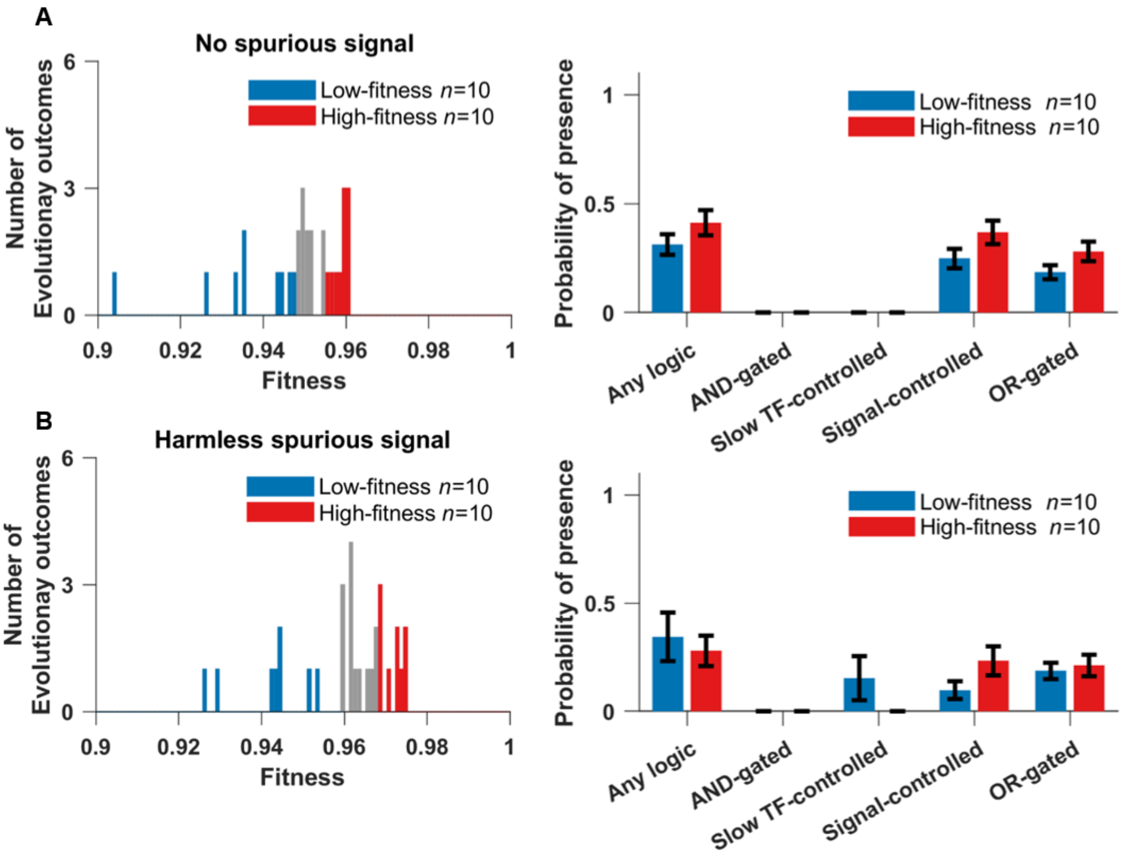


**Fig. S4 Genotypes evolved under control selective conditions: (A) "no spurious signal", and (B) "harmless spurious signal".** There is no clear evidence of a multimodal distribution of fitness outcomes among replicates (left), and C1-FFLs occur equally in the 10 genotypes of the highest fitness vs. the 10 genotypes of the lowest fitness (right), and so the entire distribution (left) was used to produce **Fig. 6**. Data are shown as mean±SE over evolutionary replicates.

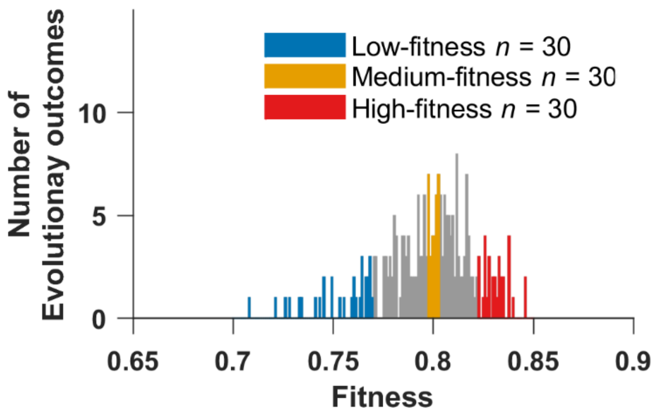


**Fig. S5 Fitness distribution of 258 evolutionary replicates under selection for filtering out short spurious signals, when the signal cannot directly regulate the effector.** The fitness of a replicate is the average genotype fitness over the last 10,000 evolutionary steps. Colors indicate replicates analyzed elsewhere.

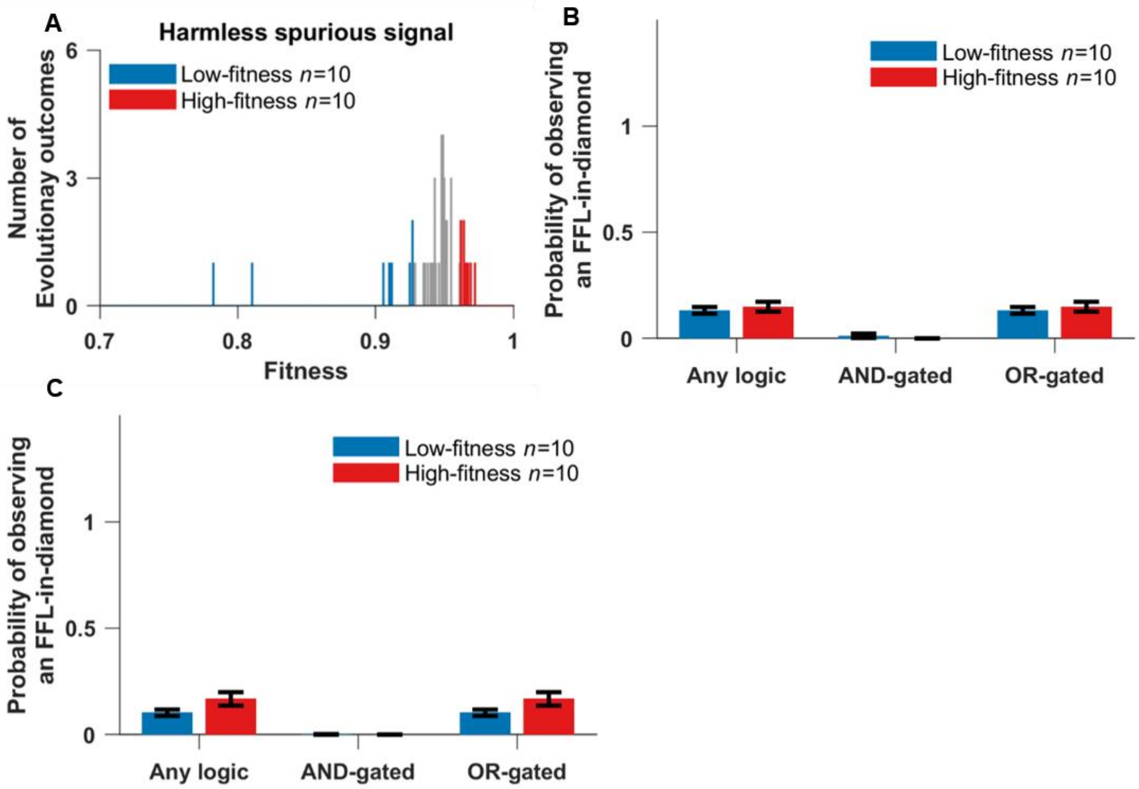


**Fig. S6 Evolution when responding to a spurious signal is harmless, when the signal is not allowed to directly regulate the effector. (A)** Fitness distribution of 50 replicate simulations. The occurrence of both **(B)** FFL-in-diamonds and **(C)** isolated diamonds were similar in the 10 genotypes with the highest fitness vs. in 10 genotypes with the lowest fitness. Weak (two-mismatch) TFBSs are included when scoring motifs. Data are shown as mean±SE over replicates. Isolated C1-FFLs rarely evolve under this condition, therefore their occurrence is not plotted.

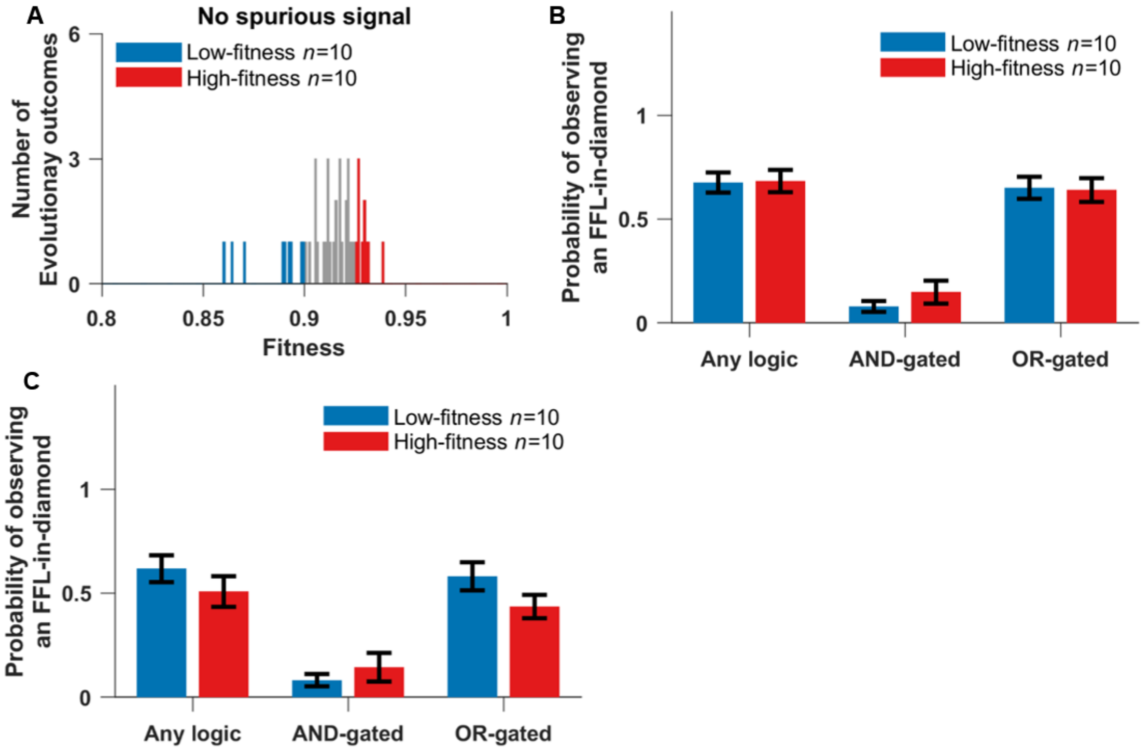


**Fig. S7 Evolution when there is no spurious signal, when the signal is not allowed to directly regulate the effector. (A)** Fitness distribution of 46 replicate simulations. The occurrence of both **(B)** FFL-in-diamonds and **(C)** isolated diamonds were similar in the 10 genotypes with the highest fitness vs. in the 10 genotypes with the lowest fitness. Weak (two-mismatch) TFBSs are included when scoring motifs. Data are shown as mean±SE over replicates. Isolated C1-FFLs rarely evolve under this condition, therefore their occurrence is not plotted.

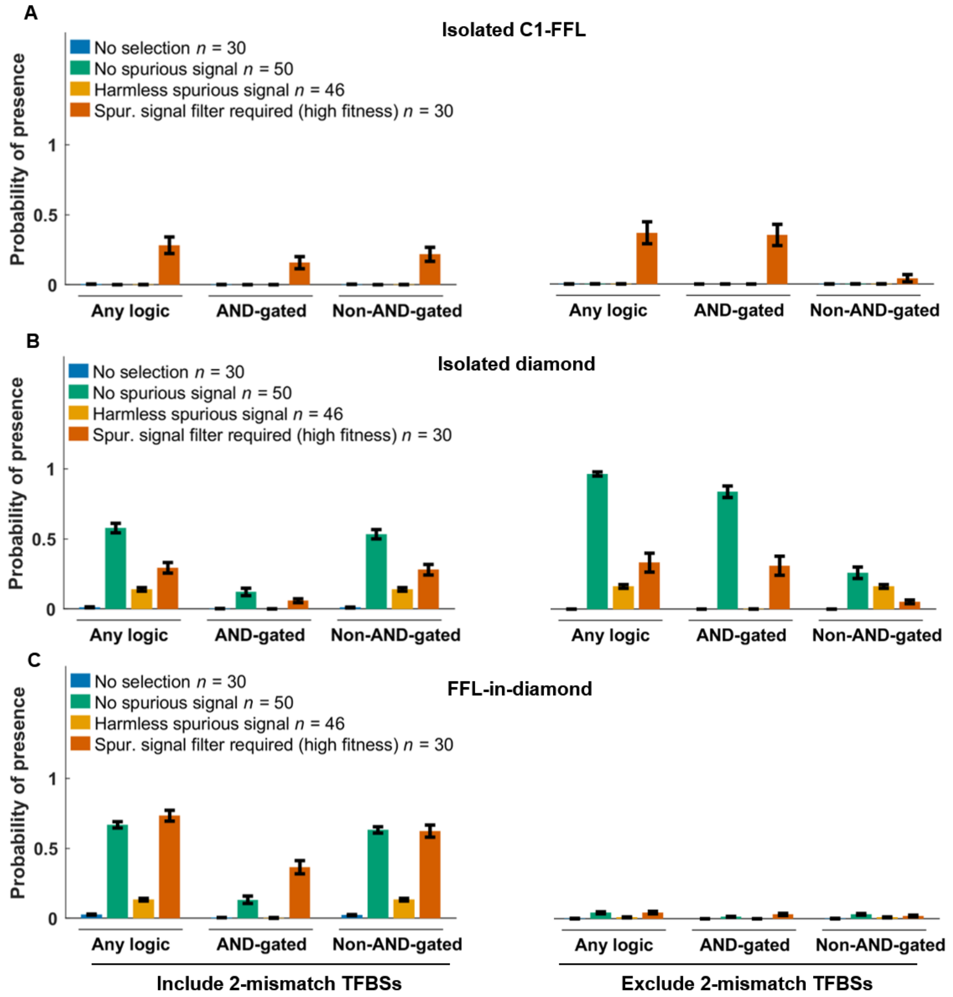


**Fig. S8 Selection for filtering out a short spurious signal is the primary way to evolve AND-gated C1-FFLs (A), but AND-gated isolated diamonds also evolve in the absence of spurious signals (B).** The signal is not allowed to directly regulate the effector, and the right panels of **(A)** and **(B)** are identical to **Fig. 10**. When scoring motifs, we either include (left) or exclude (right) all two-mismatch TFBSs in the cis-regulatory sequences of intermediate TF genes and effector genes. We excluded "no regulation" (**Fig. 2**) diamonds from the "Any logic" and "Non-AND-gated" tallies in **(B)**; this was necessary because of their high occurrence due to duplication and divergence of intermediate TFs. See Section 11 for the calculation of y-axis. Data are shown as mean±SE over evolutionary replicates.

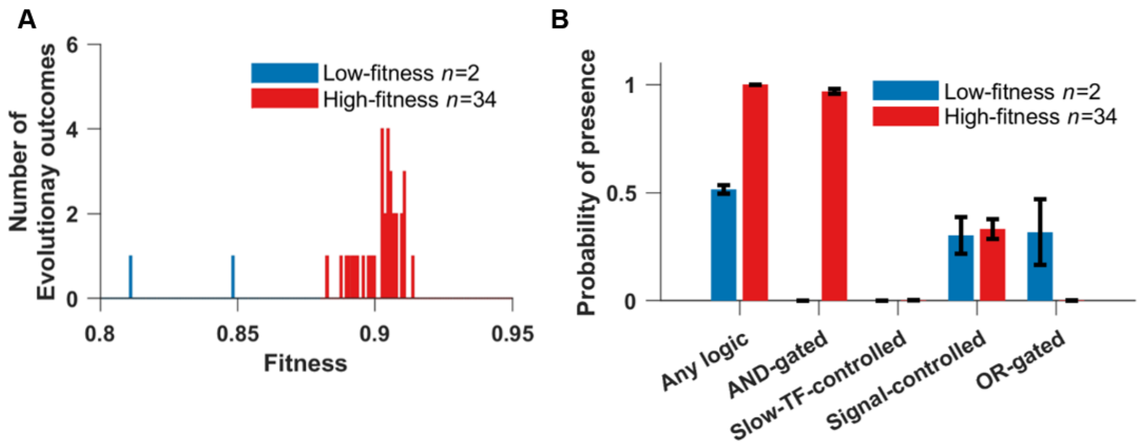


**Fig. S9 After removing cost of gene expression, AND-gated C1-FFLs are still associated with a successful response to selection for filtering out a short spurious signal.** The signal can directly regulate the effector genes. **(A)** We arbitrarily divide the 36 replicate simulations into high-fitness (red) and low-fitness (blue) groups. **(B)** The high-fitness replicates still evolve AND-gated C1-FFLs. Bars are mean±SE of the occurrence over replicate evolutionary simulations.

### Supporting Text

#### 1. TF binding

Transcription of each gene is controlled by TFBSs present within a 150-bp cis-regulatory region, corresponding to a typical yeast nucleosome-free region within a promoter (Yuan et al. 2005). The perfect TFBS for a typical yeast TF has information content equivalent to 13.8 bits (Wunderlich & Mirny 2009); this means that in a simplified model of binding where only one of the four nucleotides is a good match at each site, ~7 bp are recognized as an optimal consensus binding site. Maerkl & Quake (2007) reported that the TFBSs of two yeast TFs, Pho4p and Cbf1p, can have up to 2 mismatched sites within their 6 bp consensus binding sequence, while still binding the TF above background levels (Maerkl & Quake 2007). Our model therefore tracks TFBSs with up to 2 mismatches. This low information content implies a higher density of TFBSs within our cis-regulatory regions than our algorithm was able to handle, so we instead assigned each TF an 8-bp consensus sequence. Two TFs cannot simultaneously occupy overlapping stretches (Fig. S10), which we assume extend beyond the recognition sequence to occupy a total of 14 bp (Zhu & Zhang 1999); this captures competitive binding. The consequences of hindrance between TFBSs for the regulation of effector gene expression are shown in **Fig. 2**.

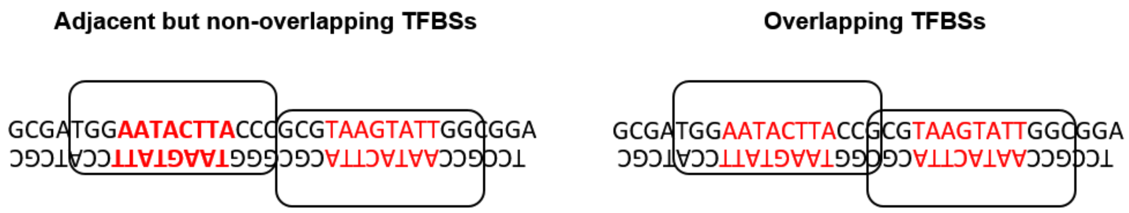


**Fig. S10** TFs (white boxes) recognize 8 bp (red) sites while occupying and thus excluding other TFs from a 14 bp long space. TFs are assumed to bind in either orientations (Sharon et al. 2012). The sequence on the left allows simultaneous binding but that on the right does not.

We denote the dissociation constant of a TFBS with $m$ mismatches as $K_d(m)$. Sites with $m$>3 mismatches are assumed to still bind at a background rate equal to $m$=3 mismatches, with dissociation constant $K_d(3) = 10^{-5}$ mole/liter (Maerkl & Quake 2007) for all TFs. We assume that each of the last three base pairs makes an equal and independent additive contribution $\Delta G_{bp} < 0$ to the binding energy (Benos et al. 2002): although not always true, this approximates average behavior well (Maerkl & Quake 2007). We ignore cooperativity in binding. Dissociation constants of eukaryotic TFs for perfect TFBSs can range from $10^{-5}$ mole/liter (Park et al. 2004) to $10^{-11}$ mole/liter (Nalefski et al. 2006). We initialize each TF with its own value of $\log_{10}(K_d(0))$ sampled from a uniform distribution between -6 and -9, with mutation capable of further expanding this range, subject to $K_d(0) < 10^{-5}$ mole/liter. Substituting $m$=0 and $m$=3 into

$$\Delta G_m = -RTlnK_d(m) = \Delta G_0 - \min(m, 3)\Delta G_{bp},$$

we can solve for $\Delta G_{bp}$ and $\Delta G_0$, and thus obtain $K_d(1)$ and $K_d(2)$ (the dissociation constants for TFBS with one and two mismatches, respectively).

Because TFs bind non-specifically to DNA at a high background rate, each nucleosome-free stretch of 14 bp can be considered to be a non-specific binding site (NSBS). A haploid *S. cerevisiae* genome is 12 Mb, 80% of which is wrapped in nucleosomes (Lee et al. 2007), yielding approximately $10^6$ potential non-specific binding sites (NSBSs). In a yeast nucleus of volume $3\times10^{-15}$ liters, the NSBS concentration is of order $10^{-4}$ mole/liter. To find the concentration of free TF [TF] in the nucleus given a total nucleic TF concentration of $C_{TF}$, we consider

$$K_d = \frac{[\text{binding\_site}][\text{TF}]}{[\text{binding\_site} \cdot \text{TF}]},$$

in the context of NSBSs, substitute [TF·NSBS] with $C_{TF}$ - [TF], and solve for

$$[\text{TF}] = \frac{K_d(3)}{K_d(3)+[\text{NSBS}]} C_{TF} = \frac{10^{-5}}{10^{-5}+10^{-4}} C_{TF} \approx 0.1 C_{TF}.$$

Thus, about 90% of total TFs are bound non-specifically, leaving about 10% free. The relatively small number of specific TFBSs is not enough to significantly perturb the proportion of free TFs, and so for the specific TFBSs with $m$<3 that are of interest in our model, we simply use $\widehat{K_d}(m)$ = $10K_d(m)$ to account for the reduction in the amount of available TF due to non-specific binding. We also convert $\widehat{K_d}$ from the units of mole/liter in which $K_d$ is estimated empirically to the more convenient molecules/nucleus. The rescaling factor $r$ for which $\widehat{K_d}$(in molecule/nucleus) = $r\widehat{K_d}$ (in mole/liter) is $3\times10^{-15}$ liter/nucleus $\times$ $6.02\times10^{23}$ molecule/mole = $1.8\times10^{9}$ molecule cell$^{-1}$ liter mole$^{-1}$. Taken together, $\widehat{K_d}$(molecule/nucleus) = $10rK_d$ (mole/liter), where the factor 10 accounts for non-specific TF binding.

#### 2. TF occupancy

Here we calculate the probability that there are $A$ activators and $R$ repressors bound to a given cis-regulatory region at a given moment in developmental time. First we note that if we consider TF $i$ binding to TFBS $j$ in isolation from all other TFs and TFBSs, Eq. S1 gives us probability of being bound:

$$P_b(j) = 1 - P_u(j) = \frac{C_i}{\widehat{K_d}+C_i}. \qquad \text{(S1)}$$

Let $P_{A,R}^{(n)}$ be a term proportional (for a given value of $n$) to the combined probability of all binding configurations in which exactly *A* activators and *R* repressors are bound to the first *n* binding sites along the cis-regulatory sequence. We calculate $P_{A,R}^{(n)}$ recursively, considering one additional TFBS at each step. Note that if two different TFs bind to exactly the same location on a cis-regulatory region, we treat this as two TFBSs, not as one, and treat first one and then the other in our recursive algorithm.

Consider the case where the (*n*+1)th binding site belongs to an activator. The case where this activator is not bound contributes $P_{A,R}^{(n)} P_u(n+1)$ to $P_{A,R}^{(n+1)}$. If it is bound, then we must also take into account that the (*n*+1)th binding site overlaps (partially or completely) with the last $H \geq 0$ sites, and so contributes $P_{A-1,R}^{(n-H)} P_b(n+1) \prod_{j=n-H+1}^{n} P_u(j)$. Taken together,

$$P_{A,R}^{(n+1)} = P_{A,R}^{(n)} P_u(n+1) + P_{A-1,R}^{(n-H)} P_b(n+1) \prod_{j=n-H+1}^{n} P_u(j).$$

Similarly, if the (*n*+1)th site belongs to a repressor, we have

$$P_{A,R}^{(n+1)} = P_{A,R}^{(n)} P_u(n+1) + P_{A,R-1}^{(n-H)} P_b(n+1) \prod_{j=n-H+1}^{n} P_u(j).$$

By definition, $P_{A,R}^{(n)} = 0$ for binding configurations that are impossible, e.g. those with negative *A* or negative *R*. We initialize the recursion at *n* = 0, where the only valid binding configuration is for *A* = *R* = 0, i.e. $P_{0,0}^{(0)} = 1$. At *n* = 1, $P_{0,0}^{(1)} \propto P_u(1)$, and if the binding site belongs to an activator, $P_{1,0}^{(1)} \propto P_b(1)$; otherwise, $P_{0,1}^{(1)} \propto P_b(1)$. For *N* = 1, the two probabilities sum to 1 and normalization is unnecessary. For higher values of $N = N_A + N_R$ TFBSs, we normalize $P_{A,R}^{(N)}$ at the end of the recursion by dividing by $\sum_{A=0}^{N_A}\sum_{R=0}^{N_R} P_{A,R}^{(N)}$ to get the probability of binding configurations that include exactly *A* activators and *R* repressors.

The total rate of all Gillespie events is

$$r_{total} = \sum_i^{Rep} r_{Rep\_to\_Int\_i} + \sum_i^{Int}(r_{Int\_to\_Rep\_i} + r_{Int\_to\_Act\_i}) + \sum_i^{Act}(r_{Act\_to\_Int\_i} + r_{transc}) + \sum_i^{genes} r_{mRNA\_deg\_i} N_{mRNA\_i},$$

where *Rep*, *Int*, and *Act* are the numbers of gene copies in our haploid model that are in the Repressed, Intermediate, and Active chromatin states, respectively, and $N_{mRNA\_i}$ is the number of completely transcribed mRNA molecules from gene *i*. We only simulate degradation of full transcribed mRNA, and not that of mRNA that are still being transcribed, because the latter are already captured implicitly by $r_{max\_transc\_init}$, which is based on mRNAs that complete transcription (Brown et al. 2013). Once an mRNA finishes transcription, it is subjected to degradation regardless of whether ribosome loading is complete.

The waiting time $\Delta t$ before the next Gillespie event is

$$\Delta t = \frac{x}{r_{total}},$$

where *x* is random number drawn from an exponential distribution with mean 1. Which Gillespie event takes place next is sampled only if a different update does not happen first. If a fixed event is scheduled to happen first at $\Delta t_1 < \Delta t$, we advance time by $\Delta t_1$, update the state of the cell, and calculate a new $r_{total}'$. Since the cellular activity has been going on with the old rate $r_{total}$ for $\Delta t_1$, the remaining "labor" required to trigger the Gillespie event planned earlier is reduced. The new waiting time, $\Delta t'$, to trigger the planned Gillespie event is

$$\Delta t' = \frac{x - r_{total}\Delta t_1}{r_{total}'}.$$

Gene duplication creates $n \geq 1$ genes producing the same protein, where each copy *i* might have diverged in their production rate $r_{protein\_syn\_i}$ and degradation rate $r_{protein\_deg\_i}$. Complete proteins are produced continuously once an mRNA molecule is fully loaded with ribosomes, which occurs 0.5 + *L* / 330 minutes after transcription is complete – the concentration of such molecules is denoted $N_{mRNA\_aft\_delay\_i}(t)$. Total protein concentration obeys:

$$N'_{protein}(t) = \sum_i^n \left( r_{protein\_syn\_i} N_{mRNA\_aft\_delay\_i}(t) - r_{protein\_deg\_i} N_{protein\_i}(t) \right). \qquad \text{(S3)}$$

Protein concentrations are updated using a closed-form integral of Eq. S3

$$N_{protein}(t_1) = \sum_i^n \left( \frac{r_{protein\_syn\_i} N_{mRNA\_aft\_delay\_i}}{r_{protein\_deg\_i}} + \left( N_{protein\_i}(t_0) - \frac{r_{protein\_syn\_i} N_{mRNA\_aft\_delay\_i}}{r_{protein\_deg\_i}} \right) e^{-r_{protein\_deg\_i}(t_1 - t_0)} \right) \qquad \text{(S4)}$$

with this expression updated every time a Gillespie or fixed event at time $t_1$ changes the value of $N_{mRNA\_aft\_delay\_i}$.

In between updates, values of $P_A$, $P_R$, $P_{A\_no\_R}$, and $P_{notA\_no\_R}$, and hence chromatin transition rates, are calculated under the approximation of constant $N_{protein}$. Additional updates, above and beyond fixed and Gillespie events, are performed in order to ensure that chromatin transition rates do not change too dramatically from one update to the next. We use a target of $D = 0.01$ for the amount of change tolerated in the values of $P_A$, $P_R$, $P_{A\_no\_R}$, and $P_{notA\_no\_R}$, in order to schedule updates after time $\Delta t^*$, which are triggered when neither a Gillespie event nor a fixed event occurs before this time has elapsed, i.e. when $\Delta t^* < \Delta t_1$ and $\Delta t^* < \Delta t$.

There is the greatest potential for large changes after an update that changes the value of $N_{mRNA\_aft\_delay\_i}$. In this case, we use Eq. S1 to solve for the time interval for which the probability that TF $i$ would be bound to a single perfect and non-overlapping TFBS would change by $D$, by choosing $\Delta t^* > 0$ that satisfies

$$\left|\frac{N_i(t)}{N_i(t)+K_{d,i}^*(0)} - \frac{N_i(t+\Delta t^*)}{N_i(t+\Delta t^*)+K_{d,i}^*(0)}\right| = D. \qquad \text{(S5)}$$

A solution for $\Delta t$ may not exist, e.g. if the concentration of TF $i$ is decreasing but $P_b(t_2) < D$. In such cases, we set $\Delta t^*$ to infinity.

When the previous update does not change any $N_{mRNA\_aft\_delay\_i}$ values, then we modify $\Delta t^*$ adaptively. Let $d$ be the maximum of $\Delta P_A$, $\Delta P_R$, $\Delta P_{A\_no\_R}$, and $\Delta P_{notA\_no\_R}$ during the last update. We then schedule an update at

$$\Delta t^{*\prime} = \frac{D}{d}\Delta t^{*}. \quad \text{(S6)}$$

After an update that changes the value of $N_{mRNA\_aft\_delay\_i}$, we use the smaller value from Eqs. S5 and S6. These additional update times are discarded and recalculated when a Gillespie or fixed event occurs first.

In **Fig. S11**, we see that simulations rarely exceed our target of $D$=0.01, and do so only modestly.

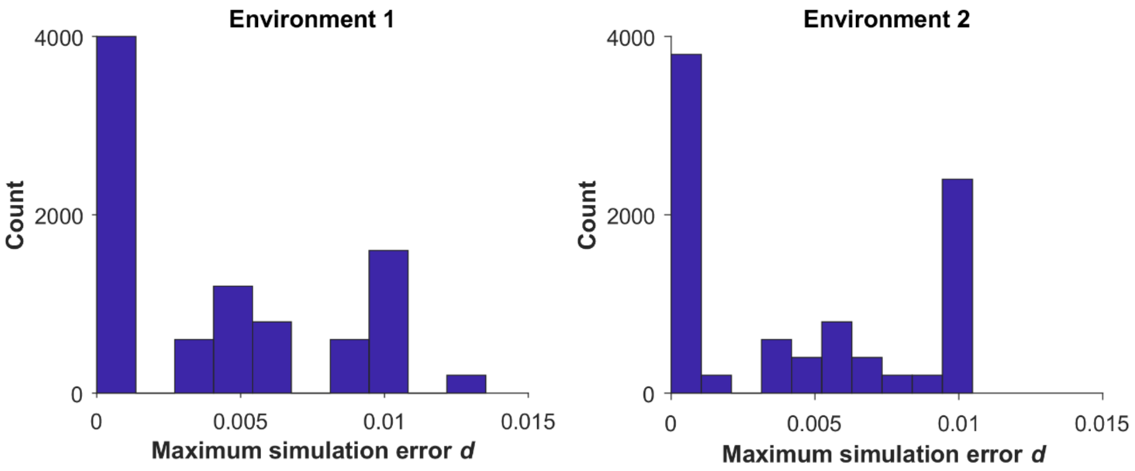


**Fig. S11 Our updating algorithm is able to limit simulation errors.** The distribution across 9,000 simulations of the maximum value of *d* over the course of development. For each of the 45 evolutionary replicates in **Fig. 4**, we run 200 simulations of development of the final evolved genotype. These genotypes were the outcome of evolution under selection for filtering out short spurious signals, in which direct regulation of the effector by the signal is not allowed. In environment 1 a genotype responds to a constant "ON" signal and in environment 2 it responds to a short spurious signal (**Fig. 3**).

The overall cost of gene expression at time *t*, *C*(*t*) is:

$$C(t) = c_{transl}\left(\sum_1^n \frac{L_i}{10^{2.568}} r_{transl\_init\_i} N_{mRNA\_aft\_delay\_i}(t) + \sum_1^n \frac{L_i}{10^{2.568}} \frac{r_{transl\_init\_i}}{2} N_{mRNA\_during\_delay\_i}(t)\right).$$

The second term represents transcripts that are on average half-loaded with ribosomes, and hence experiencing on average half the cost of translation. We integrate *C*(*t*) within segments of constant *C*(*t*) to obtain the overall cost of gene expression during a simulation.

#### 9. Mutation

Because we use an origin-fixation approach, only the relative and not the absolute values of our mutation rates matter. In *S. cerevisiae,* the rates of small indels and of single nucleotide substitutions have been estimated as $0.2\times10^{-10}$ per base pair and $3.3\times10^{-10}$ per base pair, respectively (Lynch et al. 2008). Thus, cis-regulatory sequences are primarily shaped by single nucleotide substitutions. We do not model small indels in the cis-regulatory sequence, but increase the single nucleotide substitution up to $3.5\times10^{-10}$ per base pair to compensate. This corresponds to a rate of $5.25\times10^{-8}$ per 150 bp cis-regulatory sequence.

Lynch et al. (2008) also report a rate of gene duplication of $1.5\times10^{-6}$ per gene and of deletion of $1.3\times10^{-6}$ per gene (not including non-deletion-based loss of function mutations). These values turned out to swamp the evolution of TFBSs and hence significantly slow down our simulations, so we chose values 10-fold lower, making both gene duplication and gene deletion occur at rate $1.5\times10^{-7}$ per gene. This preserves their numerical excess but reduces its magnitude.

The value of $\sigma$ controls mutational effect size. We set the value of $\sigma$ such that 1% of mutational changes from $x$=$10^{\mu}$ go beyond the boundary values, for simplicity approximating by considering only the closer of the two boundary values on a log scale, i.e. we solve Eq. S8 for $\sigma$:

$$\begin{cases} P(\mu + \text{Normal}(0,\sigma) \geq \log_{10}U) = 0.01, & \text{if the upper bound U is closer} \\ P(\mu + \text{Normal}(0,\sigma) \leq \log_{10}L) = 0.01, & \text{if the lower bound L is closer} \end{cases} \quad \text{(S8)}$$

For example, the upper and the lower bounds of $r_{mRNA\_deg}$ are 0.54 $\text{min}^{-1}$ and $7.5\times10^{-4}$ $\text{min}^{-1}$; on a log-scale, the upper bound is closer to $10^{\mu}$ = $10^{-1.19}$ $\text{min}^{-1}$. Plugging these values in Eq. S8 and solving for $\sigma$, we have $\sigma$ = 0.396. We set the values of $\sigma$ for $r_{protein\_syn}$, and $r_{protein\_deg}$ in the same way. However for $r_{Act\_to\_Int}$, $\sigma$ is set according to the lower bound, even though it is the more distant from $10^{\mu}$, because otherwise a stable preinitiation complex will evolve too rarely. Under this high mutational variance, evolutionary outcomes at the two bounds are still only observed 5% of the time. For $K_d(0)$, because its upper bound is equal to $10^{\mu}$, we set $\sigma$ to 0.776, such that 1% of mutations can change the values of $K_d(0)$ by 100-fold or more.

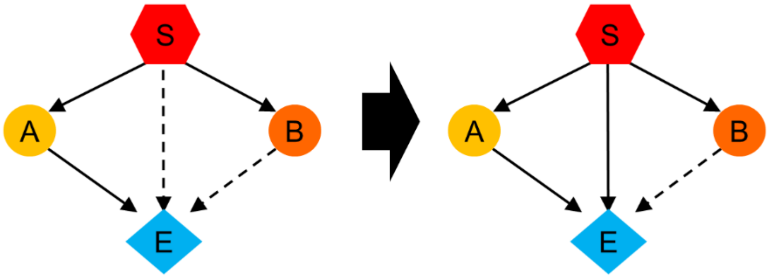


**Fig. S12 Examples of confounding motifs in perturbation analysis.** The TRN on the left contains a slow TF-controlled C1-FFL (S-B-E) and an AND-gated C1-FFL (S-C-E). To convert S-C-E into a signal-controlled C1-FFL, we need to add one TFBS for the signal to the cis-regulatory sequence of E. However, this change also makes S-B-E OR-gated, making it difficult to conclude whether it is the AND gate logic of S-B-E that matters for fitness.
